# CRISPR-GEM: A Novel Machine Learning Model for CRISPR Genetic Target Discovery and Evaluation

**DOI:** 10.1101/2024.07.01.601587

**Authors:** Josh P. Graham, Yu Zhang, Lifang He, Tomas Gonzalez-Fernandez

## Abstract

CRISPR gene editing strategies are shaping cell therapies through precise and tunable control over gene expression. However, achieving reliable therapeutic effects with improved safety and efficacy requires informed target gene selection. This depends on a thorough understanding of the involvement of target genes in gene regulatory networks (GRNs) that regulate cell phenotype and function. Machine learning models have been previously used for GRN reconstruction using RNA-seq data, but current techniques are limited to single cell types and focus mainly on transcription factors. This restriction overlooks many potential CRISPR target genes, such as those encoding extracellular matrix components, growth factors, and signaling molecules, thus limiting the applicability of these models for CRISPR strategies. To address these limitations, we have developed CRISPR-GEM, a multi-layer perceptron (MLP)-based synthetic GRN constructed to accurately predict the downstream effects of CRISPR gene editing. First, input and output nodes are identified as differentially expressed genes between defined experimental and target cell/tissue types respectively. Then, MLP training learns regulatory relationships in a black-box approach allowing accurate prediction of output gene expression using only input gene expression. Finally, CRISPR-mimetic perturbations are made to each input gene individually and the resulting model predictions are compared to those for the target group to score and assess each input gene as a CRISPR candidate. The top scoring genes provided by CRISPR-GEM therefore best modulate experimental group GRNs to motivate transcriptomic shifts towards a target group phenotype. This machine learning model is the first of its kind for predicting optimal CRISPR target genes and serves as a powerful tool for enhanced CRISPR strategies across a range of cell therapies.

## 1. Introduction

Clustered regularly interspaced short palindromic repeats (CRISPR) is a bioinspired gene editing tool for regulating gene expression at specific genomic locations^1^. Distinct CRISPR-associated (Cas) proteins are available for precise knock-in, knock-out, activation or inhibition of target genes, which must be carefully selected based on the CRISPR strategy and therapeutic application^2^. Target genes are often well-established genes or biomarkers in tissues or disease states due to their known effects on cell behavior. However, CRISPR-mediated alteration of single-gene expression can influence the expression of thousands of genes as governed by complex gene regulatory networks (GRNs). For example, activating *COL2A1*, a major component of cartilage extracellular matrix (ECM), using CRISPR activation (CRISPRa) in adipose-derived stem cells resulted in 2,532 differentially expressed genes (DEGs)^3^. Similarly, knocking-out *PRDM1* in CAR-T cells resulted in 2,192 DEGs which drove an early memory phenotype and dynamic cytokine secretion for enhanced anti-tumor activity^4^. These broad transcriptomic changes demonstrate that CRISPR interventions have cascading effects on larger GRNs, resulting in unexpected transcriptomic outcomes. Therefore, CRISPR-based cell therapies require informed target gene selection methods to best modulate GRNs for precise phenotypic regulation and enhanced therapeutic effects.

Mapping and reconstructing GRNs represents a promising strategy for identifying optimal CRISPR target genes. Initially, knock-out experiments were used to characterize the downstream transcriptomic effects of specific genes and generate large gene regulation databases. However, the complexity of the genome coupled with variation in experimental procedures made these methods challenging to interpret. More recently, machine learning (ML) have been successful in decoding gene regulatory relationships and elucidating important transcription factors and regulatory mechanisms. Early ML methods employed linear, tree-based, weighted correlation, RNA velocity, or enrichment analysis strategies to learn regulatory relationships from RNA-seq data^5–14^. Okawa *et al.* applied such principles to the prediction of candidate genes for stem cell differentiation using a Boolean model^15^. Still, these frameworks fall short in capturing the non-linear nature of gene regulation caused by feedback loops, competing interactions, and post-translational modifications. González-Blas *et al.* sought to partially address this by combining gene expression and transcription factor binding data to construct an enhancer-driven GRN, enabling the identification of *RUNX3* as an important catalyst of melanoma formation^16^. This strategy greatly improved the interpretability of GRNs but could not distinguish the effects of multiple transcription factors acting on one binding region. More complex models are therefore required to decode all non-linear relationships involved in gene expression for accurate and interpretable GRN construction. To address this, a surge of deep learning (DL) models have employed multilayer perceptrons (MLPs), graph neural networks (GNNs), recurrent neural networks (RNNs), or transformers to capture non-linear relationships in GRNs^17–21^. While DL has shown remarkable improvements in GRN accuracy, more strategic model design is required to best tune these models for evaluating CRISPR target genes.

DL models are highly influenced by the data source used for training. Many DL-based GRN models are trained using single cell RNA-seq data, which provides clear insights into genomic regulation without the conflicting signals from cellular subpopulations. However, bulk RNA-seq might better represent CRISPR-mediated transcriptomic shifts as they affect entire biological systems, comprised of diverse cell types and subpopulations. Magnusson *et al.* established effective GRN reconstruction using bulk RNA-seq data by training an MLP to predict expression throughout the entire genome^18^. Their model achieved over 95% accuracy using only the expression of 1600 known transcription factors as inputs^18^. This approach better forecasts how transcription factor expression drives cellular processes in a biologically relevant manner compared to single cell RNA-seq models. Additionally, the MLP structure accurately predicts complex and irregular gene regulatory relationships without the hyperparameter sensitivity or speculative assumptions required by simpler support vector regression (SVR) models or more complex GNNs. Further, the black-box framework allows for accurate prediction of gene expression while indirectly accounting for variables such as external signaling, ECM interactions, or post-translational protein modification. However, by limiting inputs to transcription factors, many potential CRISPR target genes can be discarded such as those encoding for ECM components, growth factors, and signaling molecules. Although these genes are regulated by transcription factors, they play important roles in cellular processes and can provide feedback that alters transcription factor expression. Well-informed CRISPR strategies must consider all genes to achieve the maximal therapeutic effect.

To address these limitations, we have developed CRISPR-GEM (Gene Evaluation through Machine Learning). This system involves a synthetic MLP-based GRN with input/output genes filtered to accommodate specific CRISPR applications. We then utilize this GRN to score the therapeutic impact of individual CRISPR target gene candidates to aid in the design of well-informed CRISPR strategies. CRISPR-GEM defines two groups: (1) the experimental group, consisting of the cell or tissue to be CRISPR edited, and (2) the target group, representing the desired therapeutic phenotype. These groups are compared to identify DEGs, which are filtered based on the CRISPR strategy (knock-in, knock-out, activation, or inhibition) to define input and output genes for the neural network. This tissue-specific MLP is trained on bulk RNA-seq data to learn the regulatory relationships between the identified genes, the broader genome and all regulatory mechanisms controlling the transcriptome. Finally, each gene in the experimental group RNA-seq datasets is perturbed in a CRISPR-mimetic manner and input into the trained MLP. The CRISPR-GEM scoring system compares the magnitude and direction of changes in gene expression between model predictions for the perturbed dataset and target group datasets to assess the overall therapeutic effect for each perturbed gene. As a proof of concept, we identify promising CRISPR target genes for 1) the differentiation of induced pluripotent stem cells (iPSCs) to regulatory T-cells (Tregs), 2) the chondrogenic differentiation of mesenchymal stem cells (MSCs), and 3) the reversal of osteoarthritis (OA) in cartilage tissues. Our results identify well-established genes *SRGN, PRG4, and FAP* among top scoring genes for the respective applications which validates model performance. Furthermore, CRISPR-GEM identifies novel gene targets and highlights the essential genes and cellular pathways necessary for specifically modulating cell phenotype to inspire advanced CRISPR strategies.

## 2. Methods & Procedures

### 2.1. Data Preparation

RNA-seq data for primary human cells and tissues collected using Illumina HiSeq 2000 or HiSeq 2500 platforms was identified using the gene expression omnibus (GEO) and can be found in Supp. Table 1. Data was downloaded from the sequencing read archive (SRA) using the SRA toolkit “prefetch” and “fasterq-dump” executables^22^. The raw fastq files were then aligned to the GRCh38.p14 genome assembly using the kallisto algorithm which yields sequencing data in transcript per million (tpm) or estimated counts^23,24^. The data for tpm or estimated counts was then sorted into separate data frames with samples in the rows and each ensemble ID in the columns. The column names were then mapped to their gene symbol and grouped per gene by taking the maximum value.

**Table 1.** Input & Output gene selection parameters.

| **CRISPR Strategy** | **Input Gene Selection Method** |  | **Output Gene Selection Method** |
| --- | --- | --- | --- |
| CRISPRko | $\log_2FC < 1$ | $p < 0.01$ | $p < 0.01$ |
| CRISPRki | $\log_2FC > 0.7$ | $p < 0.01$ | $p < 0.01$ |
| CRISPRa | $\log_2FC > 0.7$ | $p < 0.01$ | $p < 0.01$ |
| CRISPRi | $\log_2FC < 1$ | $p < 0.01$ | $p < 0.01$ |

### 2.2. Description of CRISPR-GEM Graphical User Interface and Differential Expression Analysis

CRISPR-GEM requires the input of the CRISPR strategy to be performed and the desired phenotypic changes sought following gene editing. A graphical user interface (GUI) is designed to facilitate this input and was made available through GitHub (https://github.com/jog822/CRISPR-GEM.git) (Supp. Fig. 1). The interface first requires the selection of the CRISPR strategy to be employed (CRISPR activation: “CRISPRa”, CRISPR inhibition: “CRISPRi”, CRISPR knock-out: “CRISPRko”, and CRISPR knock-in: “CRISPRki”). Next, the cell/tissue types for the gene editing strategy must be specified. The ‘end-to-end’ strategy requires the selection of the cell/tissue groups to be treated using CRISPR (experimental) and desired following this treatment (target). Additionally, the ‘intermediate cell type’ strategy allows for the additional input of a third intermediate group that lies on the progression from experimental to target groups such as a milder disease state or developing tissue. These groups may be selected using the dropdown menus and are all labelled in Supp. Table 1.

The interface then allows for feature selection using differential expression analysis to identify critical genes for distinguishing each group. To reduce bias in differential expression analysis, the GUI identifies the selected group with the lowest number of samples and randomly filters the remaining groups to have the same quantity. By clicking the “Get Inputs” button (Supp. Fig. 1F), the experimental, target, and possibly intermediate groups are used to create metadata. The pyDEseq2 algorithm then identifies DEGs between each group^25^. Significant DEGs are filtered according to the selected CRISPR strategy to determine the input and output genes for neural network analysis (Supp. Tables 5-10). Input genes are identified as the DEGs in the target group compared to the experimental group with the highest magnitude log_2_ fold change (log_2_FC) in the appropriate direction for the selected CRISPR strategy as described in **Table 1**. Output genes are identified as the genes with the lowest p value or most significant differential expression regardless of direction for a broad summary of the phenotypic differences.

When an intermediate cell/tissue type is identified, input genes are sorted by comparing the intermediate group instead of the target group while output gene selection is unchanged. The selected genes, statistical results, and the specific datasets used for the analysis may be saved to the location of the GUI executables by typing in a short description and clicking the “save” button. The selected genes and statistical analysis can be viewed in interactive datasets using the “Display Input Genes” or “Display Output Genes” buttons (Supp. Fig. 1F). For more detailed analysis of specific genes, the “Graph Gene” button can be used to plot the DeSeq2 normalized relative gene expression between each group. Further, the “Plot UMAP” button (Sup. Fig. 1H) can be applied to provide a two-dimensional projection describing the broad differences in the data selected for differential expression analysis^26^.

### 2.3. Deep Neural Network

Differential expression analysis was used to identify two distinct input and output gene sets of variable lengths based on the tissue or cell types compared. Each input and output node of the tissue specific MLP represents one of these genes, so the model architecture is defined by the number of input and output genes, respectively. The model connects the expression of input and output genes through three fully connected hidden layers constructed to represent CRISPR candidates’ interactions with the broader genome. This architecture allows learned relationships between input and output genes with the broader genome while preventing overfitting. Prior to model training, the tpm dataset prepared using kallisto is filtered for the input and output genes and any repeated genes are removed from the outputs yielding two unique datasets. Both datasets are then log normalized, split into 85% training and 15% testing subsets^27^. Finally, the model is trained using Keras built on top of TensorFlow with an rectified linear unit (ReLU) activation function, learning rate = 0.0001, ε = 1e-8, batch size = 50, validation split = 0.15, mean squared error (MSE) loss function, 200 training epochs and early stopping with a patience of 10^18,28,29^. The entire process from splitting into training and testing subsets to model training is repeated five times to cross-validate model performance.

### 2.4. Data Perturbation

The trained MLP predicts the gene expression of all output genes based on the expression of the input or CRISPR candidate genes; but further assessment is required to determine which genes have the most profound effect on directing cell phenotype. To assess these candidates, tpm data for the experimental group is filtered for CRISPR candidate genes before individually perturbing each gene to mimic the specified CRISPR strategy according to **Table 2**.

**Table 2.** Fold change perturbation for each CRISPR method.

| CRISPR Strategy | Perturbation | Equation # |
| --- | --- | --- |
| CRISPRko | $p = x \cdot 0$ | eq. 1 |
| CRISPRki | $p = x + k_i$ | eq. 2 |
| CRISPRa | $p = x \cdot \left(1 + \frac{k_a}{x}\right)$ | eq. 3 |
| CRISPRi | $p = \frac{x}{1 + \ln(x + 1)}$ | eq. 4 |

where: p=perturbed gene expression; x = unperturbed relative gene expression; $k_i$ is a constant defined based on knock-in promoter, set to 5 as default; and $k_a$ = constant that can be changed based on cell type, set to 15 as default.

CRISPRko is simulated by setting gene expression to zero to mimic the complete loss of gene expression in cultured knock-out cells **(Table 2 eq. 1)**. Since gene expression is a function of enhancer regions surrounding a gene and promoter strength, the transcriptomic changes resulting from CRISPRki are highly variable based on the knock-in coordinates and promoter used^30^. Therefore, we simulate knock-in by adding a constant expression value (k_i_) that simulates the addition of a second, independent copy of the gene and can be manually modified based on the gRNA and donor template sequences **(Table 2 eq. 2)**. The default k_i_ value is set to 5 since it is slightly above the average log normalized tpm value across all datasets of 35. The strong relationship between basal gene expression and activation or inhibition fold-changes is represented in **eqs. 3 and 4 (Table 2)** for CRISPRa and CRISPRi, respectively^31,32^. CRISPR activation levels are inversely correlated to basal expression which is modeled by multiplying the initial expression by a function of itself to dampen the activation in highly expressed genes **(Table 2 eq. 3)**. Alternatively, to simulate the positive correlation between CRISPRi and initial expression, the unperturbed expression is divided by a function of the natural logarithm of itself so that highly expressed genes are more greatly inhibited **(Table 2 eq. 4**).

### 2.4. Gene Evaluation

After perturbation, modified experimental group datasets for each gene are input into the DNN to predict the expression output genes following CRISPR editing. The predicted expression levels are compared to the model predictions for unperturbed target group datasets to assess how the perturbations shift the transcriptomic profile towards that of the target group. A scoring system was developed to incorporate the direction and magnitude of changes in gene expression and their benefit to the induction of the target phenotype. Briefly, for each perturbation input into the model, the predicted change in gene expression is calculated for every output gene as described in **eq. 5**:

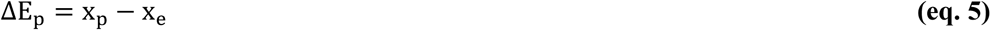

where: ΔE_p_ denotes the predicted change in gene expression, x_p_ represents the predicted gene expression from perturbed experimental data, and x_e_ represents the predicted expression from original experimental group data.

The predicted transcriptomic shifts following CRISPR editing are compared to model predictions for the experimental group to eliminate any significant bias from model training. Similarly, the desired change in gene expression (**eq. 6**) relates the predicted expression of the target group to the experimental group and represents the change in gene expression necessary for a fully effective therapy.

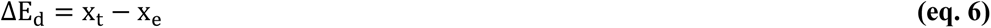

where: ΔE_d_ denotes the desired change in gene expression and x_t_ represents the predicted gene expression from non-perturbed target group data.

The final score is obtained by calculating the difference between the predicted and desired changes and normalizing to the initial gene expression value to account for the magnitude of that gene’s expression. Finally, the sum of the values for each gene is calculated and divided by the total number of output genes to yield the score for each CRISPR target gene **(eq. 7)**.

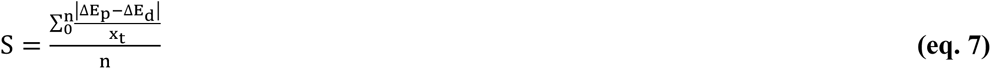

where: S represents the evaluation score for perturbed gene, and n represents the total number of output genes.

**Eq. 7** is used to assess the expression changes resultant of editing each CRISPR candidate and the lowest scoring genes are returned as the most promising candidates for the specified CRISPR strategy. The 50 highest scoring genes for each CRISPR-GEM application are input into STRING: Functional Protein Association Networks to visualize the protein-protein interactions that may occur downstream of the target genes^33^.

## 3. Results & Discussion

### 3.1. CRISPR-GEM Workflow

CRISPR-GEM represents a novel ML approach for the identification of promising target genes for specific therapeutic outputs. CRISPR-GEM constructs a synthetic GRN that captures the regulatory relationships within specific tissues then uses this to draw conclusions on the impact of CRISPR therapies **(Fig. 1)**. To validate CRISPR-GEM, we applied the algorithm to three regenerative medicine applications: **(1)** Regulatory T-cell differentiation from induced pluripotent stem cells (iPSCs), **(2)** Enhanced chondrogenesis of MSCs through CRISPRa, **(3)** the reversal of osteoarthritis progression using CRISPRi. Input/output gene selection and DNN training is performed consistently across each application with model predictions explaining over 95% of variance in testing data. For each prospective therapy we outline the motivation, group selection, and highlight promising candidate genes.

**Fig. 1.**
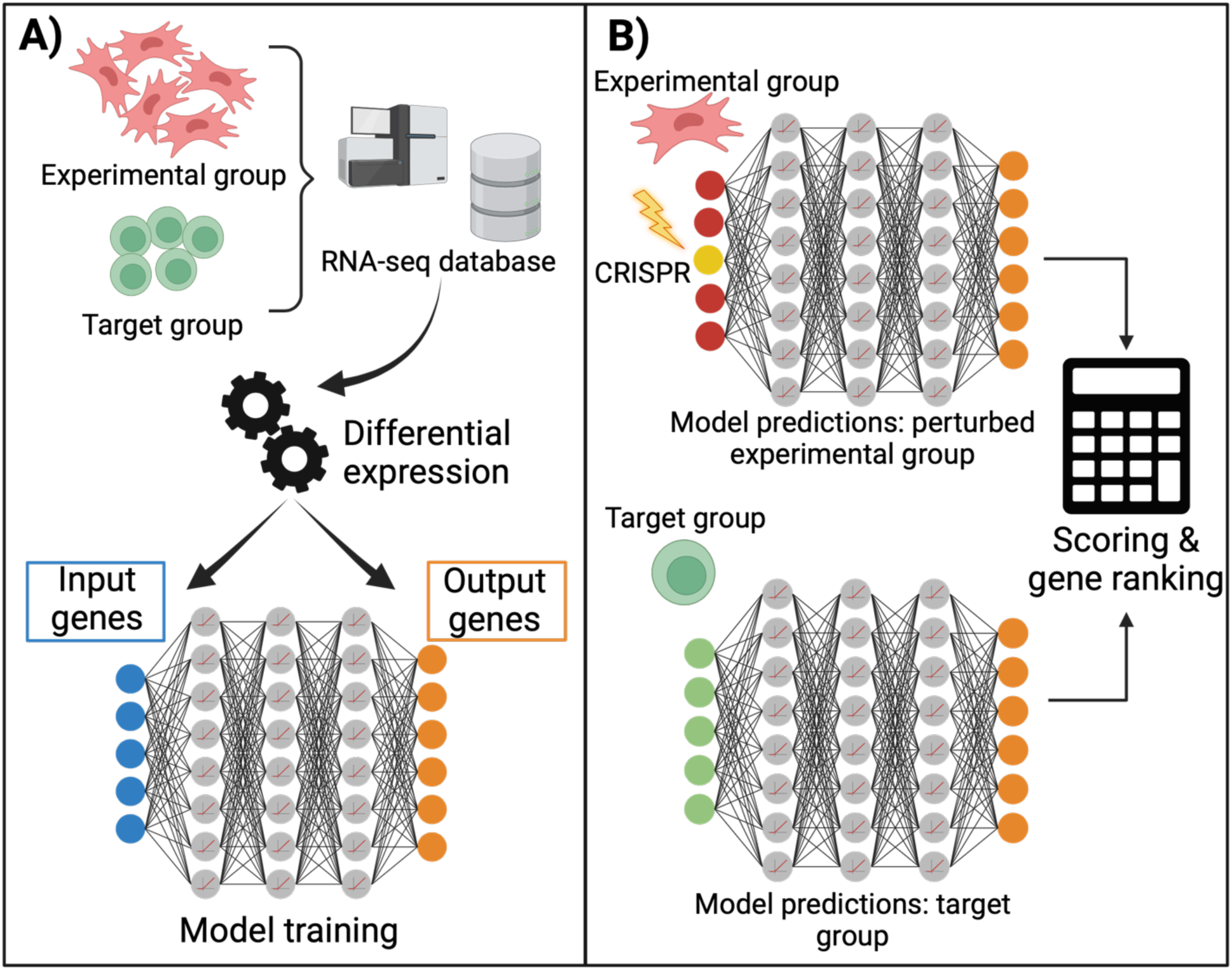
Schematic of CRISPR-GEM. **A)** Constructing tissue specific GRNs to predict gene expression relevant to the selected CRISPR therapy. DeSeq2 is utilized to select plausible CRISPR candidates and identify important genes for distinguishing experimental and target cell/tissue types. The resulting genes are used as input and output nodes respectively to train a MLP to predict the expression of output genes using only the expression of input genes. **B)** Assessing CRISPR strategies using the constructed GRN. RNA-seq data for the experimental group is perturbed by amplifying or suppressing the relative expression value of each gene individually. The perturbed experimental group dataset and an un-modified target group dataset are input into the trained MLP and model predictions for each are compared to assess the efficacy of the editing strategy for driving the target group transcriptomic profile.

### 3.2. Model Validation: Differentiation of iPSCs to Treg Cells

Tregs are a subset of CD4^+^CD25^+^ T-cells, that constitute 5-15% of peripheral T cells and help to control the adaptive immune response and mitigate inflammatory signals^34^. These immunosuppressive cells have shown great efficacy in inducing and maintaining immune homeostasis for the prevention of a range of autoimmune disorders such as diabetes, arthritis, and irritable bowel syndrome^35–39^. The main challenge in Treg and general T-cell therapies is the difficulty of cell generation, isolation, and expansion from patients. This issue could be solved through the differentiation and reprogramming of rapidly dividing iPSCs into Tregs^40^. Strategies to accomplish this employ viral vectors encoding Treg specific transgenes, namely *FOXP3,* and have successfully alleviated diabetes symptoms as well as inflammation and joint destruction in arthritic environments^38,40^. Despite the promise of these strategies, iPSCs hold the capacity to dedifferentiate or differentiate to various cell types including leukemia or lymphoma cells emphasizing the importance of reliable lineage commitment. To address this, we employed CRISPR-GEM to identify safe and reliable CRISPRa target genes to facilitate the differentiation of iPSCs to Treg cells.

iPSCs are defined as the experimental group with Treg cells as the target group. A large number of DEGs distinguished the iPSCs from Tregs as visualized in a low dimension UMAP projection (**Fig. 2A**). The separate iPSC clusters likely represent global differences stemming from variability in cell sourcing. Input and output gene selections were made by filtering DEGs between iPSCs and Tregs. Model training accurately captured the transcriptomic relationships between iPSC and Treg cell genes as demonstrated by a loss of 0.0*51* ± 0.002 (**Fig. 2B**) and r^2^ of 0.9*52* ± 0.002 (**Fig. 2C**). The accuracy of model predictions can further be visualized by linear trend between predicted and actual expression (**Fig. 2D**).

**Fig. 2.**
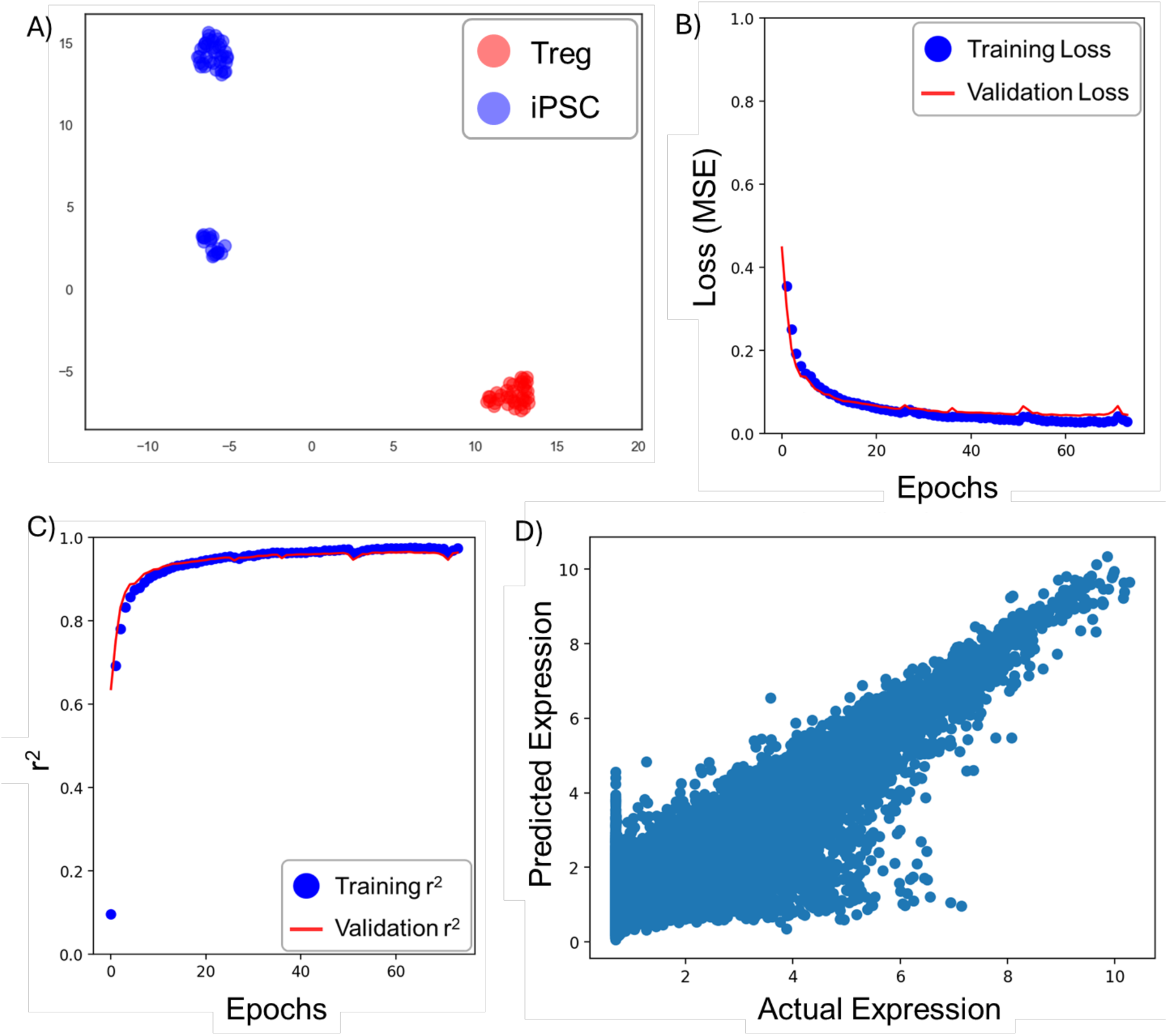
Model Training results given by CRISPR-GEM predictions for CRISPRa candidates to drive Treg differentiation. **A)** Treg cells display distinct phenotypes from iPSCs as visualized in reduced dimension via UMAP projection, **B)** Model training predicted output gene expression with an MSE of 0.051 *±* 0.002, **C)** an r^2^ of 0.952 *±* 0.002 (p = 1.11 · 10^−16^). These results suggest that the CRISPR-GEM neural network can predict over 94% of the transcriptomic variance throughout the iPSC-Treg transition, and **D)** The strong linear trend between actual and predicted output gene expression values verifies model accuracy.

Assessment of CRISPRa candidate genes reveals a highly connected regulatory network comprised mainly of immunomodulatory genes (**Fig. 3A)**. These genes are also connected to transcription factors such as *JUNB* and *KLF2* which are known to be critical regulators in the differentiation and function of Tregs^41,42^. These immunomodulatory genes may therefore act upstream or downstream of the transcription factors to drive phenotypic changes necessary for Treg differentiation. Tregs are also classified by their strong expression of forkhead box P3 (*FOXP3*), a transcription factor specific to regulatory T cells. Closer analysis of the top ten gene candidates reveals many known regulatory interactions or positive correlations to *FOXP3* expression (**Table 3**, **Fig. 3B**). Top scoring *SRGN* and high ranking *CD48* accomplish this through interactions with *BATF, a* transcription factor that is required for Treg differentiation, homeostasis and function^43–46^. *FOX3P* upregulation also occurs through downstream of *LTB, TSC22D3* genes*. LTB* contributes to the LTα1β2 heterotrimer that is highly expressed in Treg cells compared to other CD4^+^ subsets and directly correlates to *FOXP3 and CD25* expression^47^. *TSC22D3* encodes glucocorticoid-induced leucine zipper (GLIZ) which upregulates *FOXP3* and hence Treg production via the TGF-β and SMAD signaling pathways^48^. *CASP4* and *SELL* (CD26L) expression is not directly associated with *FOXP3* but has a strong positive correlation. *CASP4* encodes caspase 4, an endoprotease that regulates inflammation and has been notably upregulated during T cell activation^49^. *SELL* or L-selectin (CD-62) expression in Treg cells improves immunosuppressive activity in a range of autoimmune diseases^49,50^.

**Figure 3.**
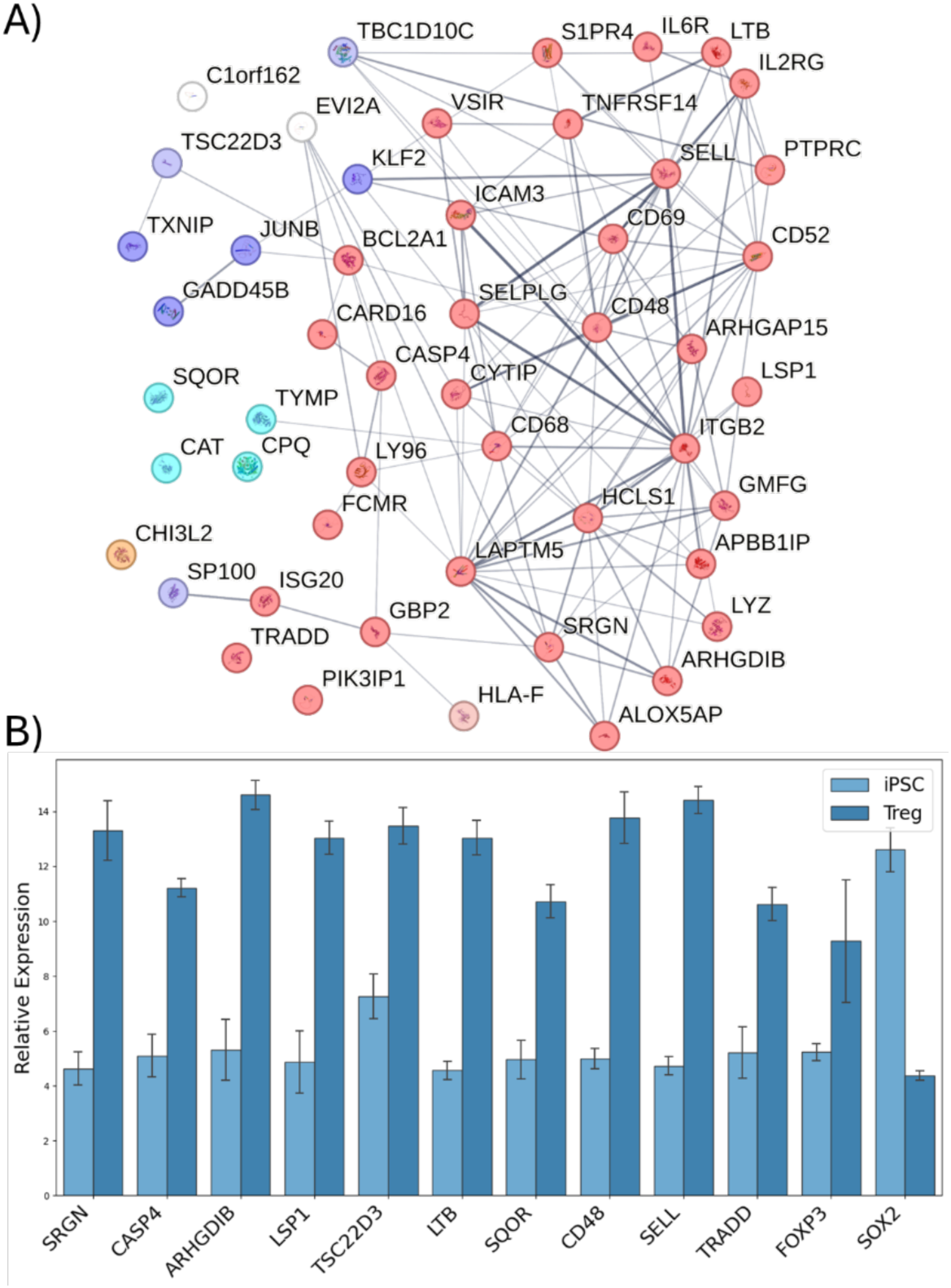
Assessment of model predictions for the optimal CRISPRa targets to drive Treg differentiation from iPSCs. **A)** STRING analysis of the top 50 ranked genes reveals a highly connected network of surface receptors and immunomodulatory genes predicted to strongly encourage iPSCs to commit to the Treg differentiation. The encoded proteins are associated with many pathways including immunomodulation (red), growth and differentiation (green), ECM composition (orange), cell structure (yellow), signal transduction (dark blue), transcription factors (light blue), and metabolism (teal). **B)** All gene targets identified by CRISPR-GEM are highly upregulated in Treg cells, most were more confidently upregulated than *FOXP3*, the genetic signature specific to Treg cells. Expression of *SOX2,* a pluripotency factor, is also given as a reference for iPSC function.

**Table 3.** Top scoring CRISPR candidates for CRISPRa induced Treg differentiation of iPSCs.

| Rank | CRISPR Candidate | Description | Reference(s) |
| --- | --- | --- | --- |
| 1 | *SRGN* | Encodes for serglycin, a proteoglycan and core marker for Tregs. It is universally upregulated in Tregs with strong positive correlation to basic leucine zipper ATF-like transcription factor (*BATF*), a recognized Treg transcription factor. | 43,44 |
| 2 | *CASP4* | Encodes for caspase 4. Upregulated during T-cell activation during a treatment for graft-versus-host disease. | 49 |
| 3 | *ARHGDIB* | Encodes for rho GDP dissociation inhibitor beta. Identified as an important regulator of T-cell activation. Inhibits Rho kinase resulting in greater Treg/Th17 ratio. | 51,52 |
| 4 | *LSP1* | Encodes for lymphocyte specific protein. Prompts the infiltration of regulatory T cells. Overexpression enhances expression of immunosuppressive genes. Deficiency is associated with rheumatoid arthritis. | 53,54 |
| 5 | *TSC22D3* | Encodes for glucocorticoid-induced leucine zipper (GLIZ). Enhances Treg production by upregulating *FOXP3* via TGF-β and SMAD signaling pathways. | 48 |
| 6 | *LTB* | Encodes for lymphotoxin beta (LT-β). It guides Treg migration and stability when bound to LT-α. | 55,56 |
| 7 | *SQOR* | Encodes for sulfide quinone oxidoreductase (SQOR). Uncharacterized in regulatory T-cells. |  |
| 8 | *CD48* | Encodes for CD48 which protects early effector T cells both by promoting autophagy and upregulation of pro-survival genes such as *BATF*. | 45 |
| 9 | *SELL* | Encodes for L-selectin (CD62). Tregs expressing L-selectin have superior immunosuppressive capacity roles in different diseases. | 49,50 |
| 10 | *TRADD* | Encodes for tumor necrosis factor receptor type 1-associated death domain protein. This protein is responsible for recruiting *TRAF2*, a gene that is upregulated during Treg infiltration. | 57,58 |

Besides these *FOXP3* modulators, *ARHGDIB, LY96,* and *LSP1* serve as immunomodulatory genes that regulate inflammatory environments*. ARHGDIB* is an important regulator of T-cell activation that inhibits Rho kinase causing higher proportions of Treg compared to Th17 T-cells^51,52^. *TRADD* participates in TNF signaling pathways and binds TRAF2 which aids in T-cell differentiation and signaling^57,58^. Finally, *LSP1* is a lymphocyte specific protein that prompts T-cell infiltration to suppress local inflammation^53,54^. The close association with the function of these genes and Tregs suggests that they could also impact T-cell differentiation and Treg activity.

### 3.3. Model Validation: Identifying CRISPRa Target Genes for Enhanced MSC Chondrogenesis

Degenerative and traumatic injuries to cartilage tissues have limited healing capacity due to a lack of vasculature and innervation which drives long-term pain and debilitation in many patients^59^. MSCs have emerged as a promising cell source to facilitate cartilage regeneration due to their accessibility, *in vitro* proliferative capacity, multipotency, and immunomodulatory properties^60^. The chondrogenic differentiation of MSCs is achieved by administering supraphysiological levels of growth factors such as transforming growth factor beta 3 (TGF-β3) or overexpressing transgenes^60–62^. However, these methods often result in phenotype instability and limited chondrogenic capacity *in vivo*. To address this, CRISPRa has been explored to upregulate specific genes that are associated with MSC chondrogenesis or chondrogenic ECM deposition to enhance this lineage commitment^3,63,64^. These strategies increased collagen type II deposition but led to incomplete chondrogenesis or had non-specific effects. For example, *COL2A1* upregulation induced DEGs associated with ECM remodeling but also endoplasmic reticulum stress and IREF1- mediated unfolded protein response^3,63^. Therefore, we implemented CRISPR-GEM to identify genetic targets for CRISPRa therapies and achieve mature chondrocyte gene expression patterns in MSCs.

We selected MSCs as the experimental group to be edited using CRISPRa and juvenile cartilage as the target group, representing healthy cartilage tissues. To further enhance model predictions, fetal chondrocytes were identified as an intermediate cell type representing developing cartilage. To illustrate the differences between each group, RNA-seq profiles are projected into a reduced dimension using UMAP revealing clear MSCs and cartilage clusters and distinct fetal and juvenile cartilage subpopulations **(Fig. 4A)**. Input and output genes were selected by filtering DEGs between MSCs to fetal cartilage and MSCs to mature cartilage respectively. Highly upregulated DEGs included *CNMD, UCMA, COL2A1, TGFB3, and SOX9* which are all strongly associated with cartilage tissues^65,66^. Neural network training across five replicates resulted in an average loss of 0.098 ± 0.018 and r^2^ of 0.952 ± 0.009 (p = 1.11 · 10^−16^). **(Fig. 4B and 4C)**. These results indicate that the DNN can predict over 96% of the variance amongst key MSC chondrogenic genes which can be visualized by the linear trend between actual and predicted gene expression in **Fig. 4D**. Following training, each CRISPR candidate gene in MSC RNA-seq datasets was perturbed using **eq. 3** to mimic CRISPRa gene upregulation. The resulting model predictions were input into our scoring system to rank CRISPR candidate genes for their capacity to induce chondrogenesis.

**Fig. 4.**
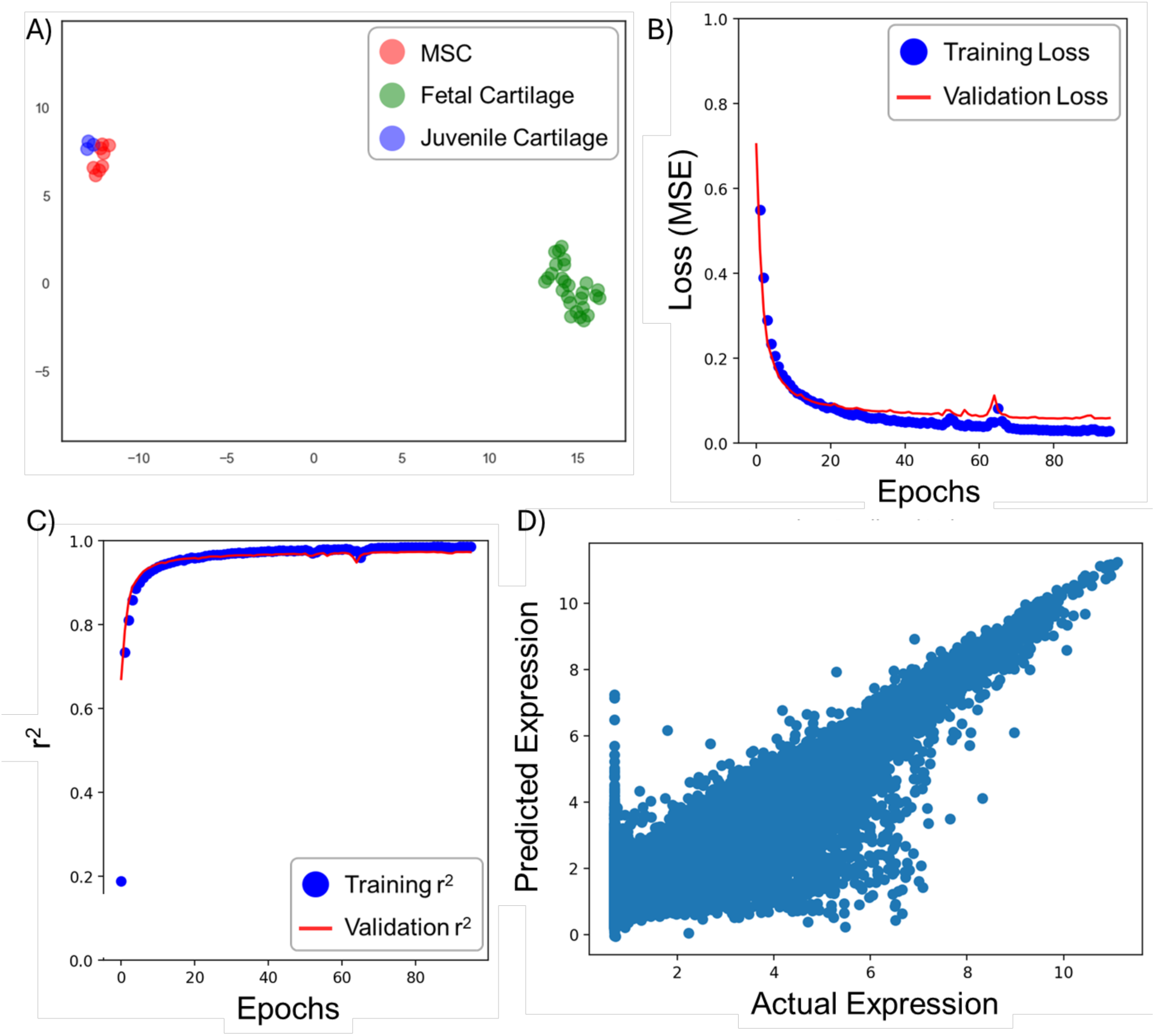
Group selection and model training for MSC chondrogenic characterization. MSCs, fetal cartilage, and mature cartilage were selected for experimental, intermediate, and target groups respectively. **A)** UMAP analysis confirms the clustering of cartilage cells and MSCs with fetal and mature cartilage subpopulations, **B)** The neural network was then trained with mean squared error (MSE) as the loss function, **C)** the resulting model was able to predict over 95% of the variance in the expression of chondrogenic genes as demonstrated by r^2^, and **D)** The model resulted in a clear linear trend between actual and predicted expression values for each gene.

The top scoring gene candidates yield a semi-connected STRING protein association network across a wide range of functionalities pathways, most notably ECM regulation and immunomodulation. This diversity suggests the complicated combination of signals required for driving signals and the importance of selecting CRISPR target genes that can modulate each of these pathways through GRNs **(Fig. 5A)**. *BMP2,* ranked sixth among all gene candidates, is highly connected within this network and is a potent inducer of MSC condensation, chondrogenic differentiation, chondrocyte proliferation and hypertrophic differentiation^67,68^. This result confirms the capacity of top scoring genes to influence GRNs and sustain a chondrogenic phenotype after activation. Further it demonstrates the importance of transient activation as modeled by CRISPR- GEM since sustained *BMP2* upregulation will cause cartilage hypertrophy and osteogenesis.

**Fig. 5.**
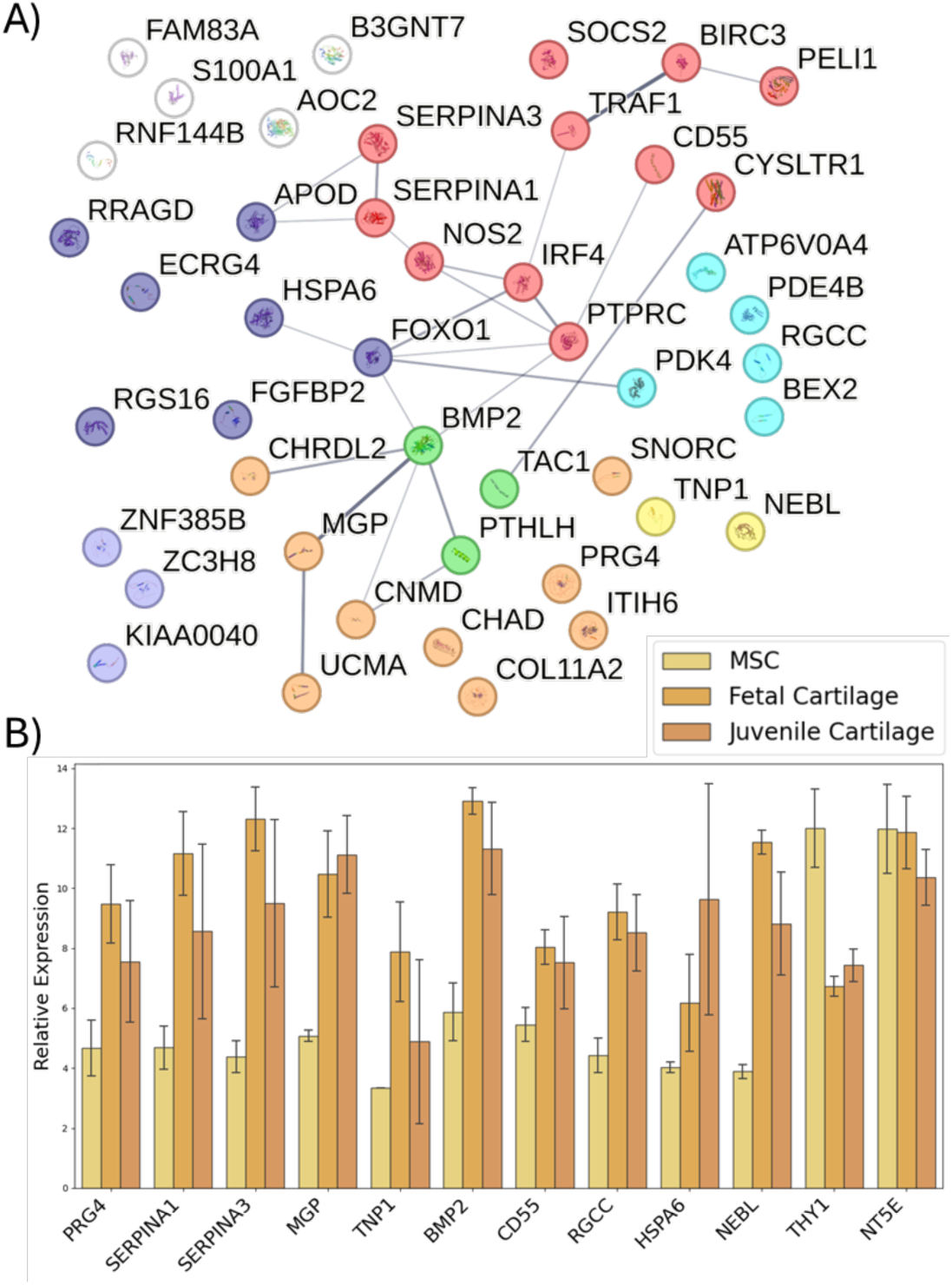
CRISPR-GEM results for CRISPRa induced chondrogenesis. **A)** STRING co-expression matrix for top scoring genes demonstrating the importance of ECM catabolism and immune regulation in the chondrogenic process. The encoded proteins are associated with many pathways including immunomodulation (red), growth and differentiation (green), ECM composition (orange), cell structure (yellow), signal transduction (dark blue), transcription factors (light blue), and metabolism (teal). **B)** All top scoring genes are significantly upregulated during the chondrogenic process. *THY1* and *NT5E*, two MSC surface receptors, demonstrate the shift from progenitor cells to chondrocytes.

The remaining top ten genes also hold great potential for MSC chondrogenesis **(Table 4**, **Fig. 5B)**. The top scoring *PRG4* encodes lubricin, a cartilage-specific glycoprotein that is necessary for regulating the surface active mucosal layer and chondroprotection^69–72^. Lubricin blocks cell adhesion and therefore could alter MSC microenvironments and stimulate cellular pathways to motivate chondrogenic differentiation. Interestingly, *SERPINA1* and *SERPINA3* inhibit the serine proteases elastase and cathepsin G respectively which both cleave lubricin and other important ECM components while mediating pro-inflammatory signaling^73^. *SERPINA1* and *SERPINA3* are also known chondrogenic markers suggesting that our top 3 gene candidates collaborate to establish a lubricin-based pro-chondrogenic matrix within MSC cultures^77–79^.

**Table 4.**
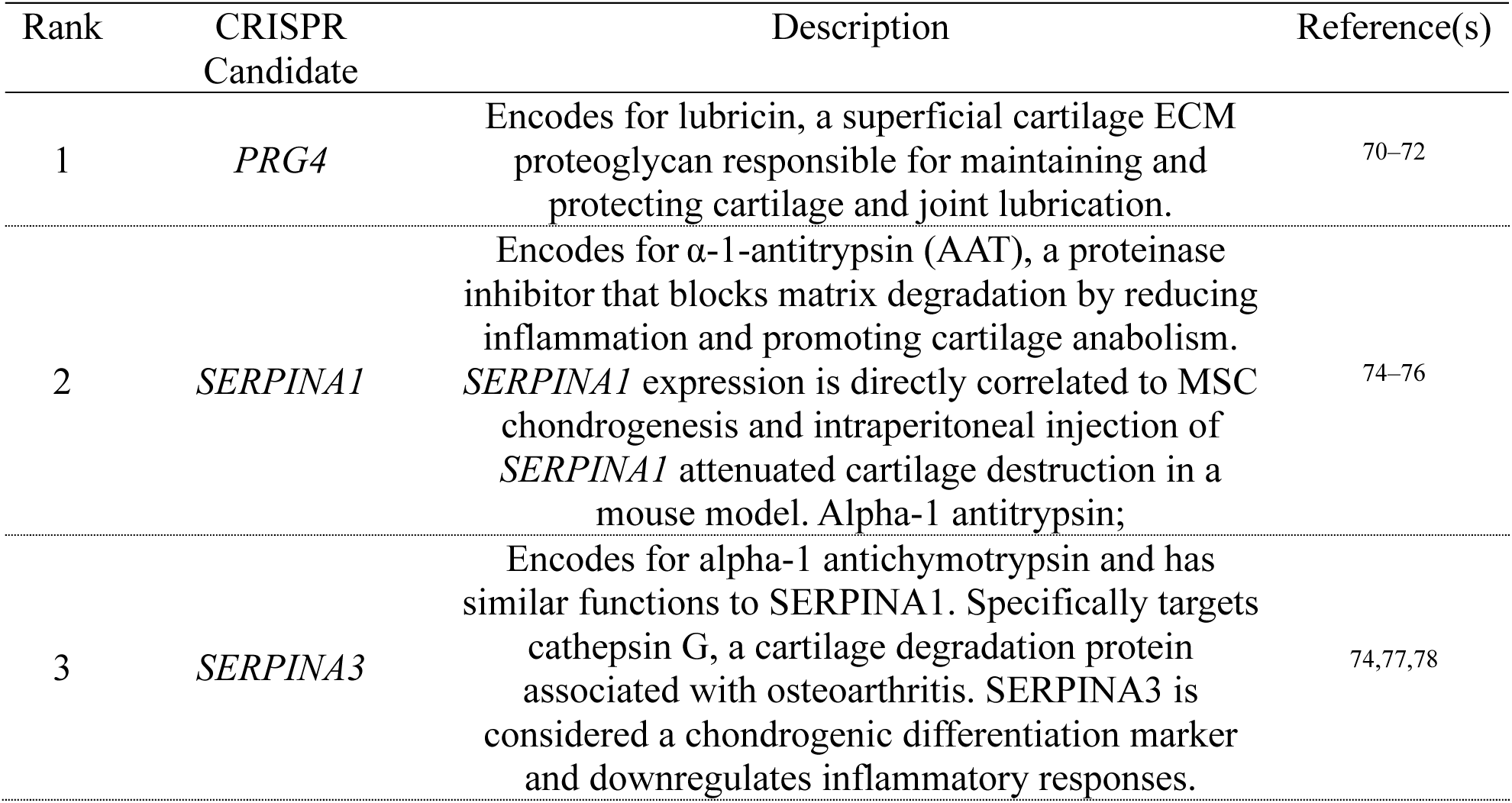
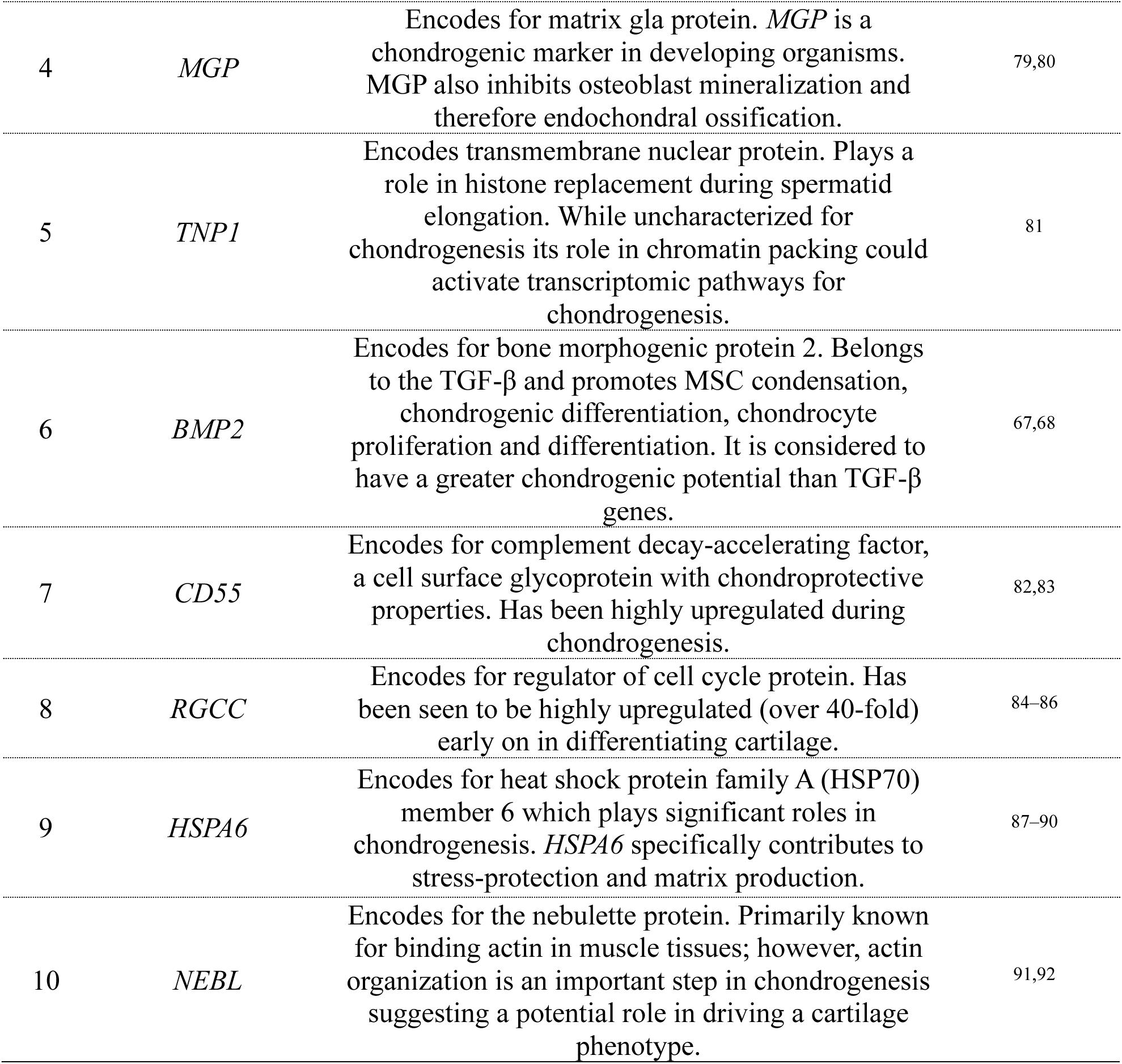
Top scoring CRISPR candidates for chondrogenic differentiation of MSCs.

In addition to the SERPINs, *MGP* and *CD55* also protect cartilage matrix during chondrogenesis. *MGP* is a chondrogenic marker in developing organisms that also inhibits osteoblast mineralization and therapy protecting cartilage tissues from endochondral ossification^79,80^. *CD55* is a cell surface glycoprotein that protects chondrocytes from immune degradation. The upregulation of these chondroprotective and immunomodulatory proteins could dysregulate cellular pathways that repress MSC chondrogenesis and represent a promising therapeutic strategy.

Beyond ECM and immune regulation, top scoring genes are also responsible for a range of other cell properties. *HSPA6* encodes HSP70, a stress-response protein that is known known to play essential roles in chondrogenesis and chondroprotection and is an established marker for cartilage homeostasis^90^ ^93^. *RGCC* is an important regulator of cell cycle that is highly upregulated during chondrogenesis and could encourage the metabolic changes that initiate the early stages of MSC differentiation ^84–86^. Finally, *NEBL* encodes the nebulette protein that is responsible for actin-binding in muscle cells but may contribute to the actin re-organization necessary for chondrogenic differentiation^91,92^.

Other notable high-scoring genes include *UCMA, CNMD,* and *SNORC* (ranked 13 and 14 and 22 respectively) encode cartilage specific proteins that comprise or regulate matrix production and control chondrocyte differentiation. *UCMA* interacts with collagens, inhibits aggrecanases, and protects cartilage to promote chondrogenesis^69,94^. *CNMD* stimulates chondrocyte growth while inhibiting angiogenesis to promote a sustained chondrocyte phenotype^95,96^. Finally, *SNORC* encodes for small novel rich in cartilage and is tightly controlled by SOX9 and is upregulated during chondrogenesis^97^.

In summary, CRISPR-GEM identified critical genes as the most promising CRISPR candidates to induce chondrogenesis of MSCs. Most of the top candidates have been reported to directly induce chondrogenic differentiation; however, the chondrogenic roles of other genes such as *RGCC and NEBL* remain to be fully characterized.

### 3.4. Model Validation: CRISPRi Target Selection for the Treatment of Osteoarthritis

Osteoarthritis (OA) is the most prevalent musculoskeletal disorder and leading cause of disability globally^98^. It is characterized by the presence of inflammatory signals that trigger cartilage degradation leading to joint pain and weakness^99^. CRISPR strategies for inhibiting OA-specific inflammation and matrix degradation pathways hold great potential for reversing the effects of osteoarthritis and restoring healthy cartilage. For example, CRISPRko of matrix metalloproteinase 13 (*MMP13*) and interleukin-1β (*IL1B*) genes has been reported to attenuate cartilage degradation. Alternatively, CRISPR has been used to alleviate OA induced pain through the knock-out nerve growth factor (*NGF*)^100^. Despite this promise, gene knock-out causes irreversible changes to the transcriptome and may have negative long-term impacts. Instead, CRISPRi based inhibition is more favorable for guiding cartilage tissues back to a healthy phenotype in a safe yet effective manner. Farhang *et al.* successfully suppressed immune response using CRISPRi to down-regulate tumor necrosis factor receptor 1 (*TNFR1*) and interleukin 1 receptor like 1 (*IL1R1*) expression; however, downregulating these genes failed to eliminate OA induced inflammation as seen by insufficient reduction in *IL1B* inflammatory pathways^101^. Therefore, we sought address the need for more well-informed target gene selection by identifying novel CRISPRi gene targets for reversing OA disease progression.

To accomplish this, osteoarthritic cartilage was compared to healthy cartilage as the experimental and target groups respectively. UMAP clustering reveals distinct clusters for each group revealing evident phenotypic differences in OA cartilage (**Fig. 6A)**. The neural network for OA genes was able to attain an MSE of 0.088 ± 0.0021 (**Fig. 6B**) and an r^2^ of 0.943 ± 0.001 (p = 1.11 · 10^−16^). (**Fig. 6C)** on testing data as visualized by the linear trend between predicted and actual expression values (**Fig 6D).**

**Fig. 6.**
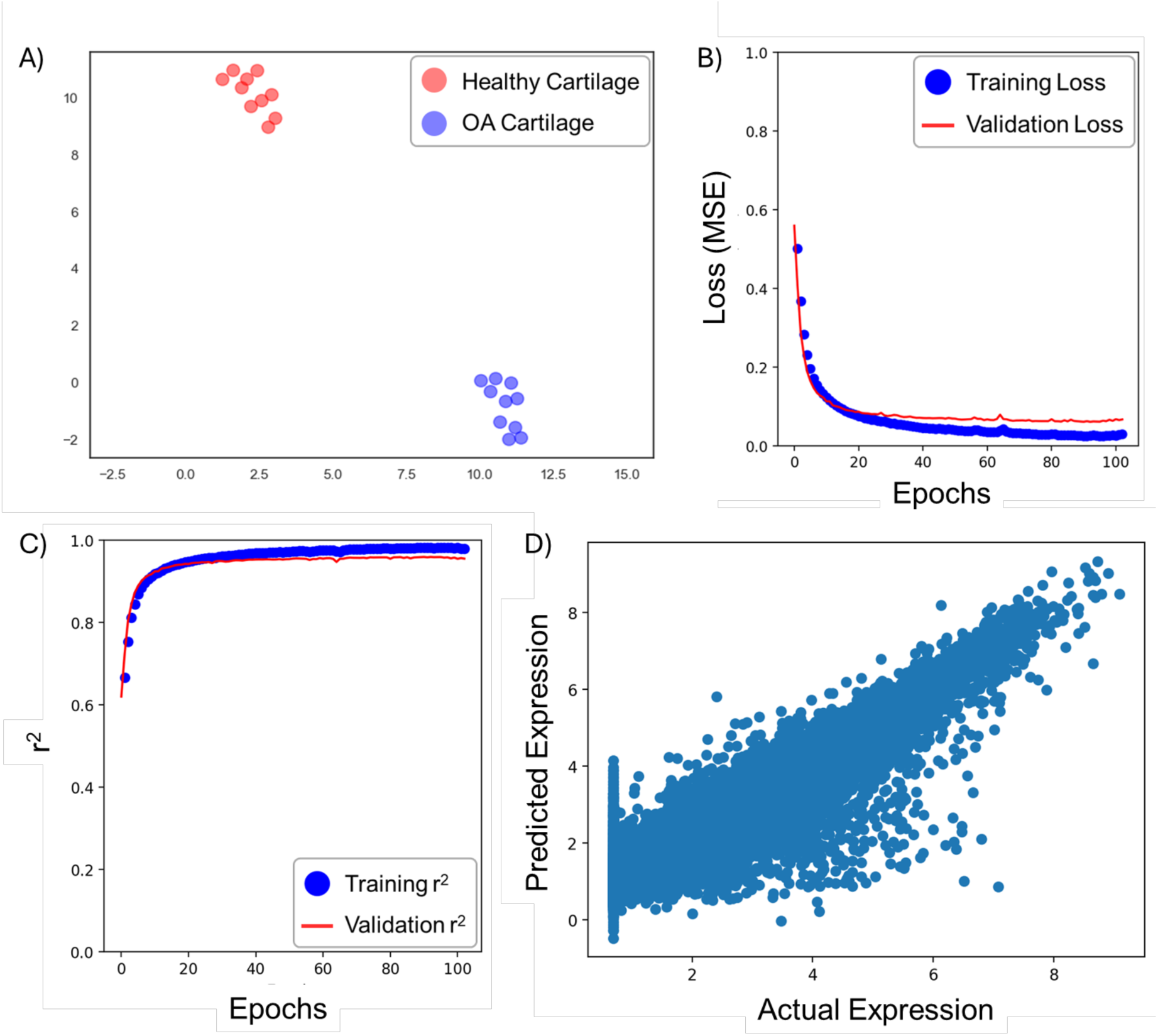
Group selection and model training for the reversal of OA using CRISPRi. Osteoarthritic and healthy cartilage tissues were selected as the experimental and target groups respectively. **A)** UMAP clustering confirms the substantial variance between diseased and healthy cartilage, **B)** Neural network training resulted in a mean squared error (MSE) of 0.088 ± 0.0021 while predicting output gene expression, **C)** prompting an r^2^ of 0.943 ± 0.001, and **D)** The model resulted in a clear linear trend between actual and predicted expression values for each gene.

The top scoring genes reveal a tightly connected ECM and immunomodulatory STRING network with interactions from a range of other cell processes (**Fig. 7A**). This network highlights the importance of matrix dysregulation in OA progression suggesting that inhibiting pro-fibrotic genes is an important therapeutic approach for attenuating OA. This observation correlates to *FAP* ranking as the top scoring CRISPRi candidate (**Table 5, Supp. Table 13**, **Fig. 7B**). *FAP* is an established biomarker and *in vitro* model for OA due to its role in ECM dysregulation through degrading denatured and MMP-13 cleaved type II collagen^102–104^. *MMP13,* an OA associated matrix metalloprotease, also ranks highly (33) amongst candidate genes due to its role in degrading types II, IV, and IX collagen, proteoglycans, osteonectin and perlecan^105^. In addition to ECM decomposition, both *COL1A1 and COL1A2* rank in the top 5 CRISPRi target genes. This suggests that the overproduction of type I collagen might cause fibrocartilage formation and therefore the loss of cartilage mechanical properties, contributing to pain and inflammation^106^. Another ECM-related gene ranked by CRISPR-GEM is *SPP1* (encoding for osteopontin), which is linked to matrix mineralization and therefore stiffening of cartilage tissues^107,108^. CRISPR-GEM clearly identified matrix dysregulation through the breakdown of healthy ECM and remodeling with inferior components as a crucial CRISPRi target.

**Table 5.** Top scoring CRISPR candidates for reversing OA.

| Rank | CRISPR Candidate | Description | Reference |
| --- | --- | --- | --- |
| 1 | *FAP* | Encodes for fibroblast activation protein (FAP) which collagen type II. A key genetic factor in OA that had been used to experimentally model OA. Inhibition of FAP ameliorates OA. | 102–104 |
| 2 | *IGFL4* | Encodes for insulin-like growth factor 4 (IGFL-4). Its role in OA and cartilage is uncharacterized. | 109 |
| 3 | *VCAM1* | Encodes for vascular cell adhesion molecule 1 (VCAM1) is upregulated during OA progression and provides adhesion sites for monocytes causing an influx of inflammatory signals further progressing OA. | 110–112 |
| 4 | *COL1A1* | Encodes for collagen type I which is a marker for fibrocartilage formation and its presence has been implicated as an early marker for osteoarthritis. | 106 |
| 5 | *COL1A2* | See *COL1A1* |  |
| 6 | *DIO2* | Encodes iodothyronine deiodinase 2 (DIO2), an enzyme that intracellularly activates thyroid hormone causing pro-inflammatory responses. Inhibition of DIO2 alleviates inflammatory systems in mice while overexpression causes immediate osteoarthritic symptoms. | 113,114 |
| 7 | *TNFAIP6* | Encodes for tumor necrosis factor alpha-induced protein 6 (TNFAIP6) which is highly regulated during osteoarthritis. It modulates macrophage phenotype and plays important roles in ECM remodeling, leukocyte migration and cell proliferation. | 115–117 |
| 8 | *CHI3L2* | Encodes for chitinase-3 like-protein-1, a growth and differentiation factor involved in cartilage homeostasis and a recognized biomarker for osteoarthritis. Highly expressed in earlier phases of degeneration. Chemotactic factor for monocytes and stimulates angiogenesis. | 118,119 |
| 9 | *SPP1* | Encodes for osteopontin, a phosphoprotein found in mineralized tissues and around inflammation sites involved in the cartilage to bone transition. Expression significantly correlates with disease severity. | 107,108 |
| 10 | *EDEM3* | Encodes for ER Degradation Enhancing Alpha-Mannosidase Like Protein 3. It is involved in endoplasmic reticulum stress and the degradation of misfolded glycoproteins. | 120,121 |

**Fig. 7.**
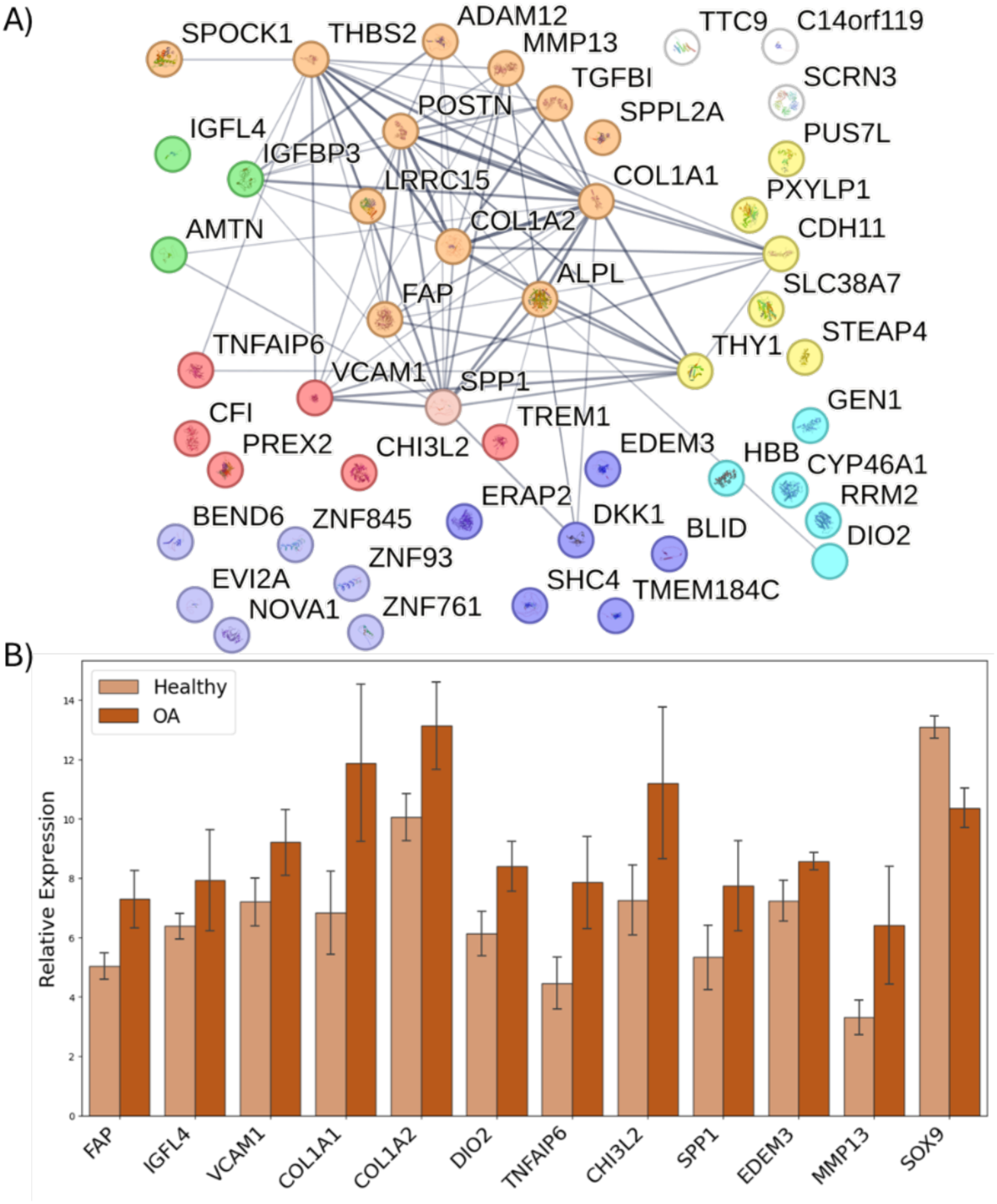
Visualizing the results for the top ranking CRISPRi targets to prevent OA. **A)** STRING co-expression matrix for top scoring genes shows a large network containing proteins involved in fibrosis, ECM catabolism, and immune regulation implicating these processes in OA. The encoded proteins are associated with many pathways including immunomodulation (red), growth and differentiation (green), ECM composition (orange), cell structure (yellow), signal transduction (dark blue), transcription factors (light blue), and metabolism (teal). **B)** The top ten scoring genes given by CRISPR-GEM are all upregulated during OA progression. These results positively correlated with MMP13 expression, a key OA marker; but they negatively correlated to the pro-regenerative, anti-inflammatory *SOX9*.

Immune-induced vascularization is another key driver of OA pathogenesis that CRISPR-GEM has identified. The chitinase-3 like-protein-1 protein encoded by *CHI3L2* is an established chemotactic factor for the recruitment of monocytes and a driver of angiogenesis. *CHI3L2* could therefore help to initiate the progressive inflammatory immune response responsible for OA^118,119^. Vascular cell adhesion molecule 1 (*VCAM1)* is another pro-angiogenic gene that provides adhesion sites for infiltrating monocytes therefore providing the means for the progressive worsening of inflammatory immune responses in cartilage tissues^110–112^. Macrophage infiltration and phenotype is also modulated by *TNFAIP6*, a member of the tumor necrosis factor family that is induced by IL-1β and TNF-α to facilitate ECM degradation and remodeling^115–117^. The inflammatory signals of *VCAM1 and TNFAIP6* may also be amplified by *DIO2* which intracellularly activates thyroid hormone causing inflammation in osteoarthritic patients^113,114^.

In addition to ECM dysregulation and immune response *EDEM3,* and *IGFL4* were identified as gene targets to prevent OA. *EDEM3* is responsible for degrading misfolded glycoproteins in the endoplasmic reticulum and is upregulated during OA^120,121^. Further study should be conducted to determine if this *EDEM3* upregulation causes abnormal degradation of functional glycoproteins. *IGFL4* is member of the insulin growth factor-like family that is yet to be characterized for its role in OA^109^. *EDEM3* and *IGFL4* serve as novel gene targets for OA prevention that must be evaluated.

## Conclusion

We have developed CRISPR-GEM, a tunable ML algorithm for the evaluation and discovery of CRISPR gene targets in a wide range of biomedical applications, from cell-specific differentiation to whole tissue disorders. This model uses differential expression to filter potential gene candidates for a desired application, then uses the selected genes to train an MLP that serves as a surrogate GRN for the specified tissue type. This neural network is subsequently utilized to assess the effects of CRISPR-mimetic perturbations to RNA-seq data and their capacity for inducing the desired cell phenotype.

To validate this machine learning model, we have applied it to identify CRISPRa gene targets for iPSC differentiation to Treg cells and the chondrogenic differentiation of MSCs, as well as the use of CRISPRi as a treatment for OA. In each application, CRISPR-GEM successfully identified both established and novel genes. All identified genes hold the potential to have a profound therapeutic impact for the given application and must be validated experimentally. Interrogation of these targets through *in vitro* CRISPR screens should be conducted to confirm the changes in gene expression predicted by CRISPR-GEM and evaluate therapeutic potential.

Modeling CRISPR target genes with CRISPR-GEM requires multiple assumptions: (1) Input and output gene feature selection is performed by filtering DEGs to eliminate unplausible gene candidates. This approach improves DNN accuracy by reducing noise, but it requires the assumption that genes following atypical expression patterns will not outperform the genes that positively correlate with the selected CRISPR editing strategy. Since CRISPR-GEM seeks to best recapitulate natural transcriptomic mechanisms this assumption helps reduce the risk of unintended downstream consequences by eliminating these unplausible gene candidates. (2) The selected genes are then utilized for MLP training to predict gene expression relationships and serve as a synthetic GRN. This MLP accurately predicts gene expression relationships without the need for detailed user input and therefore reduces bias and improves model accuracy compared to alternative techniques such as SVR or GNN. However, gene expression does not necessarily correlate to protein translation or post-translational modification^122^. The development of a model that explicitly accounts for all regulatory effects would require the input of multiple high-dimensional data types which complicates ML approaches and adds noise. Instead, CRISPR-GEM is built as a black box model to indirectly account for the effects of these regulatory mechanisms on the transcriptome. Therefore, this model assumes that the neural network is learning these relationships. (3) An additional limitation is that our MLP learns the regulatory relationships in gene expression without a temporal component. Thus, it is assumed that sustained gene activation can be achieved in the laboratory to accomplish the same transcriptomic impacts represented in CRISPR-GEM. Future work should consider collecting large datasets on the temporal development of primary tissues. This could be applied to training RNNs to explain the temporal aspects of tissue development or disease progression more definitively^123^.

In conclusion, CRISPR-GEM stands as the first ML based model for predicting optimal CRISPR target genes. We have tested this model in cell-specific differentiation and whole tissue disorders, demonstrating its potential for its application to a wide range of biomedical applications. Ultimately, this model will improve the efficacy and precision of CRISPR gene editing strategies.

## Acknowledgements

Tomas Gonzalez-Fernandez would like to acknowledge the start-up funds provided by the Department of Bioengineering and the P.C. Rossin College of Engineering & Applied Science at Lehigh University, the Orthoregeneration Network (ON) Kick Starter grant, and the Career Development Award from the American Society of Gene & Cell Therapy. The content is solely the responsibility of the authors and does not necessarily represent the official views of the American Society of Gene & Cell Therapy. Joshua P. Graham would like to acknowledge the support provided through the National Science Foundation Graduate Research Fellowship under Grant No #2234658. Lifang He’s research was partially supported by the NIH under grant R21EY034179, and the NSF through grants IIS-2319451 and MRI-2215789. Lifang He and Yu Zhang would like to acknowledge the support from the Lehigh University’s Accelerator and CORE grants.

## Notes

### Competing Interest Statement

The authors have declared no competing interest.

