## Supplementary material for "CRISPR-GEM: A Novel Machine Learning Model for CRISPR Genetic Target Discovery and Evaluation": Supp_Information_Biorxiv.pdf

**Josh Graham[1], Yu Zhang[1,2], Lifang He[3], Tomas Gonzalez-Fernandez[1]**
[1]Department of Bioengineering, Lehigh University, Bethlehem, PA, USA.
[2]Department of Electrical and Computer Engineering, Lehigh University, Bethlehem, PA, USA.
[3]Department of Computer Science and Engineering, Lehigh University, Bethlehem, PA, USA.

### SUPPLEMENTAL DATA

**Data Composition:**

**Supplemental Table 1.** Datasets used for model training and available for differential expression analysis.

| SRR | Label | Description | GSE | GSM | Platform | Ref. |
|---|---|---|---|---|---|---|
| SRR13018695 | MSC | Healthy | GSE161176 | GSM4891037 | HiSeq2500 | 1 |
| SRR13018696 | MSC | Healthy | GSE161176 | GSM4891038 | HiSeq2500 | 1 |
| SRR13018697 | MSC | Healthy | GSE161176 | GSM4891039 | HiSeq2500 | 1 |
| SRR13018698 | MSC | Treated | GSE161176 | GSM4891040 | HiSeq2500 | 1 |
| SRR13018699 | MSC | Treated | GSE161176 | GSM4891041 | HiSeq2500 | 1 |
| SRR13018700 | MSC | Treated | GSE161176 | GSM4891042 | HiSeq2500 | 1 |
| SRR13018701 | MSC | Treated | GSE161176 | GSM4891043 | HiSeq2500 | 1 |
| SRR13018702 | MSC | Treated | GSE161176 | GSM4891044 | HiSeq2500 | 1 |
| SRR13018703 | MSC | Treated | GSE161176 | GSM4891045 | HiSeq2500 | 1 |
| SRR13018704 | MSC | Treated | GSE161176 | GSM4891046 | HiSeq2500 | 1 |
| SRR13018705 | MSC | Treated | GSE161176 | GSM4891047 | HiSeq2500 | 1 |
| SRR13018706 | MSC | Treated | GSE161176 | GSM4891048 | HiSeq2500 | 1 |
| SRR13018707 | MSC | Treated | GSE161176 | GSM4891049 | HiSeq2500 | 1 |
| SRR13018708 | MSC | Treated | GSE161176 | GSM4891050 | HiSeq2500 | 1 |
| SRR13018709 | MSC | Treated | GSE161176 | GSM4891051 | HiSeq2500 | 1 |
| SRR13018710 | MSC | Treated | GSE161176 | GSM4891052 | HiSeq2500 | 1 |
| SRR13018711 | MSC | Treated | GSE161176 | GSM4891053 | HiSeq2500 | 1 |
| SRR13018712 | MSC | Treated | GSE161176 | GSM4891054 | HiSeq2500 | 1 |
| SRR13018713 | MSC | Treated | GSE161176 | GSM4891055 | HiSeq2500 | 1 |
| SRR13018714 | MSC | Treated | GSE161176 | GSM4891056 | HiSeq2500 | 1 |
| SRR13018715 | MSC | Treated | GSE161176 | GSM4891057 | HiSeq2500 | 1 |
| SRR13018716 | MSC | Treated | GSE161176 | GSM4891058 | HiSeq2500 | 1 |
| SRR13018717 | MSC | Treated | GSE161176 | GSM4891059 | HiSeq2500 | 1 |
| SRR13018718 | MSC | Treated | GSE161176 | GSM4891060 | HiSeq2500 | 1 |
| SRR13018719 | Cartilage | Healthy | GSE161176 | GSM4891061 | HiSeq2500 | 1 |
| SRR13018720 | Cartilage | Healthy | GSE161176 | GSM4891062 | HiSeq2500 | 1 |
| SRR13018721 | Cartilage | Healthy | GSE161176 | GSM4891063 | HiSeq2500 | 1 |
| SRR13018722 | Cartilage | Treated | GSE161176 | GSM4891064 | HiSeq2500 | 1 |
| SRR13018723 | Cartilage | Treated | GSE161176 | GSM4891065 | HiSeq2500 | 1 |
| SRR13018724 | Cartilage | Treated | GSE161176 | GSM4891066 | HiSeq2500 | 1 |
| SRR13018725 | Cartilage | Treated | GSE161176 | GSM4891067 | HiSeq2500 | 1 |
| SRR13018726 | Cartilage | Treated | GSE161176 | GSM4891068 | HiSeq2500 | 1 |
| SRR13018727 | Cartilage | Treated | GSE161176 | GSM4891069 | HiSeq2500 | 1 |
| SRR13018728 | Cartilage | Treated | GSE161176 | GSM4891070 | HiSeq2500 | 1 |
| SRR13018729 | Cartilage | Treated | GSE161176 | GSM4891071 | HiSeq2500 | 1 |
| SRR13018730 | Cartilage | Treated | GSE161176 | GSM4891072 | HiSeq2500 | 1 |
| SRR13018731 | Cartilage | Treated | GSE161176 | GSM4891073 | HiSeq2500 | 1 |
| SRR13018732 | Cartilage | Treated | GSE161176 | GSM4891074 | HiSeq2500 | 1 |
| SRR13018733 | Cartilage | Treated | GSE161176 | GSM4891075 | HiSeq2500 | 1 |
| SRR13018734 | Cartilage | Treated | GSE161176 | GSM4891076 | HiSeq2500 | 1 |
| SRR13018735 | Cartilage | Treated | GSE161176 | GSM4891077 | HiSeq2500 | 1 |
| SRR13018736 | Cartilage | Treated | GSE161176 | GSM4891078 | HiSeq2500 | 1 |
| SRR13018737 | Cartilage | Treated | GSE161176 | GSM4891079 | HiSeq2500 | 1 |
| SRR13018738 | Cartilage | Treated | GSE161176 | GSM4891080 | HiSeq2500 | 1 |

| | | | | | | |
|---|---|---|---|---|---|---|
| SRR13018739 | Cartilage | Treated | GSE161176 | GSM4891081 | HiSeq2500 | 1 |
| SRR13018740 | Cartilage | Treated | GSE161176 | GSM4891082 | HiSeq2500 | 1 |
| SRR13018741 | Cartilage | Treated | GSE161176 | GSM4891083 | HiSeq2500 | 1 |
| SRR13018742 | Cartilage | Treated | GSE161176 | GSM4891084 | HiSeq2500 | 1 |
| SRR13018743 | MSC | Healthy | GSE161176 | GSM4891085 | HiSeq2500 | 1 |
| SRR13018744 | MSC | Healthy | GSE161176 | GSM4891086 | HiSeq2500 | 1 |
| SRR13018745 | MSC | Healthy | GSE161176 | GSM4891087 | HiSeq2500 | 1 |
| SRR13018746 | MSC | Treated | GSE161176 | GSM4891088 | HiSeq2500 | 1 |
| SRR13018747 | MSC | Treated | GSE161176 | GSM4891089 | HiSeq2500 | 1 |
| SRR13018748 | MSC | Treated | GSE161176 | GSM4891090 | HiSeq2500 | 1 |
| SRR13018749 | MSC | Treated | GSE161176 | GSM4891091 | HiSeq2500 | 1 |
| SRR13018750 | MSC | Treated | GSE161176 | GSM4891092 | HiSeq2500 | 1 |
| SRR13018751 | MSC | Treated | GSE161176 | GSM4891093 | HiSeq2500 | 1 |
| SRR13018752 | MSC | Treated | GSE161176 | GSM4891094 | HiSeq2500 | 1 |
| SRR13018753 | MSC | Treated | GSE161176 | GSM4891095 | HiSeq2500 | 1 |
| SRR13018754 | MSC | Treated | GSE161176 | GSM4891096 | HiSeq2500 | 1 |
| SRR13018755 | MSC | Treated | GSE161176 | GSM4891097 | HiSeq2500 | 1 |
| SRR13018756 | MSC | Treated | GSE161176 | GSM4891098 | HiSeq2500 | 1 |
| SRR13018757 | MSC | Treated | GSE161176 | GSM4891099 | HiSeq2500 | 1 |
| SRR13018758 | MSC | Treated | GSE161176 | GSM4891100 | HiSeq2500 | 1 |
| SRR13018759 | MSC | Treated | GSE161176 | GSM4891101 | HiSeq2500 | 1 |
| SRR13018760 | MSC | Treated | GSE161176 | GSM4891102 | HiSeq2500 | 1 |
| SRR13018761 | MSC | Treated | GSE161176 | GSM4891103 | HiSeq2500 | 1 |
| SRR13018762 | MSC | Treated | GSE161176 | GSM4891104 | HiSeq2500 | 1 |
| SRR13018763 | MSC | Treated | GSE161176 | GSM4891105 | HiSeq2500 | 1 |
| SRR13018764 | MSC | Treated | GSE161176 | GSM4891106 | HiSeq2500 | 1 |
| SRR13018765 | MSC | Treated | GSE161176 | GSM4891107 | HiSeq2500 | 1 |
| SRR13018766 | MSC | Treated | GSE161176 | GSM4891108 | HiSeq2500 | 1 |
| SRR13018767 | MSC | Treated | GSE161176 | GSM4891109 | HiSeq2500 | 1 |
| SRR13018768 | MSC | Treated | GSE161176 | GSM4891110 | HiSeq2500 | 1 |
| SRR13018769 | MSC | Treated | GSE161176 | GSM4891111 | HiSeq2500 | 1 |
| SRR13018770 | MSC | Treated | GSE161176 | GSM4891112 | HiSeq2500 | 1 |
| SRR13018771 | MSC | Treated | GSE161176 | GSM4891113 | HiSeq2500 | 1 |
| SRR13018772 | MSC | Treated | GSE161176 | GSM4891114 | HiSeq2500 | 1 |
| SRR13018773 | Cartilage | Healthy | GSE161176 | GSM4891115 | HiSeq2500 | 1 |
| SRR13018774 | Cartilage | Healthy | GSE161176 | GSM4891116 | HiSeq2500 | 1 |
| SRR13018775 | Cartilage | Healthy | GSE161176 | GSM4891117 | HiSeq2500 | 1 |
| SRR13018776 | Cartilage | Treated | GSE161176 | GSM4891118 | HiSeq2500 | 1 |
| SRR13018777 | Cartilage | Treated | GSE161176 | GSM4891119 | HiSeq2500 | 1 |
| SRR13018778 | Cartilage | Treated | GSE161176 | GSM4891120 | HiSeq2500 | 1 |
| SRR13018779 | Cartilage | Treated | GSE161176 | GSM4891121 | HiSeq2500 | 1 |
| SRR13018780 | Cartilage | Treated | GSE161176 | GSM4891122 | HiSeq2500 | 1 |
| SRR13018781 | Cartilage | Treated | GSE161176 | GSM4891123 | HiSeq2500 | 1 |
| SRR13018782 | Cartilage | Treated | GSE161176 | GSM4891124 | HiSeq2500 | 1 |
| SRR13018783 | Cartilage | Treated | GSE161176 | GSM4891125 | HiSeq2500 | 1 |
| SRR13018784 | Cartilage | Treated | GSE161176 | GSM4891126 | HiSeq2500 | 1 |

| | | | | | | |
|---|---|---|---|---|---|---|
| SRR13018785 | Cartilage | Treated | GSE161176 | GSM4891127 | HiSeq2500 | [1] |
| SRR13018786 | Cartilage | Treated | GSE161176 | GSM4891128 | HiSeq2500 | [1] |
| SRR13018787 | Cartilage | Treated | GSE161176 | GSM4891129 | HiSeq2500 | [1] |
| SRR7107481 | Cartilage | Healthy | GSE114007 | GSM3130531 | HiSeq2000 | [2] |
| SRR7107483 | Cartilage | Healthy | GSE114007 | GSM3130533 | HiSeq2000 | [2] |
| SRR7107484 | Cartilage | Healthy | GSE114007 | GSM3130534 | HiSeq2000 | [2] |
| SRR7107486 | Cartilage | Healthy | GSE114007 | GSM3130536 | HiSeq2000 | [2] |
| SRR7107487 | Cartilage | Healthy | GSE114007 | GSM3130537 | HiSeq2000 | [2] |
| SRR7107488 | Cartilage | Healthy | GSE114007 | GSM3130538 | HiSeq2000 | [2] |
| SRR7107489 | Cartilage | OA | GSE114007 | GSM3130539 | HiSeq2000 | [2] |
| SRR7107490 | Cartilage | OA | GSE114007 | GSM3130540 | HiSeq2000 | [2] |
| SRR7107491 | Cartilage | OA | GSE114007 | GSM3130541 | HiSeq2000 | [2] |
| SRR7107492 | Cartilage | OA | GSE114007 | GSM3130542 | HiSeq2000 | [2] |
| SRR7107493 | Cartilage | OA | GSE114007 | GSM3130543 | HiSeq2000 | [2] |
| SRR7107494 | Cartilage | OA | GSE114007 | GSM3130544 | HiSeq2000 | [2] |
| SRR7107495 | Cartilage | OA | GSE114007 | GSM3130545 | HiSeq2000 | [2] |
| SRR7107496 | Cartilage | OA | GSE114007 | GSM3130546 | HiSeq2000 | [2] |
| SRR7107497 | Cartilage | OA | GSE114007 | GSM3130547 | HiSeq2000 | [2] |
| SRR7107498 | Cartilage | OA | GSE114007 | GSM3130548 | HiSeq2000 | [2] |
| SRR6337326 | Cartilage | Juvenile | GSE106292 | GSM2871635 | HiSeq2500 | [3,4] |
| SRR6337327 | Cartilage | Juvenile | GSE106292 | GSM2871636 | HiSeq2500 | [3,4] |
| SRR6337328 | Cartilage | Juvenile | GSE106292 | GSM2871637 | HiSeq2500 | [3,4] |
| SRR6337329 | Cartilage | Juvenile | GSE106292 | GSM2871638 | HiSeq2500 | [3,4] |
| SRR6337330 | Cartilage | Juvenile | GSE106292 | GSM2871639 | HiSeq2500 | [3,4] |
| SRR6337331 | Cartilage | Juvenile | GSE106292 | GSM2871640 | HiSeq2500 | [3,4] |
| SRR6337332 | Cartilage | Juvenile | GSE106292 | GSM2871641 | HiSeq2500 | [3,4] |
| SRR6337333 | Cartilage | Juvenile | GSE106292 | GSM2871642 | HiSeq2500 | [3,4] |
| SRR6337334 | Cartilage | Juvenile | GSE106292 | GSM2871643 | HiSeq2500 | [3,4] |
| SRR6337335 | Cartilage | Juvenile | GSE106292 | GSM2871644 | HiSeq2500 | [3,4] |
| SRR6337336 | Cartilage | Juvenile | GSE106292 | GSM2871645 | HiSeq2500 | [3,4] |
| SRR6337337 | Cartilage | Juvenile | GSE106292 | GSM2871646 | HiSeq2500 | [3,4] |
| SRR6337338 | Cartilage | Juvenile | GSE106292 | GSM2871647 | HiSeq2500 | [3,4] |
| SRR3536062 | Cartilage | Fetal | GSE81526 | GSM2155101 | HiSeq2500 | [5] |
| SRR3536063 | Cartilage | Fetal | GSE81526 | GSM2155102 | HiSeq2500 | [5] |
| SRR3536064 | Cartilage | Fetal | GSE81526 | GSM2155103 | HiSeq2500 | [5] |
| SRR3536065 | Cartilage | Fetal | GSE81526 | GSM2155104 | HiSeq2500 | [5] |
| SRR3536066 | Cartilage | Healthy | GSE81526 | GSM2155105 | HiSeq2500 | [5] |
| SRR3536067 | Cartilage | Healthy | GSE81526 | GSM2155106 | HiSeq2500 | [5] |
| SRR3536068 | Cartilage | Healthy | GSE81526 | GSM2155107 | HiSeq2500 | [5] |
| SRR3536069 | Cartilage | Cultured | GSE81526 | GSM2155108 | HiSeq2500 | [5] |
| SRR3536070 | Cartilage | Cultured | GSE81526 | GSM2155109 | HiSeq2500 | [5] |
| SRR3536071 | Cartilage | Cultured | GSE81526 | GSM2155110 | HiSeq2500 | [5] |
| SRR3536072 | Cartilage | Cultured | GSE81526 | GSM2155111 | HiSeq2500 | [5] |
| SRR3536073 | Cartilage | Cultured | GSE81526 | GSM2155112 | HiSeq2500 | [5] |
| SRR3536074 | Cartilage | Cultured | GSE81526 | GSM2155113 | HiSeq2500 | [5] |
| SRR3536075 | Cartilage | Cultured | GSE81526 | GSM2155114 | HiSeq2500 | [5] |

| | | | | | | |
|---|---|---|---|---|---|---|
| SRR3536076 | Cartilage | Treated | GSE81526 | GSM2155115 | HiSeq2500 | 5 |
| SRR3536077 | Cartilage | Treated | GSE81526 | GSM2155116 | HiSeq2500 | 5 |
| SRR3536078 | Cartilage | Treated | GSE81526 | GSM2155117 | HiSeq2500 | 5 |
| SRR3536079 | Cartilage | Treated | GSE81526 | GSM2155118 | HiSeq2500 | 5 |
| SRR3536080 | Cartilage | Treated | GSE81526 | GSM2155119 | HiSeq2500 | 5 |
| SRR3536081 | Cartilage | Treated | GSE81526 | GSM2155120 | HiSeq2500 | 5 |
| SRR3536082 | Cartilage | Treated | GSE81526 | GSM2155121 | HiSeq2500 | 5 |
| SRR6321830 | Cartilage | Injured | GSE111358 | GSM2864660 | HiSeq2500 | 6,7 |
| SRR6321831 | Cartilage | Injured | GSE111358 | GSM2864661 | HiSeq2500 | 6,7 |
| SRR6321832 | Cartilage | Injured | GSE111358 | GSM2864662 | HiSeq2500 | 6,7 |
| SRR6321833 | Cartilage | Injured | GSE111358 | GSM2864663 | HiSeq2500 | 6,7 |
| SRR6321834 | Cartilage | Injured | GSE111358 | GSM2864664 | HiSeq2500 | 6,7 |
| SRR6321835 | Cartilage | Injured | GSE111358 | GSM2864665 | HiSeq2500 | 6,7 |
| SRR6798765 | Cartilage | OA | GSE111358 | GSM3029071 | HiSeq2500 | 6,7 |
| SRR6798766 | Cartilage | OA | GSE111358 | GSM3029072 | HiSeq2500 | 6,7 |
| SRR6798767 | Cartilage | OA | GSE111358 | GSM3029073 | HiSeq2500 | 6,7 |
| SRR6798768 | Cartilage | OA | GSE111358 | GSM3029074 | HiSeq2500 | 6,7 |
| SRR6798769 | Cartilage | OA | GSE111358 | GSM3029075 | HiSeq2500 | 6,7 |
| SRR6798770 | Cartilage | OA | GSE111358 | GSM3029076 | HiSeq2500 | 6,7 |
| SRR6798771 | Cartilage | OA | GSE111358 | GSM3029077 | HiSeq2500 | 6,7 |
| SRR6798772 | Cartilage | OA | GSE111358 | GSM3029078 | HiSeq2500 | 6,7 |
| SRR6798773 | Cartilage | OA | GSE111358 | GSM3029079 | HiSeq2500 | 6,7 |
| SRR6798774 | Cartilage | OA | GSE111358 | GSM3029080 | HiSeq2500 | 6,7 |
| SRR17190416 | Cartilage | Healthy | GSE190615 | GSM5726497 | HiSeq2500 | 8 |
| SRR17190417 | Cartilage | Healthy | GSE190615 | GSM5726498 | HiSeq2500 | 8 |
| SRR17190418 | Cartilage | Healthy | GSE190615 | GSM5726499 | HiSeq2500 | 8 |
| SRR17190419 | Cartilage | Healthy | GSE190615 | GSM5726500 | HiSeq2500 | 8 |
| SRR17190420 | Cartilage | Healthy | GSE190615 | GSM5726501 | HiSeq2500 | 8 |
| SRR17190421 | Cartilage | Healthy | GSE190615 | GSM5726502 | HiSeq2500 | 8 |
| SRR17190422 | Cartilage | Healthy | GSE190615 | GSM5726503 | HiSeq2500 | 8 |
| SRR17190423 | Cartilage | Healthy | GSE190615 | GSM5726504 | HiSeq2500 | 8 |
| SRR17190504 | Bone | Healthy | GSE190615 | GSM5726585 | HiSeq2500 | 8 |
| SRR17190505 | Bone | Healthy | GSE190615 | GSM5726586 | HiSeq2500 | 8 |
| SRR17190506 | Bone | Healthy | GSE190615 | GSM5726587 | HiSeq2500 | 8 |
| SRR17190507 | Bone | Healthy | GSE190615 | GSM5726588 | HiSeq2500 | 8 |
| SRR17190508 | Bone | Healthy | GSE190615 | GSM5726589 | HiSeq2500 | 8 |
| SRR17190509 | Bone | Healthy | GSE190615 | GSM5726590 | HiSeq2500 | 8 |
| SRR17190510 | Bone | Healthy | GSE190615 | GSM5726591 | HiSeq2500 | 8 |
| SRR17190511 | Bone | Healthy | GSE190615 | GSM5726592 | HiSeq2500 | 8 |
| SRR17190568 | Muscle | Healthy | GSE190615 | GSM5726649 | HiSeq2500 | 8 |
| SRR17190569 | Muscle | Healthy | GSE190615 | GSM5726650 | HiSeq2500 | 8 |
| SRR17190570 | Muscle | Healthy | GSE190615 | GSM5726651 | HiSeq2500 | 8 |
| SRR17190571 | Muscle | Healthy | GSE190615 | GSM5726652 | HiSeq2500 | 8 |
| SRR17190572 | Muscle | Healthy | GSE190615 | GSM5726653 | HiSeq2500 | 8 |
| SRR17190573 | Muscle | Healthy | GSE190615 | GSM5726654 | HiSeq2500 | 8 |
| SRR17190574 | Muscle | Healthy | GSE190615 | GSM5726655 | HiSeq2500 | 8 |

| | | | | | | |
|---|---|---|---|---|---|---|
| SRR17190575 | Muscle | Healthy | GSE190615 | GSM5726656 | HiSeq2500 | [8] |
| SRR10603518 | Bone | Diseased | GSE141614 | GSM4209329 | HiSeq2000 | [9] |
| SRR10603519 | Bone | Diseased | GSE141614 | GSM4209330 | HiSeq2000 | [9] |
| SRR10603520 | Bone | Diseased | GSE141614 | GSM4209331 | HiSeq2000 | [9] |
| SRR10603521 | Bone | Diseased | GSE141614 | GSM4209332 | HiSeq2000 | [9] |
| SRR10603522 | Bone | Diseased | GSE141614 | GSM4209333 | HiSeq2000 | [9] |
| SRR10603523 | Bone | Diseased | GSE141614 | GSM4209334 | HiSeq2000 | [9] |
| SRR10603524 | Bone | Diseased | GSE141614 | GSM4209335 | HiSeq2000 | [9] |
| SRR10603525 | Bone | Diseased | GSE141614 | GSM4209336 | HiSeq2000 | [9] |
| SRR10603526 | Bone | Diseased | GSE141614 | GSM4209337 | HiSeq2000 | [9] |
| SRR10603527 | Bone | Diseased | GSE141614 | GSM4209338 | HiSeq2000 | [9] |
| SRR10603528 | Bone | Diseased | GSE141614 | GSM4209339 | HiSeq2000 | [9] |
| SRR10603529 | Bone | Diseased | GSE141614 | GSM4209340 | HiSeq2000 | [9] |
| SRR10603530 | Bone | Diseased | GSE141614 | GSM4209341 | HiSeq2000 | [9] |
| SRR10603531 | Bone | Diseased | GSE141614 | GSM4209342 | HiSeq2000 | [9] |
| SRR10603532 | Bone | Diseased | GSE141614 | GSM4209343 | HiSeq2000 | [9] |
| SRR10603533 | Bone | Healthy | GSE141614 | GSM4209344 | HiSeq2000 | [9] |
| SRR10603534 | Bone | Healthy | GSE141614 | GSM4209345 | HiSeq2000 | [9] |
| SRR10603535 | Bone | Healthy | GSE141614 | GSM4209346 | HiSeq2000 | [9] |
| SRR10603536 | Bone | Healthy | GSE141614 | GSM4209347 | HiSeq2000 | [9] |
| SRR10603537 | Bone | Healthy | GSE141614 | GSM4209348 | HiSeq2000 | [9] |
| SRR10603538 | Bone | Healthy | GSE141614 | GSM4209349 | HiSeq2000 | [9] |
| SRR10603539 | Bone | Healthy | GSE141614 | GSM4209350 | HiSeq2000 | [9] |
| SRR10603540 | Bone | Healthy | GSE141614 | GSM4209351 | HiSeq2000 | [9] |
| SRR10603541 | Bone | Healthy | GSE141614 | GSM4209352 | HiSeq2000 | [9] |
| SRR10603542 | Bone | Healthy | GSE141614 | GSM4209353 | HiSeq2000 | [9] |
| SRR10603543 | Bone | Healthy | GSE141614 | GSM4209354 | HiSeq2000 | [9] |
| SRR10603544 | Bone | Healthy | GSE141614 | GSM4209355 | HiSeq2000 | [9] |
| SRR10603545 | Bone | Healthy | GSE141614 | GSM4209356 | HiSeq2000 | [9] |
| SRR10603546 | Bone | Healthy | GSE141614 | GSM4209357 | HiSeq2000 | [9] |
| SRR10603547 | Bone | Healthy | GSE141614 | GSM4209358 | HiSeq2000 | [9] |
| SRR10603694 | Bone | Diseased | GSE141614 | GSM4209502 | HiSeq2000 | [9] |
| SRR10603695 | Bone | Diseased | GSE141614 | GSM4209503 | HiSeq2000 | [9] |
| SRR10603696 | Bone | Diseased | GSE141614 | GSM4209504 | HiSeq2000 | [9] |
| SRR10603697 | Bone | Diseased | GSE141614 | GSM4209505 | HiSeq2000 | [9] |
| SRR10603698 | Bone | Diseased | GSE141614 | GSM4209506 | HiSeq2000 | [9] |
| SRR10603699 | Bone | Diseased | GSE141614 | GSM4209507 | HiSeq2000 | [9] |
| SRR10603700 | Bone | Diseased | GSE141614 | GSM4209508 | HiSeq2000 | [9] |
| SRR10603701 | Bone | Diseased | GSE141614 | GSM4209509 | HiSeq2000 | [9] |
| SRR10603702 | Bone | Diseased | GSE141614 | GSM4209510 | HiSeq2000 | [9] |
| SRR10603703 | Bone | Diseased | GSE141614 | GSM4209511 | HiSeq2000 | [9] |
| SRR10603704 | Bone | Diseased | GSE141614 | GSM4209512 | HiSeq2000 | [9] |
| SRR10603705 | Bone | Diseased | GSE141614 | GSM4209513 | HiSeq2000 | [9] |
| SRR10603706 | Bone | Diseased | GSE141614 | GSM4209514 | HiSeq2000 | [9] |
| SRR10603707 | Bone | Diseased | GSE141614 | GSM4209515 | HiSeq2000 | [9] |
| SRR10603708 | Bone | Diseased | GSE141614 | GSM4209516 | HiSeq2000 | [9] |

| | | | | | | |
|---|---|---|---|---|---|---|
| SRR10603709 | Bone | Healthy | GSE141614 | GSM4209517 | HiSeq2000 | [9] |
| SRR10603710 | Bone | Healthy | GSE141614 | GSM4209518 | HiSeq2000 | [9] |
| SRR10603712 | Bone | Healthy | GSE141614 | GSM4209520 | HiSeq2000 | [9] |
| SRR10603713 | Bone | Healthy | GSE141614 | GSM4209521 | HiSeq2000 | [9] |
| SRR10603714 | Bone | Healthy | GSE141614 | GSM4209522 | HiSeq2000 | [9] |
| SRR10603716 | Bone | Healthy | GSE141614 | GSM4209524 | HiSeq2000 | [9] |
| SRR10603718 | Bone | Healthy | GSE141614 | GSM4209526 | HiSeq2000 | [9] |
| SRR10603719 | Bone | Healthy | GSE141614 | GSM4209527 | HiSeq2000 | [9] |
| SRR10603720 | Bone | Healthy | GSE141614 | GSM4209528 | HiSeq2000 | [9] |
| SRR10603723 | Bone | Healthy | GSE141614 | GSM4209531 | HiSeq2000 | [9] |
| SRR19666560 | Bone | Healthy | GSE141614 | GSM6246351 | HiSeq2000 | [9] |
| SRR19666561 | Bone | Healthy | GSE141614 | GSM6246352 | HiSeq2000 | [9] |
| SRR19666562 | Bone | Healthy | GSE141614 | GSM6246353 | HiSeq2000 | [9] |
| SRR19666563 | Bone | Healthy | GSE141614 | GSM6246354 | HiSeq2000 | [9] |
| SRR19666564 | Bone | Healthy | GSE141614 | GSM6246355 | HiSeq2000 | [9] |
| SRR19666565 | Bone | Healthy | GSE141614 | GSM6246356 | HiSeq2000 | [9] |
| SRR19666566 | Bone | Healthy | GSE141614 | GSM6246357 | HiSeq2000 | [9] |
| SRR2937498 | Bone | Healthy | GSE75234 | GSM1946487 | HiSeq2000 | [10] |
| SRR2937499 | Bone | Healthy | GSE75234 | GSM1946488 | HiSeq2000 | [10] |
| SRR1175185 | Bone | Diseased | GSE55282 | GSM1333378 | HiSeq2000 | [11] |
| SRR1175186 | Bone | Diseased | GSE55282 | GSM1333379 | HiSeq2000 | [11] |
| SRR1175187 | Bone | Diseased | GSE55282 | GSM1333380 | HiSeq2000 | [11] |
| SRR1175188 | Bone | Diseased | GSE55282 | GSM1333381 | HiSeq2000 | [11] |
| SRR1175189 | Bone | Diseased | GSE55282 | GSM1333382 | HiSeq2000 | [11] |
| SRR1175190 | Bone | Diseased | GSE55282 | GSM1333383 | HiSeq2000 | [11] |
| SRR1175191 | Bone | Diseased | GSE55282 | GSM1333384 | HiSeq2000 | [11] |
| SRR1175192 | Bone | Diseased | GSE55282 | GSM1333385 | HiSeq2000 | [11] |
| SRR1175193 | Bone | Diseased | GSE55282 | GSM1333386 | HiSeq2000 | [11] |
| SRR1175194 | Bone | Diseased | GSE55282 | GSM1333387 | HiSeq2000 | [11] |
| SRR1175195 | Bone | Diseased | GSE55282 | GSM1333388 | HiSeq2000 | [11] |
| SRR1175196 | Bone | Diseased | GSE55282 | GSM1333389 | HiSeq2000 | [11] |
| SRR1175197 | Bone | Diseased | GSE55282 | GSM1333390 | HiSeq2000 | [11] |
| SRR1175198 | Bone | Diseased | GSE55282 | GSM1333391 | HiSeq2000 | [11] |
| SRR1175199 | Bone | Diseased | GSE55282 | GSM1333392 | HiSeq2000 | [11] |
| SRR1175200 | Bone | Diseased | GSE55282 | GSM1333393 | HiSeq2000 | [11] |
| SRR1175201 | Bone | Diseased | GSE55282 | GSM1333394 | HiSeq2000 | [11] |
| SRR1175202 | Bone | Diseased | GSE55282 | GSM1333395 | HiSeq2000 | [11] |
| SRR1175203 | Bone | Diseased | GSE55282 | GSM1333396 | HiSeq2000 | [11] |
| SRR1175204 | Bone | Diseased | GSE55282 | GSM1333397 | HiSeq2000 | [11] |
| SRR1175205 | Bone | Diseased | GSE55282 | GSM1333398 | HiSeq2000 | [11] |
| SRR1175206 | Bone | Diseased | GSE55282 | GSM1333399 | HiSeq2000 | [11] |
| SRR1175207 | Bone | Diseased | GSE55282 | GSM1333400 | HiSeq2000 | [11] |
| SRR1175208 | Bone | Healthy | GSE55282 | GSM1333401 | HiSeq2000 | [11] |
| SRR1175209 | Bone | Healthy | GSE55282 | GSM1333402 | HiSeq2000 | [11] |
| SRR1175210 | Bone | Healthy | GSE55282 | GSM1333403 | HiSeq2000 | [11] |
| SRR1175211 | Bone | Healthy | GSE55282 | GSM1333404 | HiSeq2000 | [11] |

| | | | | | | |
|---|---|---|---|---|---|---|
| SRR1175212 | Bone | Healthy | GSE55282 | GSM1333405 | HiSeq2000 | [11] |
| SRR1175214 | Bone | Healthy | GSE55282 | GSM1333407 | HiSeq2000 | [11] |
| SRR1175215 | Bone | Healthy | GSE55282 | GSM1333408 | HiSeq2000 | [11] |
| SRR13610768 | Ligament | Stressed | GSE165988 | GSM5059761 | HiSeq2500 | [12] |
| SRR13610769 | Ligament | Healthy | GSE165988 | GSM5059762 | HiSeq2500 | [12] |
| SRR13610770 | Ligament | Stressed | GSE165988 | GSM5059763 | HiSeq2500 | [12] |
| SRR13610771 | Ligament | Healthy | GSE165988 | GSM5059764 | HiSeq2500 | [12] |
| SRR13610772 | Ligament | Stressed | GSE165988 | GSM5059765 | HiSeq2500 | [12] |
| SRR13610773 | Ligament | Healthy | GSE165988 | GSM5059766 | HiSeq2500 | [12] |
| SRR13610774 | Ligament | Stressed | GSE165988 | GSM5059767 | HiSeq2500 | [12] |
| SRR13610775 | Ligament | Stressed | GSE165988 | GSM5059768 | HiSeq2500 | [12] |
| SRR13610776 | Ligament | Healthy | GSE165988 | GSM5059769 | HiSeq2500 | [12] |
| SRR13610777 | Ligament | Stressed | GSE165988 | GSM5059770 | HiSeq2500 | [12] |
| SRR13610778 | Ligament | Healthy | GSE165988 | GSM5059771 | HiSeq2500 | [12] |
| SRR13610779 | Ligament | Stressed | GSE165988 | GSM5059772 | HiSeq2500 | [12] |
| SRR13610780 | Ligament | Healthy | GSE165988 | GSM5059773 | HiSeq2500 | [12] |
| SRR13610781 | Ligament | Stressed | GSE165988 | GSM5059774 | HiSeq2500 | [12] |
| SRR13610782 | Ligament | Stressed | GSE165988 | GSM5059775 | HiSeq2500 | [12] |
| SRR13610783 | Ligament | Healthy | GSE165988 | GSM5059776 | HiSeq2500 | [12] |
| SRR13610784 | Ligament | Healthy | GSE165988 | GSM5059777 | HiSeq2500 | [12] |
| SRR13610785 | Ligament | Stressed | GSE165988 | GSM5059778 | HiSeq2500 | [12] |
| SRR13610786 | Ligament | Stressed | GSE165988 | GSM5059779 | HiSeq2500 | [12] |
| SRR13610787 | Ligament | Healthy | GSE165988 | GSM5059780 | HiSeq2500 | [12] |
| SRR13610788 | Ligament | Healthy | GSE165988 | GSM5059781 | HiSeq2500 | [12] |
| SRR13610789 | Ligament | Stressed | GSE165988 | GSM5059782 | HiSeq2500 | [12] |
| SRR13610790 | Ligament | Stressed | GSE165988 | GSM5059783 | HiSeq2500 | [12] |
| SRR13610791 | Ligament | Healthy | GSE165988 | GSM5059784 | HiSeq2500 | [12] |
| SRR13610792 | Ligament | Healthy | GSE165988 | GSM5059785 | HiSeq2500 | [12] |
| SRR13610793 | Ligament | Stressed | GSE165988 | GSM5059786 | HiSeq2500 | [12] |
| SRR13610794 | Ligament | Healthy | GSE165988 | GSM5059787 | HiSeq2500 | [12] |
| SRR13610795 | Ligament | Stressed | GSE165988 | GSM5059788 | HiSeq2500 | [12] |
| SRR13610796 | Ligament | Healthy | GSE165988 | GSM5059789 | HiSeq2500 | [12] |
| SRR1781814 | Ligament | Injured | GSE65469 | GSM1598127 | HiSeq2500 | [13] |
| SRR1781815 | Ligament | Injured | GSE65469 | GSM1598128 | HiSeq2500 | [13] |
| SRR1781816 | Ligament | Injured | GSE65469 | GSM1598129 | HiSeq2500 | [13] |
| SRR1781817 | Ligament | Injured | GSE65469 | GSM1598130 | HiSeq2500 | [13] |
| SRR1781818 | Ligament | Injured | GSE65469 | GSM1598131 | HiSeq2500 | [13] |
| SRR1781819 | Ligament | Injured | GSE65469 | GSM1598132 | HiSeq2500 | [13] |
| SRR1781820 | Ligament | Injured | GSE65469 | GSM1598133 | HiSeq2500 | [13] |
| SRR1781821 | Ligament | Injured | GSE65469 | GSM1598134 | HiSeq2500 | [13] |
| SRR6870939 | Bone | Treated | GSE112101 | GSM3057704 | HiSeq2000 | [14] |
| SRR6870941 | Bone | Healthy | GSE112101 | GSM3057706 | HiSeq2000 | [14] |
| SRR6870944 | Bone | Treated | GSE112101 | GSM3057709 | HiSeq2000 | [14] |
| SRR6870946 | Bone | Healthy | GSE112101 | GSM3057711 | HiSeq2000 | [14] |
| SRR6870953 | Fibroblast | Healthy | GSE112101 | GSM3057718 | HiSeq2000 | [14] |
| SRR6870955 | Fibroblast | Healthy | GSE112101 | GSM3057720 | HiSeq2000 | [14] |

| | | | | | | |
|---|---|---|---|---|---|---|
| SRR6870960 | Fibroblast | Healthy | GSE112101 | GSM3057725 | HiSeq2000 | [14] |
| SRR6870961 | Fibroblast | Healthy | GSE112101 | GSM3057726 | HiSeq2000 | [14] |
| SRR6870968 | Muscle | Healthy | GSE112101 | GSM3057733 | HiSeq2000 | [14] |
| SRR6870970 | Muscle | Healthy | GSE112101 | GSM3057735 | HiSeq2000 | [14] |
| SRR6870973 | Muscle | Healthy | GSE112101 | GSM3057738 | HiSeq2000 | [14] |
| SRR6870976 | Muscle | Healthy | GSE112101 | GSM3057741 | HiSeq2000 | [14] |
| SRR21009221 | Cartilage | Healthy | GSE210984 | GSM6443689 | HiSeq2500 | [15] |
| SRR21009222 | Cartilage | Healthy | GSE210984 | GSM6443688 | HiSeq2500 | [15] |
| SRR21009223 | Cartilage | Healthy | GSE210984 | GSM6443687 | HiSeq2500 | [15] |
| SRR21009224 | MSC | Treated | GSE210984 | GSM6443686 | HiSeq2500 | [15] |
| SRR21009225 | MSC | Treated | GSE210984 | GSM6443685 | HiSeq2500 | [15] |
| SRR21009226 | MSC | Treated | GSE210984 | GSM6443684 | HiSeq2500 | [15] |
| SRR21009227 | MSC | Treated | GSE210984 | GSM6443683 | HiSeq2500 | [15] |
| SRR21009228 | MSC | Treated | GSE210984 | GSM6443682 | HiSeq2500 | [15] |
| SRR21009229 | MSC | Treated | GSE210984 | GSM6443681 | HiSeq2500 | [15] |
| SRR21009230 | MSC | Treated | GSE210984 | GSM6443680 | HiSeq2500 | [15] |
| SRR21009231 | MSC | Treated | GSE210984 | GSM6443679 | HiSeq2500 | [15] |
| SRR21009232 | MSC | Treated | GSE210984 | GSM6443678 | HiSeq2500 | [15] |
| SRR21009233 | MSC | Healthy | GSE210984 | GSM6443677 | HiSeq2500 | [15] |
| SRR21009234 | MSC | Healthy | GSE210984 | GSM6443676 | HiSeq2500 | [15] |
| SRR21009235 | MSC | Healthy | GSE210984 | GSM6443675 | HiSeq2500 | [15] |
| SRR18550705 | MSC | Healthy | GSE199826 | GSM5988053 | HiSeq2000 | [16] |
| SRR18550706 | MSC | Healthy | GSE199826 | GSM5988052 | HiSeq2000 | [16] |
| SRR18550707 | MSC | Healthy | GSE199826 | GSM5988051 | HiSeq2000 | [16] |
| SRR18550709 | MSC | Healthy | GSE199826 | GSM5988049 | HiSeq2000 | [16] |
| SRR18550710 | MSC | Healthy | GSE199826 | GSM5988048 | HiSeq2000 | [16] |
| SRR18550711 | MSC | Healthy | GSE199826 | GSM5988047 | HiSeq2000 | [16] |
| SRR18550712 | MSC | Healthy | GSE199826 | GSM5988046 | HiSeq2000 | [16] |
| SRR18550713 | MSC | Healthy | GSE199826 | GSM5988045 | HiSeq2000 | [16] |
| SRR22560355 | MSC | Healthy | GSE220294 | GSM6797908 | HiSeq2500 | [17] |
| SRR22560356 | MSC | Treated | GSE220294 | GSM6797907 | HiSeq2500 | [17] |
| SRR22560357 | MSC | Healthy | GSE220294 | GSM6797906 | HiSeq2500 | [17] |
| SRR22560360 | MSC | Treated | GSE220294 | GSM6797903 | HiSeq2500 | [17] |
| SRR22560361 | MSC | Healthy | GSE220294 | GSM6797902 | HiSeq2500 | [17] |
| SRR14903835 | MSC | Healthy | GSE178804 | GSM5397973 | HiSeq2000 | [18] |
| SRR14903836 | MSC | Diseased | GSE178804 | GSM5397974 | HiSeq2000 | [18] |
| SRR14903839 | MSC | Healthy | GSE178804 | GSM5397977 | HiSeq2000 | [18] |
| SRR14903840 | MSC | Diseased | GSE178804 | GSM5397978 | HiSeq2000 | [18] |
| SRR14903841 | MSC | Healthy | GSE178804 | GSM5397979 | HiSeq2000 | [18] |
| SRR14903842 | MSC | Healthy | GSE178804 | GSM5397980 | HiSeq2000 | [18] |
| SRR3534841 | MSC | Healthy | GSE81478 | GSM2154690 | HiSeq2000 | [19,20] |
| SRR3534842 | MSC | Healthy | GSE81478 | GSM2154691 | HiSeq2000 | [19,20] |
| SRR3534843 | MSC | Healthy | GSE81478 | GSM2154692 | HiSeq2000 | [19,20] |
| SRR3534845 | MSC | Diseased | GSE81478 | GSM2154694 | HiSeq2000 | [19,20] |
| SRR6231603 | Muscle | Fetal | GSE106292 | GSM2835494 | HiSeq2500 | [3,4] |
| SRR6231604 | Muscle | Fetal | GSE106292 | GSM2835494 | HiSeq2500 | [3,4] |

| | | | | | | |
|---|---|---|---|---|---|---|
| SRR6231605 | Muscle | Fetal | GSE106292 | GSM2835494 | HiSeq2500 | 3,4 |
| SRR6231606 | Muscle | Fetal | GSE106292 | GSM2835494 | HiSeq2500 | 3,4 |
| SRR6231607 | Muscle | Fetal | GSE106292 | GSM2835495 | HiSeq2500 | 3,4 |
| SRR6231608 | Muscle | Fetal | GSE106292 | GSM2835495 | HiSeq2500 | 3,4 |
| SRR6231609 | Muscle | Fetal | GSE106292 | GSM2835495 | HiSeq2500 | 3,4 |
| SRR6231610 | Muscle | Fetal | GSE106292 | GSM2835495 | HiSeq2500 | 3,4 |
| SRR6231611 | Muscle | Fetal | GSE106292 | GSM2835496 | HiSeq2500 | 3,4 |
| SRR6231612 | Muscle | Fetal | GSE106292 | GSM2835496 | HiSeq2500 | 3,4 |
| SRR6231613 | Muscle | Fetal | GSE106292 | GSM2835496 | HiSeq2500 | 3,4 |
| SRR6231614 | Muscle | Fetal | GSE106292 | GSM2835496 | HiSeq2500 | 3,4 |
| SRR6231615 | Muscle | Fetal | GSE106292 | GSM2835497 | HiSeq2500 | 3,4 |
| SRR6231616 | Muscle | Fetal | GSE106292 | GSM2835497 | HiSeq2500 | 3,4 |
| SRR6231617 | Muscle | Fetal | GSE106292 | GSM2835497 | HiSeq2500 | 3,4 |
| SRR6231618 | Muscle | Fetal | GSE106292 | GSM2835497 | HiSeq2500 | 3,4 |
| SRR6337317 | Bone | Fetal | GSE106292 | GSM2871626 | HiSeq2500 | 3,4 |
| SRR6337318 | Bone | Fetal | GSE106292 | GSM2871627 | HiSeq2500 | 3,4 |
| SRR6337319 | Bone | Fetal | GSE106292 | GSM2871628 | HiSeq2500 | 3,4 |
| SRR6337320 | Ligament | Fetal | GSE106292 | GSM2871629 | HiSeq2500 | 3,4 |
| SRR6337321 | Ligament | Fetal | GSE106292 | GSM2871630 | HiSeq2500 | 3,4 |
| SRR6337322 | Ligament | Fetal | GSE106292 | GSM2871631 | HiSeq2500 | 3,4 |
| SRR6337323 | Tendon | Fetal | GSE106292 | GSM2871632 | HiSeq2500 | 3,4 |
| SRR6337324 | Tendon | Fetal | GSE106292 | GSM2871633 | HiSeq2500 | 3,4 |
| SRR6337325 | Tendon | Fetal | GSE106292 | GSM2871634 | HiSeq2500 | 3,4 |
| SRR21409916 | Bone | Healthy | GSE212631 | GSM6542014 | HiSeq2500 | 21 |
| SRR21409917 | Bone | Healthy | GSE212631 | GSM6542013 | HiSeq2500 | 21 |
| SRR21409918 | Bone | Healthy | GSE212631 | GSM6542012 | HiSeq2500 | 21 |
| SRR21409919 | Bone | Healthy | GSE212631 | GSM6542011 | HiSeq2500 | 21 |
| SRR21409920 | Bone | Healthy | GSE212631 | GSM6542010 | HiSeq2500 | 21 |
| SRR21409921 | Bone | Healthy | GSE212631 | GSM6542009 | HiSeq2500 | 21 |
| SRR21409922 | Bone | Healthy | GSE212631 | GSM6542008 | HiSeq2500 | 21 |
| SRR21409923 | Bone | Healthy | GSE212631 | GSM6542007 | HiSeq2500 | 21 |
| SRR21409924 | Bone | Healthy | GSE212631 | GSM6542006 | HiSeq2500 | 21 |
| SRR21409925 | Bone | Healthy | GSE212631 | GSM6542005 | HiSeq2500 | 21 |
| SRR21409926 | Bone | Healthy | GSE212631 | GSM6542004 | HiSeq2500 | 21 |
| SRR21409927 | Bone | Healthy | GSE212631 | GSM6542003 | HiSeq2500 | 21 |
| SRR21409928 | Bone | Healthy | GSE212631 | GSM6542002 | HiSeq2500 | 21 |
| SRR21409929 | Bone | Healthy | GSE212631 | GSM6542001 | HiSeq2500 | 21 |
| SRR21409930 | Bone | Healthy | GSE212631 | GSM6542000 | HiSeq2500 | 21 |
| SRR6870864 | Monocyte | Treated | GSE112101 | | HiSeq2000 | 14 |
| SRR6870867 | Monocyte | Healthy | GSE112101 | | HiSeq2000 | 14 |
| SRR6870871 | Monocyte | Treated | GSE112101 | | HiSeq2000 | 14 |
| SRR6870873 | Monocyte | Healthy | GSE112101 | | HiSeq2000 | 14 |
| SRR6870880 | PMN | Neutrophil treated | GSE112101 | | HiSeq2000 | 14 |
| SRR6870883 | PMN | Neutrophil healthy | GSE112101 | | HiSeq2000 | 14 |

| | | | | | | |
|---|---|---|---|---|---|---|
| SRR6870885 | PMN | Neutrophil treated | GSE112101 | | HiSeq2000 | [14] |
| SRR6870889 | PMN | Neutrophil healthy | GSE112101 | | HiSeq2000 | [14] |
| SRR6870894 | T cells | Treated | GSE112101 | GSE112101 | HiSeq2000 | [14] |
| SRR6870898 | T cells | Helper | GSE112101 | GSE112101 | HiSeq2000 | [14] |
| SRR6870900 | T cells | Treated | GSE112101 | GSE112101 | HiSeq2000 | [14] |
| SRR6870902 | T cells | Helper | GSE112101 | GSE112101 | HiSeq2000 | [14] |
| SRR6870908 | B cells | Treated | GSE112101 | GSE112101 | HiSeq2000 | [14] |
| SRR6870911 | B cells | Healthy | GSE112101 | GSE112101 | HiSeq2000 | [14] |
| SRR6870915 | B cells | Treated | GSE112101 | GSE112101 | HiSeq2000 | [14] |
| SRR6870916 | B cells | Healthy | GSE112101 | GSE112101 | HiSeq2000 | [14] |
| SRR6870923 | Endothelial Cell | Treated | GSE112101 | GSE112101 | HiSeq2000 | [14] |
| SRR6870927 | Endothelial Cell | Healthy | GSE112101 | | HiSeq2000 | [14] |
| SRR6870930 | Endothelial Cell | Treated | GSE112101 | | HiSeq2000 | [14] |
| SRR6870932 | Endothelial Cell | Healthy | GSE112101 | | HiSeq2000 | [14] |
| SRR6870983 | preadipocyte | Treated | GSE112101 | | HiSeq2000 | [14] |
| SRR6870986 | preadipocyte | Healthy | GSE112101 | | HiSeq2000 | [14] |
| SRR6870990 | preadipocyte | Treated | GSE112101 | | HiSeq2000 | [14] |
| SRR6870991 | preadipocyte | Healthy | GSE112101 | | HiSeq2000 | [14] |
| SRR16204410 | Tendon | Healthy | GSE180836 | | HiSeq2500 | [22] |
| SRR16204407 | Tendon | Healthy | GSE180836 | | HiSeq2500 | [22] |
| SRR11582313 | T cells | Mature | GSE149050 | GSM4489145 | HiSeq2500 | [23] |
| SRR11582314 | T cells | Mature | GSE149050 | GSM4489146 | HiSeq2500 | [23] |
| SRR11582316 | T cells | Mature | GSE149050 | GSM4489148 | HiSeq2500 | [23] |
| SRR11582317 | T cells | Mature | GSE149050 | GSM4489149 | HiSeq2500 | [23] |
| SRR11582318 | T cells | Mature | GSE149050 | GSM4489150 | HiSeq2500 | [23] |
| SRR11582319 | T cells | Mature | GSE149050 | GSM4489151 | HiSeq2500 | [23] |
| SRR11582320 | T cells | Mature | GSE149050 | GSM4489152 | HiSeq2500 | [23] |
| SRR11582321 | T cells | Mature | GSE149050 | GSM4489153 | HiSeq2500 | [23] |
| SRR11582322 | T cells | Mature | GSE149050 | GSM4489154 | HiSeq2500 | [23] |
| SRR11582323 | T cells | Mature | GSE149050 | GSM4489155 | HiSeq2500 | [23] |
| SRR11582324 | T cells | Mature | GSE149050 | GSM4489156 | HiSeq2500 | [23] |
| SRR11582325 | T cells | Diseased | GSE149050 | GSM4489157 | HiSeq2500 | [23] |
| SRR11582326 | T cells | Diseased | GSE149050 | GSM4489158 | HiSeq2500 | [23] |
| SRR11582327 | T cells | Diseased | GSE149050 | GSM4489159 | HiSeq2500 | [23] |
| SRR11582328 | T cells | Diseased | GSE149050 | GSM4489160 | HiSeq2500 | [23] |
| SRR11582329 | T cells | Diseased | GSE149050 | GSM4489161 | HiSeq2500 | [23] |
| SRR11582330 | T cells | Diseased | GSE149050 | GSM4489162 | HiSeq2500 | [23] |
| SRR11582331 | T cells | Diseased | GSE149050 | GSM4489163 | HiSeq2500 | [23] |
| SRR11582332 | T cells | Diseased | GSE149050 | GSM4489164 | HiSeq2500 | [23] |
| SRR11582333 | T cells | Diseased | GSE149050 | GSM4489165 | HiSeq2500 | [23] |
| SRR11582334 | T cells | Diseased | GSE149050 | GSM4489166 | HiSeq2500 | [23] |

| | | | | | | |
|---|---|---|---|---|---|---|
| SRR11582335 | T cells | Diseased | GSE149050 | GSM4489167 | HiSeq2500 | 23 |
| SRR11582336 | T cells | Diseased | GSE149050 | GSM4489168 | HiSeq2500 | 23 |
| SRR11582337 | T cells | Diseased | GSE149050 | GSM4489169 | HiSeq2500 | 23 |
| SRR11582338 | T cells | Diseased | GSE149050 | GSM4489170 | HiSeq2500 | 23 |
| SRR11582339 | T cells | Diseased | GSE149050 | GSM4489171 | HiSeq2500 | 23 |
| SRR11582340 | T cells | Diseased | GSE149050 | GSM4489172 | HiSeq2500 | 23 |
| SRR11582341 | T cells | Diseased | GSE149050 | GSM4489173 | HiSeq2500 | 23 |
| SRR11582342 | T cells | Diseased | GSE149050 | GSM4489174 | HiSeq2500 | 23 |
| SRR11582343 | T cells | Diseased | GSE149050 | GSM4489175 | HiSeq2500 | 23 |
| SRR11582344 | T cells | Diseased | GSE149050 | GSM4489176 | HiSeq2500 | 23 |
| SRR11582345 | T cells | Diseased | GSE149050 | GSM4489177 | HiSeq2500 | 23 |
| SRR11582346 | T cells | Diseased | GSE149050 | GSM4489178 | HiSeq2500 | 23 |
| SRR11582347 | T cells | Diseased | GSE149050 | GSM4489179 | HiSeq2500 | 23 |
| SRR11582348 | T cells | Diseased | GSE149050 | GSM4489180 | HiSeq2500 | 23 |
| SRR11582349 | B cells | Normal | GSE149050 | GSM4489181 | HiSeq2500 | 23 |
| SRR11582350 | B cells | Normal | GSE149050 | GSM4489182 | HiSeq2500 | 23 |
| SRR11582351 | B cells | Normal | GSE149050 | GSM4489183 | HiSeq2500 | 23 |
| SRR11582352 | B cells | Normal | GSE149050 | GSM4489184 | HiSeq2500 | 23 |
| SRR11582353 | B cells | Normal | GSE149050 | GSM4489185 | HiSeq2500 | 23 |
| SRR11582354 | B cells | Normal | GSE149050 | GSM4489186 | HiSeq2500 | 23 |
| SRR11582355 | B cells | Normal | GSE149050 | GSM4489187 | HiSeq2500 | 23 |
| SRR11582356 | B cells | Normal | GSE149050 | GSM4489188 | HiSeq2500 | 23 |
| SRR11582357 | B cells | Normal | GSE149050 | GSM4489189 | HiSeq2500 | 23 |
| SRR11582358 | B cells | Normal | GSE149050 | GSM4489190 | HiSeq2500 | 23 |
| SRR11582359 | B cells | Normal | GSE149050 | GSM4489191 | HiSeq2500 | 23 |
| SRR11582360 | B cells | Normal | GSE149050 | GSM4489192 | HiSeq2500 | 23 |
| SRR11582361 | B cells | Diseased | GSE149050 | GSM4489193 | HiSeq2500 | 23 |
| SRR11582362 | B cells | Diseased | GSE149050 | GSM4489194 | HiSeq2500 | 23 |
| SRR11582363 | B cells | Diseased | GSE149050 | GSM4489195 | HiSeq2500 | 23 |
| SRR11582364 | B cells | Diseased | GSE149050 | GSM4489196 | HiSeq2500 | 23 |
| SRR11582365 | B cells | Diseased | GSE149050 | GSM4489197 | HiSeq2500 | 23 |
| SRR11582366 | B cells | Diseased | GSE149050 | GSM4489198 | HiSeq2500 | 23 |
| SRR11582367 | B cells | Diseased | GSE149050 | GSM4489199 | HiSeq2500 | 23 |
| SRR11582368 | B cells | Diseased | GSE149050 | GSM4489200 | HiSeq2500 | 23 |
| SRR11582369 | B cells | Diseased | GSE149050 | GSM4489201 | HiSeq2500 | 23 |
| SRR11582370 | B cells | Diseased | GSE149050 | GSM4489202 | HiSeq2500 | 23 |
| SRR11582371 | B cells | Diseased | GSE149050 | GSM4489203 | HiSeq2500 | 23 |
| SRR11582372 | B cells | Diseased | GSE149050 | GSM4489204 | HiSeq2500 | 23 |
| SRR11582373 | B cells | Diseased | GSE149050 | GSM4489205 | HiSeq2500 | 23 |
| SRR11582374 | B cells | Diseased | GSE149050 | GSM4489206 | HiSeq2500 | 23 |
| SRR11582375 | B cells | Diseased | GSE149050 | GSM4489207 | HiSeq2500 | 23 |
| SRR11582376 | B cells | Diseased | GSE149050 | GSM4489208 | HiSeq2500 | 23 |
| SRR11582377 | B cells | Diseased | GSE149050 | GSM4489209 | HiSeq2500 | 23 |
| SRR11582378 | B cells | Diseased | GSE149050 | GSM4489210 | HiSeq2500 | 23 |
| SRR11582379 | B cells | Diseased | GSE149050 | GSM4489211 | HiSeq2500 | 23 |
| SRR11582380 | B cells | Diseased | GSE149050 | GSM4489212 | HiSeq2500 | 23 |

| | | | | | | |
|---|---|---|---|---|---|---|
| SRR11582381 | PMN | Normal | GSE149050 | GSM4489213 | HiSeq2500 | 23 |
| SRR11582382 | PMN | Normal | GSE149050 | GSM4489214 | HiSeq2500 | 23 |
| SRR11582383 | PMN | Normal | GSE149050 | GSM4489215 | HiSeq2500 | 23 |
| SRR11582384 | PMN | Normal | GSE149050 | GSM4489216 | HiSeq2500 | 23 |
| SRR11582385 | PMN | Normal | GSE149050 | GSM4489217 | HiSeq2500 | 23 |
| SRR11582386 | PMN | Normal | GSE149050 | GSM4489218 | HiSeq2500 | 23 |
| SRR11582387 | PMN | Normal | GSE149050 | GSM4489219 | HiSeq2500 | 23 |
| SRR11582388 | PMN | Normal | GSE149050 | GSM4489220 | HiSeq2500 | 23 |
| SRR11582389 | PMN | Normal | GSE149050 | GSM4489221 | HiSeq2500 | 23 |
| SRR11582390 | PMN | Normal | GSE149050 | GSM4489222 | HiSeq2500 | 23 |
| SRR11582391 | PMN | Normal | GSE149050 | GSM4489223 | HiSeq2500 | 23 |
| SRR11582392 | PMN | Normal | GSE149050 | GSM4489224 | HiSeq2500 | 23 |
| SRR11582393 | PMN | Diseased | GSE149050 | GSM4489225 | HiSeq2500 | 23 |
| SRR11582394 | PMN | Diseased | GSE149050 | GSM4489226 | HiSeq2500 | 23 |
| SRR11582395 | PMN | Diseased | GSE149050 | GSM4489227 | HiSeq2500 | 23 |
| SRR11582396 | PMN | Diseased | GSE149050 | GSM4489228 | HiSeq2500 | 23 |
| SRR11582397 | PMN | Diseased | GSE149050 | GSM4489229 | HiSeq2500 | 23 |
| SRR11582398 | PMN | Diseased | GSE149050 | GSM4489230 | HiSeq2500 | 23 |
| SRR11582399 | PMN | Diseased | GSE149050 | GSM4489231 | HiSeq2500 | 23 |
| SRR11582400 | PMN | Diseased | GSE149050 | GSM4489232 | HiSeq2500 | 23 |
| SRR11582401 | PMN | Diseased | GSE149050 | GSM4489233 | HiSeq2500 | 23 |
| SRR11582402 | PMN | Diseased | GSE149050 | GSM4489234 | HiSeq2500 | 23 |
| SRR11582403 | PMN | Diseased | GSE149050 | GSM4489235 | HiSeq2500 | 23 |
| SRR11582404 | PMN | Diseased | GSE149050 | GSM4489236 | HiSeq2500 | 23 |
| SRR11582405 | PMN | Diseased | GSE149050 | GSM4489237 | HiSeq2500 | 23 |
| SRR11582406 | PMN | Diseased | GSE149050 | GSM4489238 | HiSeq2500 | 23 |
| SRR11582407 | PMN | Diseased | GSE149050 | GSM4489239 | HiSeq2500 | 23 |
| SRR11582408 | PMN | Diseased | GSE149050 | GSM4489240 | HiSeq2500 | 23 |
| SRR11582409 | PMN | Diseased | GSE149050 | GSM4489241 | HiSeq2500 | 23 |
| SRR11582410 | PMN | Diseased | GSE149050 | GSM4489242 | HiSeq2500 | 23 |
| SRR11582411 | PMN | Diseased | GSE149050 | GSM4489243 | HiSeq2500 | 23 |
| SRR11582412 | PMN | Diseased | GSE149050 | GSM4489244 | HiSeq2500 | 23 |
| SRR11582413 | PMN | Diseased | GSE149050 | GSM4489245 | HiSeq2500 | 23 |
| SRR11582414 | PMN | Diseased | GSE149050 | GSM4489246 | HiSeq2500 | 23 |
| SRR11582415 | PMN | Diseased | GSE149050 | GSM4489247 | HiSeq2500 | 23 |
| SRR11582416 | PMN | Diseased | GSE149050 | GSM4489248 | HiSeq2500 | 23 |
| SRR11582417 | Dendritic cells | cDC | GSE149050 | GSM4489249 | HiSeq2500 | 23 |
| SRR11582418 | Dendritic cells | cDC | GSE149050 | GSM4489250 | HiSeq2500 | 23 |
| SRR11582419 | Dendritic cells | cDC | GSE149050 | GSM4489251 | HiSeq2500 | 23 |
| SRR11582420 | Dendritic cells | cDC | GSE149050 | GSM4489252 | HiSeq2500 | 23 |
| SRR11582421 | Dendritic cells | cDC | GSE149050 | GSM4489253 | HiSeq2500 | 23 |

| | | | | | | |
|---|---|---|---|---|---|---|
| SRR11582422 | Dendritic cells | cDC | GSE149050 | GSM4489254 | HiSeq2500 | 23 |
| SRR11582423 | Dendritic cells | cDC | GSE149050 | GSM4489255 | HiSeq2500 | 23 |
| SRR11582424 | Dendritic cells | cDC | GSE149050 | GSM4489256 | HiSeq2500 | 23 |
| SRR11582425 | Dendritic cells | cDC | GSE149050 | GSM4489257 | HiSeq2500 | 23 |
| SRR11582426 | Dendritic cells | cDC | GSE149050 | GSM4489258 | HiSeq2500 | 23 |
| SRR11582427 | Dendritic cells | cDC | GSE149050 | GSM4489259 | HiSeq2500 | 23 |
| SRR11582428 | Dendritic cells | cDC | GSE149050 | GSM4489260 | HiSeq2500 | 23 |
| SRR11582429 | Dendritic cells | cDC diseased | GSE149050 | GSM4489261 | HiSeq2500 | 23 |
| SRR11582430 | Dendritic cells | cDC diseased | GSE149050 | GSM4489262 | HiSeq2500 | 23 |
| SRR11582431 | Dendritic cells | cDC diseased | GSE149050 | GSM4489263 | HiSeq2500 | 23 |
| SRR11582432 | Dendritic cells | cDC diseased | GSE149050 | GSM4489264 | HiSeq2500 | 23 |
| SRR11582433 | Dendritic cells | cDC diseased | GSE149050 | GSM4489265 | HiSeq2500 | 23 |
| SRR11582434 | Dendritic cells | cDC diseased | GSE149050 | GSM4489266 | HiSeq2500 | 23 |
| SRR11582435 | Dendritic cells | cDC diseased | GSE149050 | GSM4489267 | HiSeq2500 | 23 |
| SRR11582436 | Dendritic cells | cDC diseased | GSE149050 | GSM4489268 | HiSeq2500 | 23 |
| SRR11582437 | Dendritic cells | cDC diseased | GSE149050 | GSM4489269 | HiSeq2500 | 23 |
| SRR11582438 | Dendritic cells | cDC diseased | GSE149050 | GSM4489270 | HiSeq2500 | 23 |
| SRR11582439 | Dendritic cells | cDC diseased | GSE149050 | GSM4489271 | HiSeq2500 | 23 |
| SRR11582440 | Dendritic cells | cDC diseased | GSE149050 | GSM4489272 | HiSeq2500 | 23 |
| SRR11582441 | Dendritic cells | cDC diseased | GSE149050 | GSM4489273 | HiSeq2500 | 23 |
| SRR11582442 | Dendritic cells | cDC diseased | GSE149050 | GSM4489274 | HiSeq2500 | 23 |
| SRR11582443 | Dendritic cells | cDC diseased | GSE149050 | GSM4489275 | HiSeq2500 | 23 |
| SRR11582444 | Dendritic cells | cDC diseased | GSE149050 | GSM4489276 | HiSeq2500 | 23 |
| SRR11582445 | Dendritic cells | cDC diseased | GSE149050 | GSM4489277 | HiSeq2500 | 23 |

| | | | | | | |
|---|---|---|---|---|---|---|
| SRR11582446 | Dendritic cells | cDC diseased | GSE149050 | GSM4489278 | HiSeq2500 | 23 |
| SRR11582447 | Dendritic cells | cDC diseased | GSE149050 | GSM4489279 | HiSeq2500 | 23 |
| SRR11582448 | Dendritic cells | cDC diseased | GSE149050 | GSM4489280 | HiSeq2500 | 23 |
| SRR11582449 | Dendritic cells | pDC | GSE149050 | GSM4489281 | HiSeq2500 | 23 |
| SRR11582450 | Dendritic cells | pDC | GSE149050 | GSM4489282 | HiSeq2500 | 23 |
| SRR11582451 | Dendritic cells | pDC | GSE149050 | GSM4489283 | HiSeq2500 | 23 |
| SRR11582452 | Dendritic cells | pDC | GSE149050 | GSM4489284 | HiSeq2500 | 23 |
| SRR11582453 | Dendritic cells | pDC | GSE149050 | GSM4489285 | HiSeq2500 | 23 |
| SRR11582454 | Dendritic cells | pDC | GSE149050 | GSM4489286 | HiSeq2500 | 23 |
| SRR11582455 | Dendritic cells | pDC | GSE149050 | GSM4489287 | HiSeq2500 | 23 |
| SRR11582456 | Dendritic cells | pDC | GSE149050 | GSM4489288 | HiSeq2500 | 23 |
| SRR11582457 | Dendritic cells | pDC | GSE149050 | GSM4489289 | HiSeq2500 | 23 |
| SRR11582458 | Dendritic cells | pDC | GSE149050 | GSM4489290 | HiSeq2500 | 23 |
| SRR11582459 | Dendritic cells | pDC | GSE149050 | GSM4489291 | HiSeq2500 | 23 |
| SRR11582460 | Dendritic cells | pDC disease | GSE149050 | GSM4489292 | HiSeq2500 | 23 |
| SRR11582461 | Dendritic cells | pDC disease | GSE149050 | GSM4489293 | HiSeq2500 | 23 |
| SRR11582462 | Dendritic cells | pDC disease | GSE149050 | GSM4489294 | HiSeq2500 | 23 |
| SRR11582463 | Dendritic cells | pDC disease | GSE149050 | GSM4489295 | HiSeq2500 | 23 |
| SRR11582464 | Dendritic cells | pDC disease | GSE149050 | GSM4489296 | HiSeq2500 | 23 |
| SRR11582465 | Dendritic cells | pDC disease | GSE149050 | GSM4489297 | HiSeq2500 | 23 |
| SRR11582466 | Dendritic cells | pDC disease | GSE149050 | GSM4489298 | HiSeq2500 | 23 |
| SRR11582467 | Dendritic cells | pDC disease | GSE149050 | GSM4489299 | HiSeq2500 | 23 |
| SRR11582468 | Dendritic cells | pDC disease | GSE149050 | GSM4489300 | HiSeq2500 | 23 |
| SRR11582469 | Dendritic cells | pDC disease | GSE149050 | GSM4489301 | HiSeq2500 | 23 |

| | | | | | | |
|---|---|---|---|---|---|---|
| SRR11582470 | Dendritic cells | pDC disease | GSE149050 | GSM4489302 | HiSeq2500 | 23 |
| SRR11582471 | Dendritic cells | pDC disease | GSE149050 | GSM4489303 | HiSeq2500 | 23 |
| SRR11582472 | Dendritic cells | pDC disease | GSE149050 | GSM4489304 | HiSeq2500 | 23 |
| SRR11582473 | Dendritic cells | pDC disease | GSE149050 | GSM4489305 | HiSeq2500 | 23 |
| SRR11582474 | Dendritic cells | pDC disease | GSE149050 | GSM4489306 | HiSeq2500 | 23 |
| SRR11582475 | Dendritic cells | pDC disease | GSE149050 | GSM4489307 | HiSeq2500 | 23 |
| SRR11582476 | Dendritic cells | pDC disease | GSE149050 | GSM4489308 | HiSeq2500 | 23 |
| SRR11582477 | Dendritic cells | pDC disease | GSE149050 | GSM4489309 | HiSeq2500 | 23 |
| SRR11582478 | Dendritic cells | pDC disease | GSE149050 | GSM4489310 | HiSeq2500 | 23 |
| SRR11582479 | Dendritic cells | pDC disease | GSE149050 | GSM4489311 | HiSeq2500 | 23 |
| SRR11582480 | Dendritic cells | pDC disease | GSE149050 | GSM4489312 | HiSeq2500 | 23 |
| SRR25254033 | T cells | Diseased | GSE237218 | GSM7597340 | HiSeq2000 | 24 |
| SRR25254034 | CD4p | Memory | GSE237218 | GSM7597339 | HiSeq2000 | 24 |
| SRR25254035 | T cells | Diseased | GSE237218 | GSM7597338 | HiSeq2000 | 24 |
| SRR25254036 | B cells | Diseased | GSE237218 | GSM7597337 | HiSeq2000 | 24 |
| SRR25254037 | B cells | Normal | GSE237218 | GSM7597336 | HiSeq2000 | 24 |
| SRR25254038 | T cells | Diseased | GSE237218 | GSM7597335 | HiSeq2000 | 24 |
| SRR25254039 | T cells | Diseased | GSE237218 | GSM7597334 | HiSeq2000 | 24 |
| SRR25254040 | T cells | Diseased | GSE237218 | GSM7597333 | HiSeq2000 | 24 |
| SRR25254041 | CD4p | Memory | GSE237218 | GSM7597332 | HiSeq2000 | 24 |
| SRR25254042 | CD4p | Memory | GSE237218 | GSM7597331 | HiSeq2000 | 24 |
| SRR25254043 | CD4p | Memory | GSE237218 | GSM7597330 | HiSeq2000 | 24 |
| SRR25254044 | T cells | Diseased | GSE237218 | GSM7597329 | HiSeq2000 | 24 |
| SRR25254045 | B cells | Normal | GSE237218 | GSM7597328 | HiSeq2000 | 24 |
| SRR25254046 | B cells | Normal | GSE237218 | GSM7597327 | HiSeq2000 | 24 |
| SRR25254047 | B cells | Normal | GSE237218 | GSM7597326 | HiSeq2000 | 24 |
| SRR25254048 | B cells | Diseased | GSE237218 | GSM7597325 | HiSeq2000 | 24 |
| SRR25254049 | B cells | Normal | GSE237218 | GSM7597324 | HiSeq2000 | 24 |
| SRR25254050 | B cells | Normal | GSE237218 | GSM7597323 | HiSeq2000 | 24 |
| SRR25254051 | B cells | Normal | GSE237218 | GSM7597322 | HiSeq2000 | 24 |
| SRR25254052 | B cells | Normal | GSE237218 | GSM7597321 | HiSeq2000 | 24 |
| SRR25254053 | B cells | Diseased | GSE237218 | GSM7597320 | HiSeq2000 | 24 |
| SRR25254054 | B cells | Diseased | GSE237218 | GSM7597319 | HiSeq2000 | 24 |
| SRR25254055 | B cells | Normal | GSE237218 | GSM7597318 | HiSeq2000 | 24 |
| SRR25254056 | B cells | Diseased | GSE237218 | GSM7596981 | HiSeq2000 | 24 |
| SRR25254057 | B cells | Normal | GSE237218 | GSM7596980 | HiSeq2000 | 24 |

| | | | | | | |
|---|---|---|---|---|---|---|
| SRR25254058 | B cells | Diseased | GSE237218 | GSM7596979 | HiSeq2000 | [24] |
| SRR25253617 | T cells | Diseased | GSE237218 | GSM7597125 | HiSeq2000 | [24] |
| SRR25253618 | T cells | Memory | GSE237218 | GSM7597124 | HiSeq2000 | [24] |
| SRR25253619 | T cells | Memory | GSE237218 | GSM7597123 | HiSeq2000 | [24] |
| SRR25253620 | T cells | Diseased | GSE237218 | GSM7597122 | HiSeq2000 | [24] |
| SRR25253621 | T cells | Diseased | GSE237218 | GSM7597121 | HiSeq2000 | [24] |
| SRR25253622 | T cells | Memory | GSE237218 | GSM7597120 | HiSeq2000 | [24] |
| SRR25253623 | T cells | Memory | GSE237218 | GSM7597119 | HiSeq2000 | [24] |
| SRR25253624 | T cells | Diseased | GSE237218 | GSM7597118 | HiSeq2000 | [24] |
| SRR25253625 | T cells | Diseased | GSE237218 | GSM7597117 | HiSeq2000 | [24] |
| SRR25253626 | T cells | Memory | GSE237218 | GSM7597116 | HiSeq2000 | [24] |
| SRR25253627 | T cells | Memory | GSE237218 | GSM7597115 | HiSeq2000 | [24] |
| SRR25253628 | T cells | Diseased | GSE237218 | GSM7597114 | HiSeq2000 | [24] |
| SRR25253629 | T cells | Diseased | GSE237218 | GSM7597113 | HiSeq2000 | [24] |
| SRR25253630 | T cells | Memory | GSE237218 | GSM7597112 | HiSeq2000 | [24] |
| SRR25253631 | T cells | Memory | GSE237218 | GSM7597111 | HiSeq2000 | [24] |
| SRR25253632 | T cells | Diseased | GSE237218 | GSM7597110 | HiSeq2000 | [24] |
| SRR25253633 | T cells | Memory | GSE237218 | GSM7597109 | HiSeq2000 | [24] |
| SRR25253634 | T cells | Diseased | GSE237218 | GSM7597108 | HiSeq2000 | [24] |
| SRR25253635 | T cells | Memory | GSE237218 | GSM7597107 | HiSeq2000 | [24] |
| SRR25253636 | T cells | Diseased | GSE237218 | GSM7597106 | HiSeq2000 | [24] |
| SRR25253637 | T cells | Diseased | GSE237218 | GSM7597105 | HiSeq2000 | [24] |
| SRR25253638 | T cells | Memory | GSE237218 | GSM7597104 | HiSeq2000 | [24] |
| SRR25253639 | T cells | Memory | GSE237218 | GSM7597103 | HiSeq2000 | [24] |
| SRR25253640 | T cells | Diseased | GSE237218 | GSM7597102 | HiSeq2000 | [24] |
| SRR25253641 | B cells | Normal | GSE237218 | GSM7597245 | HiSeq2000 | [24] |
| SRR25253642 | T cells | Regulatory | GSE237218 | GSM7597244 | HiSeq2000 | [24] |
| SRR25253643 | B cells | Diseased | GSE237218 | GSM7597243 | HiSeq2000 | [24] |
| SRR25253644 | B cells | Normal | GSE237218 | GSM7597242 | HiSeq2000 | [24] |
| SRR25253645 | B cells | Diseased | GSE237218 | GSM7597241 | HiSeq2000 | [24] |
| SRR25253646 | B cells | Normal | GSE237218 | GSM7597240 | HiSeq2000 | [24] |
| SRR25253647 | B cells | Diseased | GSE237218 | GSM7597239 | HiSeq2000 | [24] |
| SRR25253648 | B cells | Diseased | GSE237218 | GSM7597238 | HiSeq2000 | [24] |
| SRR25253649 | B cells | Normal | GSE237218 | GSM7597237 | HiSeq2000 | [24] |
| SRR25253650 | B cells | Diseased | GSE237218 | GSM7597236 | HiSeq2000 | [24] |
| SRR25253651 | B cells | Diseased | GSE237218 | GSM7597235 | HiSeq2000 | [24] |
| SRR25253652 | T cells | Diseased | GSE237218 | GSM7597234 | HiSeq2000 | [24] |
| SRR25253653 | T cells | Diseased | GSE237218 | GSM7597233 | HiSeq2000 | [24] |
| SRR25253654 | T cells | Regulatory | GSE237218 | GSM7597232 | HiSeq2000 | [24] |
| SRR25253655 | T cells | Memory | GSE237218 | GSM7597231 | HiSeq2000 | [24] |
| SRR25253656 | T cells | Memory | GSE237218 | GSM7597230 | HiSeq2000 | [24] |
| SRR25253657 | T cells | Memory | GSE237218 | GSM7597229 | HiSeq2000 | [24] |
| SRR25253658 | T cells | Memory | GSE237218 | GSM7597228 | HiSeq2000 | [24] |
| SRR25253659 | T cells | Diseased | GSE237218 | GSM7597227 | HiSeq2000 | [24] |
| SRR25253660 | T cells | Diseased | GSE237218 | GSM7597226 | HiSeq2000 | [24] |
| SRR25253661 | T cells | Regulatory | GSE237218 | GSM7597225 | HiSeq2000 | [24] |

| | | | | | | |
|---|---|---|---|---|---|---|
| SRR25253662 | T cells | Regulatory | GSE237218 | GSM7597224 | HiSeq2000 | [24] |
| SRR25253663 | T cells | Diseased | GSE237218 | GSM7597223 | HiSeq2000 | [24] |
| SRR25253664 | T cells | Regulatory | GSE237218 | GSM7597222 | HiSeq2000 | [24] |
| SRR25253665 | T cells | Diseased | GSE237218 | GSM7597173 | HiSeq2000 | [24] |
| SRR25253666 | T cells | Memory | GSE237218 | GSM7597172 | HiSeq2000 | [24] |
| SRR25253667 | T cells | Memory | GSE237218 | GSM7597171 | HiSeq2000 | [24] |
| SRR25253668 | T cells | Diseased | GSE237218 | GSM7597170 | HiSeq2000 | [24] |
| SRR25253669 | T cells | Memory | GSE237218 | GSM7597169 | HiSeq2000 | [24] |
| SRR25253670 | T cells | Diseased | GSE237218 | GSM7597168 | HiSeq2000 | [24] |
| SRR25253671 | T cells | Diseased | GSE237218 | GSM7597167 | HiSeq2000 | [24] |
| SRR25253672 | T cells | Regulatory | GSE237218 | GSM7597166 | HiSeq2000 | [24] |
| SRR25253673 | T cells | Regulatory | GSE237218 | GSM7597165 | HiSeq2000 | [24] |
| SRR25253674 | T cells | Diseased | GSE237218 | GSM7597164 | HiSeq2000 | [24] |
| SRR25253675 | T cells | Diseased | GSE237218 | GSM7597163 | HiSeq2000 | [24] |
| SRR25253676 | T cells | Diseased | GSE237218 | GSM7597162 | HiSeq2000 | [24] |
| SRR25253677 | T cells | Diseased | GSE237218 | GSM7597161 | HiSeq2000 | [24] |
| SRR25253678 | T cells | Diseased | GSE237218 | GSM7597160 | HiSeq2000 | [24] |
| SRR25253679 | T cells | Memory | GSE237218 | GSM7597159 | HiSeq2000 | [24] |
| SRR25253680 | T cells | Memory | GSE237218 | GSM7597158 | HiSeq2000 | [24] |
| SRR25253681 | T cells | Memory | GSE237218 | GSM7597157 | HiSeq2000 | [24] |
| SRR25253682 | T cells | Diseased | GSE237218 | GSM7597156 | HiSeq2000 | [24] |
| SRR25253683 | T cells | Diseased | GSE237218 | GSM7597155 | HiSeq2000 | [24] |
| SRR25253684 | T cells | Regulatory | GSE237218 | GSM7597154 | HiSeq2000 | [24] |
| SRR25253685 | T cells | Diseased | GSE237218 | GSM7597153 | HiSeq2000 | [24] |
| SRR25253686 | T cells | Diseased | GSE237218 | GSM7597152 | HiSeq2000 | [24] |
| SRR25253687 | T cells | Diseased | GSE237218 | GSM7597151 | HiSeq2000 | [24] |
| SRR25253688 | T cells | Regulatory | GSE237218 | GSM7597150 | HiSeq2000 | [24] |
| SRR25253689 | T cells | Memory | GSE237218 | GSM7597413 | HiSeq2000 | [24] |
| SRR25253690 | B cells | Normal | GSE237218 | GSM7597412 | HiSeq2000 | [24] |
| SRR25253691 | T cells | Memory | GSE237218 | GSM7597411 | HiSeq2000 | [24] |
| SRR25253692 | T cells | Memory | GSE237218 | GSM7597410 | HiSeq2000 | [24] |
| SRR25253693 | T cells | Diseased | GSE237218 | GSM7597409 | HiSeq2000 | [24] |
| SRR25253694 | T cells | Diseased | GSE237218 | GSM7597408 | HiSeq2000 | [24] |
| SRR25253695 | T cells | Diseased | GSE237218 | GSM7597407 | HiSeq2000 | [24] |
| SRR25253696 | T cells | Diseased | GSE237218 | GSM7597406 | HiSeq2000 | [24] |
| SRR25253697 | B cells | Normal | GSE237218 | GSM7597405 | HiSeq2000 | [24] |
| SRR25253698 | B cells | Normal | GSE237218 | GSM7597404 | HiSeq2000 | [24] |
| SRR25253699 | B cells | Normal | GSE237218 | GSM7597403 | HiSeq2000 | [24] |
| SRR25253700 | B cells | Diseased | GSE237218 | GSM7597402 | HiSeq2000 | [24] |
| SRR25253701 | T cells | Diseased | GSE237218 | GSM7597401 | HiSeq2000 | [24] |
| SRR25253702 | T cells | Diseased | GSE237218 | GSM7597400 | HiSeq2000 | [24] |
| SRR25253703 | T cells | Regulatory | GSE237218 | GSM7597399 | HiSeq2000 | [24] |
| SRR25253704 | T cells | Diseased | GSE237218 | GSM7597398 | HiSeq2000 | [24] |
| SRR25253705 | T cells | Diseased | GSE237218 | GSM7597397 | HiSeq2000 | [24] |
| SRR25253706 | T cells | Memory | GSE237218 | GSM7597396 | HiSeq2000 | [24] |
| SRR25253707 | B cells | Normal | GSE237218 | GSM7597395 | HiSeq2000 | [24] |

| | | | | | | |
|---|---|---|---|---|---|---|
| SRR25253708 | B cells | Diseased | GSE237218 | GSM7597394 | HiSeq2000 | 24 |
| SRR25253709 | B cells | Diseased | GSE237218 | GSM7597393 | HiSeq2000 | 24 |
| SRR25253710 | T cells | Diseased | GSE237218 | GSM7597392 | HiSeq2000 | 24 |
| SRR25253711 | T cells | Diseased | GSE237218 | GSM7597391 | HiSeq2000 | 24 |
| SRR25253712 | T cells | Diseased | GSE237218 | GSM7597390 | HiSeq2000 | 24 |
| SRR25253713 | T cells | Diseased | GSE237218 | GSM7597077 | HiSeq2000 | 24 |
| SRR25253714 | T cells | Diseased | GSE237218 | GSM7597076 | HiSeq2000 | 24 |
| SRR25253715 | T cells | Diseased | GSE237218 | GSM7597075 | HiSeq2000 | 24 |
| SRR25253716 | T cells | Diseased | GSE237218 | GSM7597074 | HiSeq2000 | 24 |
| SRR25253717 | T cells | Diseased | GSE237218 | GSM7597073 | HiSeq2000 | 24 |
| SRR25253718 | T cells | Diseased | GSE237218 | GSM7597072 | HiSeq2000 | 24 |
| SRR25253719 | T cells | Memory | GSE237218 | GSM7597071 | HiSeq2000 | 24 |
| SRR25253720 | T cells | Memory | GSE237218 | GSM7597070 | HiSeq2000 | 24 |
| SRR25253721 | T cells | Diseased | GSE237218 | GSM7597069 | HiSeq2000 | 24 |
| SRR25253722 | T cells | Diseased | GSE237218 | GSM7597068 | HiSeq2000 | 24 |
| SRR25253723 | T cells | Memory | GSE237218 | GSM7597067 | HiSeq2000 | 24 |
| SRR25253724 | T cells | Memory | GSE237218 | GSM7597066 | HiSeq2000 | 24 |
| SRR25253725 | T cells | Diseased | GSE237218 | GSM7597065 | HiSeq2000 | 24 |
| SRR25253726 | T cells | Diseased | GSE237218 | GSM7597064 | HiSeq2000 | 24 |
| SRR25253727 | T cells | Diseased | GSE237218 | GSM7597063 | HiSeq2000 | 24 |
| SRR25253728 | T cells | Memory | GSE237218 | GSM7597062 | HiSeq2000 | 24 |
| SRR25253729 | T cells | Diseased | GSE237218 | GSM7597061 | HiSeq2000 | 24 |
| SRR25253730 | T cells | Memory | GSE237218 | GSM7597060 | HiSeq2000 | 24 |
| SRR25253731 | T cells | Diseased | GSE237218 | GSM7597059 | HiSeq2000 | 24 |
| SRR25253732 | T cells | Memory | GSE237218 | GSM7597058 | HiSeq2000 | 24 |
| SRR25253733 | T cells | Memory | GSE237218 | GSM7597057 | HiSeq2000 | 24 |
| SRR25253734 | T cells | Memory | GSE237218 | GSM7597056 | HiSeq2000 | 24 |
| SRR25253735 | T cells | Diseased | GSE237218 | GSM7597055 | HiSeq2000 | 24 |
| SRR25253736 | T cells | Memory | GSE237218 | GSM7597054 | HiSeq2000 | 24 |
| SRR25253737 | B cells | Diseased | GSE237218 | GSM7597269 | HiSeq2000 | 24 |
| SRR25253738 | T cells | Regulatory | GSE237218 | GSM7597268 | HiSeq2000 | 24 |
| SRR25253739 | B cells | Normal | GSE237218 | GSM7597267 | HiSeq2000 | 24 |
| SRR25253740 | B cells | Normal | GSE237218 | GSM7597266 | HiSeq2000 | 24 |
| SRR25253741 | B cells | Normal | GSE237218 | GSM7597265 | HiSeq2000 | 24 |
| SRR25253742 | B cells | Diseased | GSE237218 | GSM7597264 | HiSeq2000 | 24 |
| SRR25253743 | B cells | Diseased | GSE237218 | GSM7597263 | HiSeq2000 | 24 |
| SRR25253744 | B cells | Normal | GSE237218 | GSM7597262 | HiSeq2000 | 24 |
| SRR25253745 | B cells | Normal | GSE237218 | GSM7597261 | HiSeq2000 | 24 |
| SRR25253746 | B cells | Diseased | GSE237218 | GSM7597260 | HiSeq2000 | 24 |
| SRR25253747 | B cells | Normal | GSE237218 | GSM7597259 | HiSeq2000 | 24 |
| SRR25253748 | B cells | Normal | GSE237218 | GSM7597258 | HiSeq2000 | 24 |
| SRR25253749 | B cells | Normal | GSE237218 | GSM7597257 | HiSeq2000 | 24 |
| SRR25253750 | T cells | Diseased | GSE237218 | GSM7597256 | HiSeq2000 | 24 |
| SRR25253751 | B cells | Diseased | GSE237218 | GSM7597255 | HiSeq2000 | 24 |
| SRR25253752 | B cells | Normal | GSE237218 | GSM7597254 | HiSeq2000 | 24 |
| SRR25253753 | B cells | Diseased | GSE237218 | GSM7597253 | HiSeq2000 | 24 |

| | | | | | | |
|---|---|---|---|---|---|---|
| SRR25253754 | B cells | Normal | GSE237218 | GSM7597252 | HiSeq2000 | [24] |
| SRR25253755 | B cells | Normal | GSE237218 | GSM7597251 | HiSeq2000 | [24] |
| SRR25253756 | B cells | Normal | GSE237218 | GSM7597250 | HiSeq2000 | [24] |
| SRR25253757 | B cells | Normal | GSE237218 | GSM7597249 | HiSeq2000 | [24] |
| SRR25253758 | B cells | Diseased | GSE237218 | GSM7597248 | HiSeq2000 | [24] |
| SRR25253759 | B cells | Normal | GSE237218 | GSM7597247 | HiSeq2000 | [24] |
| SRR25253760 | B cells | Diseased | GSE237218 | GSM7597246 | HiSeq2000 | [24] |
| SRR25253761 | T cells | Diseased | GSE237218 | GSM7597197 | HiSeq2000 | [24] |
| SRR25253762 | T cells | Diseased | GSE237218 | GSM7597196 | HiSeq2000 | [24] |
| SRR25253763 | T cells | Diseased | GSE237218 | GSM7597195 | HiSeq2000 | [24] |
| SRR25253764 | T cells | Memory | GSE237218 | GSM7597194 | HiSeq2000 | [24] |
| SRR25253765 | T cells | Memory | GSE237218 | GSM7597193 | HiSeq2000 | [24] |
| SRR25253766 | T cells | Memory | GSE237218 | GSM7597192 | HiSeq2000 | [24] |
| SRR25253767 | T cells | Diseased | GSE237218 | GSM7597191 | HiSeq2000 | [24] |
| SRR25253768 | T cells | Diseased | GSE237218 | GSM7597190 | HiSeq2000 | [24] |
| SRR25253769 | T cells | Diseased | GSE237218 | GSM7597189 | HiSeq2000 | [24] |
| SRR25253770 | T cells | Diseased | GSE237218 | GSM7597188 | HiSeq2000 | [24] |
| SRR25253771 | T cells | Diseased | GSE237218 | GSM7597187 | HiSeq2000 | [24] |
| SRR25253772 | T cells | Diseased | GSE237218 | GSM7597186 | HiSeq2000 | [24] |
| SRR25253773 | T cells | Regulatory | GSE237218 | GSM7597185 | HiSeq2000 | [24] |
| SRR25253774 | T cells | Diseased | GSE237218 | GSM7597184 | HiSeq2000 | [24] |
| SRR25253775 | T cells | Memory | GSE237218 | GSM7597183 | HiSeq2000 | [24] |
| SRR25253776 | T cells | Memory | GSE237218 | GSM7597182 | HiSeq2000 | [24] |
| SRR25253777 | T cells | Memory | GSE237218 | GSM7597181 | HiSeq2000 | [24] |
| SRR25253778 | T cells | Diseased | GSE237218 | GSM7597180 | HiSeq2000 | [24] |
| SRR25253779 | T cells | Diseased | GSE237218 | GSM7597179 | HiSeq2000 | [24] |
| SRR25253780 | T cells | Regulatory | GSE237218 | GSM7597178 | HiSeq2000 | [24] |
| SRR25253781 | T cells | Diseased | GSE237218 | GSM7597177 | HiSeq2000 | [24] |
| SRR25253782 | T cells | Diseased | GSE237218 | GSM7597176 | HiSeq2000 | [24] |
| SRR25253783 | T cells | Diseased | GSE237218 | GSM7597175 | HiSeq2000 | [24] |
| SRR25253784 | T cells | Diseased | GSE237218 | GSM7597174 | HiSeq2000 | [24] |
| SRR25253785 | T cells | Diseased | GSE237218 | GSM7597420 | HiSeq2000 | [24] |
| SRR25253786 | T cells | Memory | GSE237218 | GSM7597419 | HiSeq2000 | [24] |
| SRR25253787 | T cells | Memory | GSE237218 | GSM7597418 | HiSeq2000 | [24] |
| SRR25253919 | B cells | Normal | GSE237218 | GSM7597382 | HiSeq2000 | [24] |
| SRR25253920 | B cells | Normal | GSE237218 | GSM7597381 | HiSeq2000 | [24] |
| SRR25253921 | T cells | Regulatory | GSE237218 | GSM7597380 | HiSeq2000 | [24] |
| SRR25253922 | T cells | Regulatory | GSE237218 | GSM7597379 | HiSeq2000 | [24] |
| SRR25253923 | T cells | Memory | GSE237218 | GSM7597378 | HiSeq2000 | [24] |
| SRR25253924 | T cells | Diseased | GSE237218 | GSM7597377 | HiSeq2000 | [24] |
| SRR25253925 | T cells | Memory | GSE237218 | GSM7597376 | HiSeq2000 | [24] |
| SRR25253926 | T cells | Diseased | GSE237218 | GSM7597375 | HiSeq2000 | [24] |
| SRR25253927 | T cells | Memory | GSE237218 | GSM7597374 | HiSeq2000 | [24] |
| SRR25253928 | T cells | Memory | GSE237218 | GSM7597373 | HiSeq2000 | [24] |
| SRR25253929 | T cells | Memory | GSE237218 | GSM7597372 | HiSeq2000 | [24] |
| SRR25253930 | B cells | Diseased | GSE237218 | GSM7597371 | HiSeq2000 | [24] |

| | | | | | | |
|---|---|---|---|---|---|---|
| SRR25253931 | B cells | Normal | GSE237218 | GSM7597370 | HiSeq2000 | [24] |
| SRR25253932 | T cells | Regulatory | GSE237218 | GSM7597369 | HiSeq2000 | [24] |
| SRR25253933 | T cells | Regulatory | GSE237218 | GSM7597368 | HiSeq2000 | [24] |
| SRR25253934 | T cells | Diseased | GSE237218 | GSM7597367 | HiSeq2000 | [24] |
| SRR25253935 | T cells | Memory | GSE237218 | GSM7597366 | HiSeq2000 | [24] |
| SRR25253936 | T cells | Diseased | GSE237218 | GSM7597101 | HiSeq2000 | [24] |
| SRR25253937 | T cells | Memory | GSE237218 | GSM7597100 | HiSeq2000 | [24] |
| SRR25253938 | T cells | Memory | GSE237218 | GSM7597099 | HiSeq2000 | [24] |
| SRR25253939 | T cells | Memory | GSE237218 | GSM7597098 | HiSeq2000 | [24] |
| SRR25253940 | T cells | Memory | GSE237218 | GSM7597097 | HiSeq2000 | [24] |
| SRR25253941 | T cells | Memory | GSE237218 | GSM7597096 | HiSeq2000 | [24] |
| SRR25253942 | T cells | Memory | GSE237218 | GSM7597095 | HiSeq2000 | [24] |
| SRR25253943 | T cells | Memory | GSE237218 | GSM7597094 | HiSeq2000 | [24] |
| SRR25253944 | T cells | Diseased | GSE237218 | GSM7597093 | HiSeq2000 | [24] |
| SRR25253945 | T cells | Memory | GSE237218 | GSM7597092 | HiSeq2000 | [24] |
| SRR25253946 | T cells | Memory | GSE237218 | GSM7597091 | HiSeq2000 | [24] |
| SRR25253947 | T cells | Diseased | GSE237218 | GSM7597090 | HiSeq2000 | [24] |
| SRR25253948 | T cells | Memory | GSE237218 | GSM7597089 | HiSeq2000 | [24] |
| SRR25253949 | T cells | Memory | GSE237218 | GSM7597088 | HiSeq2000 | [24] |
| SRR25253950 | T cells | Memory | GSE237218 | GSM7597087 | HiSeq2000 | [24] |
| SRR25253951 | T cells | Memory | GSE237218 | GSM7597086 | HiSeq2000 | [24] |
| SRR25253952 | T cells | Memory | GSE237218 | GSM7597085 | HiSeq2000 | [24] |
| SRR25253953 | T cells | Diseased | GSE237218 | GSM7597084 | HiSeq2000 | [24] |
| SRR25253954 | T cells | Diseased | GSE237218 | GSM7597083 | HiSeq2000 | [24] |
| SRR25253955 | T cells | Diseased | GSE237218 | GSM7597082 | HiSeq2000 | [24] |
| SRR25253956 | T cells | Diseased | GSE237218 | GSM7597081 | HiSeq2000 | [24] |
| SRR25253957 | T cells | Memory | GSE237218 | GSM7597080 | HiSeq2000 | [24] |
| SRR25253958 | T cells | Memory | GSE237218 | GSM7597079 | HiSeq2000 | [24] |
| SRR25253959 | T cells | Memory | GSE237218 | GSM7597078 | HiSeq2000 | [24] |
| SRR25253960 | T cells | Regulatory | GSE237218 | GSM7597029 | HiSeq2000 | [24] |
| SRR25253961 | T cells | Regulatory | GSE237218 | GSM7597028 | HiSeq2000 | [24] |
| SRR25253962 | T cells | Diseased | GSE237218 | GSM7597027 | HiSeq2000 | [24] |
| SRR25253963 | T cells | Regulatory | GSE237218 | GSM7597026 | HiSeq2000 | [24] |
| SRR25253964 | T cells | Diseased | GSE237218 | GSM7597025 | HiSeq2000 | [24] |
| SRR25253965 | T cells | Regulatory | GSE237218 | GSM7597024 | HiSeq2000 | [24] |
| SRR25253966 | T cells | Regulatory | GSE237218 | GSM7597023 | HiSeq2000 | [24] |
| SRR25253967 | T cells | Diseased | GSE237218 | GSM7597022 | HiSeq2000 | [24] |
| SRR25253968 | T cells | Diseased | GSE237218 | GSM7597021 | HiSeq2000 | [24] |
| SRR25253969 | T cells | Diseased | GSE237218 | GSM7597020 | HiSeq2000 | [24] |
| SRR25253970 | T cells | Diseased | GSE237218 | GSM7597019 | HiSeq2000 | [24] |
| SRR25253971 | T cells | Regulatory | GSE237218 | GSM7597018 | HiSeq2000 | [24] |
| SRR25253972 | T cells | Regulatory | GSE237218 | GSM7597017 | HiSeq2000 | [24] |
| SRR25253973 | T cells | Regulatory | GSE237218 | GSM7597016 | HiSeq2000 | [24] |
| SRR25253974 | T cells | Regulatory | GSE237218 | GSM7597015 | HiSeq2000 | [24] |
| SRR25253975 | T cells | Regulatory | GSE237218 | GSM7597014 | HiSeq2000 | [24] |
| SRR25253976 | T cells | Regulatory | GSE237218 | GSM7597013 | HiSeq2000 | [24] |

| | | | | | | |
|---|---|---|---|---|---|---|
| SRR25253977 | T cells | Diseased | GSE237218 | GSM7597012 | HiSeq2000 | [24] |
| SRR25253978 | T cells | Regulatory | GSE237218 | GSM7597011 | HiSeq2000 | [24] |
| SRR25253979 | T cells | Regulatory | GSE237218 | GSM7597010 | HiSeq2000 | [24] |
| SRR25253980 | T cells | Diseased | GSE237218 | GSM7597009 | HiSeq2000 | [24] |
| SRR25253981 | T cells | Regulatory | GSE237218 | GSM7597008 | HiSeq2000 | [24] |
| SRR25253982 | T cells | Diseased | GSE237218 | GSM7597007 | HiSeq2000 | [24] |
| SRR25253983 | T cells | Diseased | GSE237218 | GSM7597006 | HiSeq2000 | [24] |
| SRR25253984 | B cells | Normal | GSE237218 | GSM7597293 | HiSeq2000 | [24] |
| SRR25253985 | B cells | Normal | GSE237218 | GSM7597292 | HiSeq2000 | [24] |
| SRR25253986 | B cells | Normal | GSE237218 | GSM7597291 | HiSeq2000 | [24] |
| SRR25253987 | B cells | Diseased | GSE237218 | GSM7597290 | HiSeq2000 | [24] |
| SRR25253988 | B cells | Diseased | GSE237218 | GSM7597289 | HiSeq2000 | [24] |
| SRR25253989 | B cells | Normal | GSE237218 | GSM7597288 | HiSeq2000 | [24] |
| SRR25253990 | B cells | Normal | GSE237218 | GSM7597287 | HiSeq2000 | [24] |
| SRR25253991 | B cells | Diseased | GSE237218 | GSM7597286 | HiSeq2000 | [24] |
| SRR25253992 | B cells | Normal | GSE237218 | GSM7597285 | HiSeq2000 | [24] |
| SRR25253993 | B cells | Diseased | GSE237218 | GSM7597284 | HiSeq2000 | [24] |
| SRR25253994 | B cells | Diseased | GSE237218 | GSM7597283 | HiSeq2000 | [24] |
| SRR25253995 | B cells | Diseased | GSE237218 | GSM7597282 | HiSeq2000 | [24] |
| SRR25253996 | B cells | Normal | GSE237218 | GSM7597281 | HiSeq2000 | [24] |
| SRR25253997 | T cells | Regulatory | GSE237218 | GSM7597280 | HiSeq2000 | [24] |
| SRR25253998 | B cells | Diseased | GSE237218 | GSM7597279 | HiSeq2000 | [24] |
| SRR25253999 | B cells | Diseased | GSE237218 | GSM7597278 | HiSeq2000 | [24] |
| SRR25254000 | B cells | Diseased | GSE237218 | GSM7597277 | HiSeq2000 | [24] |
| SRR25254001 | B cells | Normal | GSE237218 | GSM7597276 | HiSeq2000 | [24] |
| SRR25254002 | B cells | Diseased | GSE237218 | GSM7597275 | HiSeq2000 | [24] |
| SRR25254003 | B cells | Normal | GSE237218 | GSM7597274 | HiSeq2000 | [24] |
| SRR25254004 | B cells | Normal | GSE237218 | GSM7597273 | HiSeq2000 | [24] |
| SRR25254005 | B cells | Normal | GSE237218 | GSM7597272 | HiSeq2000 | [24] |
| SRR25254006 | B cells | Diseased | GSE237218 | GSM7597271 | HiSeq2000 | [24] |
| SRR25254007 | B cells | Diseased | GSE237218 | GSM7597270 | HiSeq2000 | [24] |
| SRR25254008 | T cells | Regulatory | GSE237218 | GSM7597005 | HiSeq2000 | [24] |
| SRR25254009 | T cells | Regulatory | GSE237218 | GSM7597004 | HiSeq2000 | [24] |
| SRR25254010 | T cells | Regulatory | GSE237218 | GSM7597003 | HiSeq2000 | [24] |
| SRR25254011 | T cells | Diseased | GSE237218 | GSM7597002 | HiSeq2000 | [24] |
| SRR25254012 | T cells | Regulatory | GSE237218 | GSM7597001 | HiSeq2000 | [24] |
| SRR25254013 | T cells | Diseased | GSE237218 | GSM7597000 | HiSeq2000 | [24] |
| SRR25254014 | T cells | Diseased | GSE237218 | GSM7596999 | HiSeq2000 | [24] |
| SRR25254015 | T cells | Regulatory | GSE237218 | GSM7596998 | HiSeq2000 | [24] |
| SRR25254016 | B cells | Diseased | GSE237218 | GSM7596997 | HiSeq2000 | [24] |
| SRR25254017 | B cells | Normal | GSE237218 | GSM7596996 | HiSeq2000 | [24] |
| SRR25254018 | B cells | Diseased | GSE237218 | GSM7596995 | HiSeq2000 | [24] |
| SRR25254019 | B cells | Diseased | GSE237218 | GSM7596994 | HiSeq2000 | [24] |
| SRR25254020 | B cells | Diseased | GSE237218 | GSM7596993 | HiSeq2000 | [24] |
| SRR25254021 | B cells | Diseased | GSE237218 | GSM7596992 | HiSeq2000 | [24] |
| SRR25254022 | B cells | Normal | GSE237218 | GSM7596991 | HiSeq2000 | [24] |

| | | | | | | |
|---|---|---|---|---|---|---|
| SRR25254023 | B cells | Normal | GSE237218 | GSM7596990 | HiSeq2000 | [24] |
| SRR25254024 | B cells | Normal | GSE237218 | GSM7596989 | HiSeq2000 | [24] |
| SRR25254025 | B cells | Normal | GSE237218 | GSM7596988 | HiSeq2000 | [24] |
| SRR25254026 | B cells | Diseased | GSE237218 | GSM7596987 | HiSeq2000 | [24] |
| SRR25254027 | B cells | Normal | GSE237218 | GSM7596986 | HiSeq2000 | [24] |
| SRR25254028 | B cells | Normal | GSE237218 | GSM7596985 | HiSeq2000 | [24] |
| SRR25254029 | B cells | Diseased | GSE237218 | GSM7596984 | HiSeq2000 | [24] |
| SRR25254030 | B cells | Normal | GSE237218 | GSM7596983 | HiSeq2000 | [24] |
| SRR25254031 | B cells | Normal | GSE237218 | GSM7596982 | HiSeq2000 | [24] |
| SRR25254032 | T cells | Regulatory | GSE237218 | GSM7597341 | HiSeq2000 | [24] |
| SRR5714385 | iPSC | Normal | GSE100223 | GSM2674875 | HiSeq2000 | [25] |
| SRR2453299 | Fibroblast | Normal | GSE73211 | GSM1888619 | HiSeq2000 | [26] |
| SRR2453300 | Fibroblast | Normal | GSE73211 | GSM1888619 | HiSeq2000 | [26] |
| SRR2453301 | iPSC | Normal | GSE73211 | GSM1888620 | HiSeq2000 | [26] |
| SRR2453302 | Fibroblast | Normal | GSE73211 | GSM1888621 | HiSeq2000 | [26] |
| SRR2453303 | iPSC | Normal | GSE73211 | GSM1888622 | HiSeq2000 | [26] |
| SRR2453304 | iPSC | Normal | GSE73211 | GSM1888622 | HiSeq2000 | [26] |
| SRR2453305 | Fibroblast | Normal | GSE73211 | GSM1888623 | HiSeq2000 | [26] |
| SRR2453306 | iPSC | Normal | GSE73211 | GSM1888624 | HiSeq2000 | [26] |
| SRR2453307 | Fibroblast | Normal | GSE73211 | GSM1888625 | HiSeq2000 | [26] |
| SRR2453308 | iPSC | Normal | GSE73211 | GSM1888626 | HiSeq2000 | [26] |
| SRR2453309 | Fibroblast | Normal | GSE73211 | GSM1888627 | HiSeq2000 | [26] |
| SRR2453310 | iPSC | Normal | GSE73211 | GSM1888628 | HiSeq2000 | [26] |
| SRR2453311 | Fibroblast | Normal | GSE73211 | GSM1888629 | HiSeq2000 | [26] |
| SRR2453312 | iPSC | Normal | GSE73211 | GSM1888630 | HiSeq2000 | [26] |
| SRR2453313 | iPSC | Normal | GSE73211 | GSM1888630 | HiSeq2000 | [26] |
| SRR2453314 | iPSC | Normal | GSE73211 | GSM1888630 | HiSeq2000 | [26] |
| SRR2453315 | ESC | Normal | GSE73211 | GSM1888631 | HiSeq2000 | [26] |
| SRR2453316 | ESC | Normal | GSE73211 | GSM1888632 | HiSeq2000 | [26] |
| SRR2453317 | ESC | Normal | GSE73211 | GSM1888633 | HiSeq2000 | [26] |
| SRR2453318 | Fibroblast | Normal | GSE73211 | GSM1888634 | HiSeq2000 | [26] |
| SRR2453319 | Fibroblast | Normal | GSE73211 | GSM1888635 | HiSeq2000 | [26] |
| SRR2453320 | Fibroblast | Normal | GSE73211 | GSM1888636 | HiSeq2000 | [26] |
| SRR2453321 | ESC | Normal | GSE73211 | GSM1888637 | HiSeq2000 | [26] |
| SRR2453322 | ESC | Normal | GSE73211 | GSM1888637 | HiSeq2000 | [26] |
| SRR2453323 | Fibroblast | Normal | GSE73211 | GSM1888638 | HiSeq2000 | [26] |
| SRR2453324 | ESC | Normal | GSE73211 | GSM1888639 | HiSeq2000 | [26] |
| SRR2453325 | ESC | Normal | GSE73211 | GSM1888639 | HiSeq2000 | [26] |
| SRR2453326 | ESC | Normal | GSE73211 | GSM1888640 | HiSeq2000 | [26] |
| SRR2453327 | ESC | Normal | GSE73211 | GSM1888640 | HiSeq2000 | [26] |
| SRR2453328 | ESC | Normal | GSE73211 | GSM1888641 | HiSeq2000 | [26] |
| SRR2453329 | ESC | Normal | GSE73211 | GSM1888642 | HiSeq2000 | [26] |
| SRR2453330 | ESC | Normal | GSE73211 | GSM1888642 | HiSeq2000 | [26] |
| SRR2453331 | ESC | Normal | GSE73211 | GSM1888643 | HiSeq2000 | [26] |
| SRR2453332 | Fibroblast | Normal | GSE73211 | GSM1888644 | HiSeq2000 | [26] |
| SRR2453333 | Fibroblast | Normal | GSE73211 | GSM1888644 | HiSeq2000 | [26] |

| | | | | | | |
|---|---|---|---|---|---|---|
| SRR2453334 | Fibroblast | Normal | GSE73211 | GSM1888645 | HiSeq2000 | [26] |
| SRR2453335 | Fibroblast | Normal | GSE73211 | GSM1888646 | HiSeq2000 | [26] |
| SRR2453336 | Fibroblast | Normal | GSE73211 | GSM1888647 | HiSeq2000 | [26] |
| SRR2453337 | ESC | Normal | GSE73211 | GSM1888648 | HiSeq2000 | [26] |
| SRR2453338 | ESC | Normal | GSE73211 | GSM1888648 | HiSeq2000 | [26] |
| SRR2453339 | ESC | Normal | GSE73211 | GSM1888649 | HiSeq2000 | [26] |
| SRR2453340 | ESC | Normal | GSE73211 | GSM1888650 | HiSeq2000 | [26] |
| SRR2453341 | Fibroblast | Normal | GSE73211 | GSM1888651 | HiSeq2000 | [26] |
| SRR2453342 | ESC | Normal | GSE73211 | GSM1888652 | HiSeq2000 | [26] |
| SRR2453343 | ESC | Normal | GSE73211 | GSM1888653 | HiSeq2000 | [26] |
| SRR2453344 | iPSC | Normal | GSE73211 | GSM1888654 | HiSeq2000 | [26] |
| SRR2453345 | iPSC | Normal | GSE73211 | GSM1888655 | HiSeq2000 | [26] |
| SRR2453346 | ESC | Normal | GSE73211 | GSM1888656 | HiSeq2000 | [26] |
| SRR2453347 | iPSC | Normal | GSE73211 | GSM1888657 | HiSeq2000 | [26] |
| SRR2453348 | iPSC | Normal | GSE73211 | GSM1888658 | HiSeq2000 | [26] |
| SRR2453349 | iPSC | Normal | GSE73211 | GSM1888659 | HiSeq2000 | [26] |
| SRR2453350 | iPSC | Normal | GSE73211 | GSM1888660 | HiSeq2000 | [26] |
| SRR2453351 | ESC | Normal | GSE73211 | GSM1888661 | HiSeq2000 | [26] |
| SRR2453352 | iPSC | Normal | GSE73211 | GSM1888662 | HiSeq2000 | [26] |
| SRR2453353 | iPSC | Normal | GSE73211 | GSM1888663 | HiSeq2000 | [26] |
| SRR2453354 | ESC | Normal | GSE73211 | GSM1888664 | HiSeq2000 | [26] |
| SRR2453355 | ESC | Normal | GSE73211 | GSM1888665 | HiSeq2000 | [26] |
| SRR2453356 | ESC | Normal | GSE73211 | GSM1888666 | HiSeq2000 | [26] |
| SRR6355965 | iPSC | Normal | GSE107654 | GSM2392700 | HiSeq2500 | [27] |
| SRR6355966 | iPSC | Normal | GSE107654 | GSM2392701 | HiSeq2500 | [27] |
| SRR6355967 | iPSC | Normal | GSE107654 | GSM2392702 | HiSeq2500 | [27] |
| SRR6355968 | iPSC | Normal | GSE107654 | GSM2392703 | HiSeq2500 | [27] |
| SRR6355969 | iPSC | Normal | GSE107654 | GSM2392704 | HiSeq2500 | [27] |
| SRR6355970 | iPSC | Normal | GSE107654 | GSM2392705 | HiSeq2500 | [27] |
| SRR6355971 | iPSC | Normal | GSE107654 | GSM2392706 | HiSeq2500 | [27] |
| SRR6355972 | iPSC | Normal | GSE107654 | GSM2392707 | HiSeq2500 | [27] |
| SRR6355973 | iPSC | Normal | GSE107654 | GSM2392708 | HiSeq2500 | [27] |
| SRR6355974 | iPSC | Normal | GSE107654 | GSM2392709 | HiSeq2500 | [27] |
| SRR6355975 | iPSC | Normal | GSE107654 | GSM2392710 | HiSeq2500 | [27] |
| SRR6355976 | iPSC | Normal | GSE107654 | GSM2392711 | HiSeq2500 | [27] |
| SRR6355977 | iPSC | Normal | GSE107654 | GSM2392712 | HiSeq2500 | [27] |
| SRR6355978 | iPSC | Normal | GSE107654 | GSM2392713 | HiSeq2500 | [27] |
| SRR6355979 | iPSC | Normal | GSE107654 | GSM2392714 | HiSeq2500 | [27] |
| SRR6355980 | iPSC | Normal | GSE107654 | GSM2392715 | HiSeq2500 | [27] |
| SRR6355981 | iPSC | Normal | GSE107654 | GSM2392716 | HiSeq2500 | [27] |
| SRR6355982 | iPSC | Normal | GSE107654 | GSM2392717 | HiSeq2500 | [27] |
| SRR6355983 | iPSC | Normal | GSE107654 | GSM2392718 | HiSeq2500 | [27] |
| SRR6355984 | iPSC | Normal | GSE107654 | GSM2392719 | HiSeq2500 | [27] |
| SRR6355985 | iPSC | Normal | GSE107654 | GSM2392720 | HiSeq2500 | [27] |
| SRR6355986 | iPSC | Normal | GSE107654 | GSM2392721 | HiSeq2500 | [27] |
| SRR6355987 | iPSC | Normal | GSE107654 | GSM2392722 | HiSeq2500 | [27] |

| | | | | | | |
|---|---|---|---|---|---|---|
| SRR6355988 | iPSC | Normal | GSE107654 | GSM2392723 | HiSeq2500 | [27] |
| SRR6355989 | iPSC | Normal | GSE107654 | GSM2392724 | HiSeq2500 | [27] |
| SRR6355990 | iPSC | Normal | GSE107654 | GSM2392725 | HiSeq2500 | [27] |
| SRR6355991 | iPSC | Normal | GSE107654 | GSM2392726 | HiSeq2500 | [27] |
| SRR6355992 | iPSC | Normal | GSE107654 | GSM2392727 | HiSeq2500 | [27] |
| SRR6355993 | iPSC | Normal | GSE107654 | GSM2392728 | HiSeq2500 | [27] |
| SRR6355994 | iPSC | Normal | GSE107654 | GSM2392729 | HiSeq2500 | [27] |
| SRR6355995 | iPSC | Normal | GSE107654 | GSM2392730 | HiSeq2500 | [27] |
| SRR6355996 | iPSC | Normal | GSE107654 | GSM2392731 | HiSeq2500 | [27] |
| SRR6355997 | iPSC | Normal | GSE107654 | GSM2392732 | HiSeq2500 | [27] |
| SRR6355998 | iPSC | Normal | GSE107654 | GSM2392733 | HiSeq2500 | [27] |
| SRR6355999 | iPSC | Normal | GSE107654 | GSM2392734 | HiSeq2500 | [27] |
| SRR6356000 | iPSC | Normal | GSE107654 | GSM2392735 | HiSeq2500 | [27] |
| SRR6356001 | iPSC | Normal | GSE107654 | GSM2392736 | HiSeq2500 | [27] |
| SRR6356002 | iPSC | Normal | GSE107654 | GSM2392737 | HiSeq2500 | [27] |

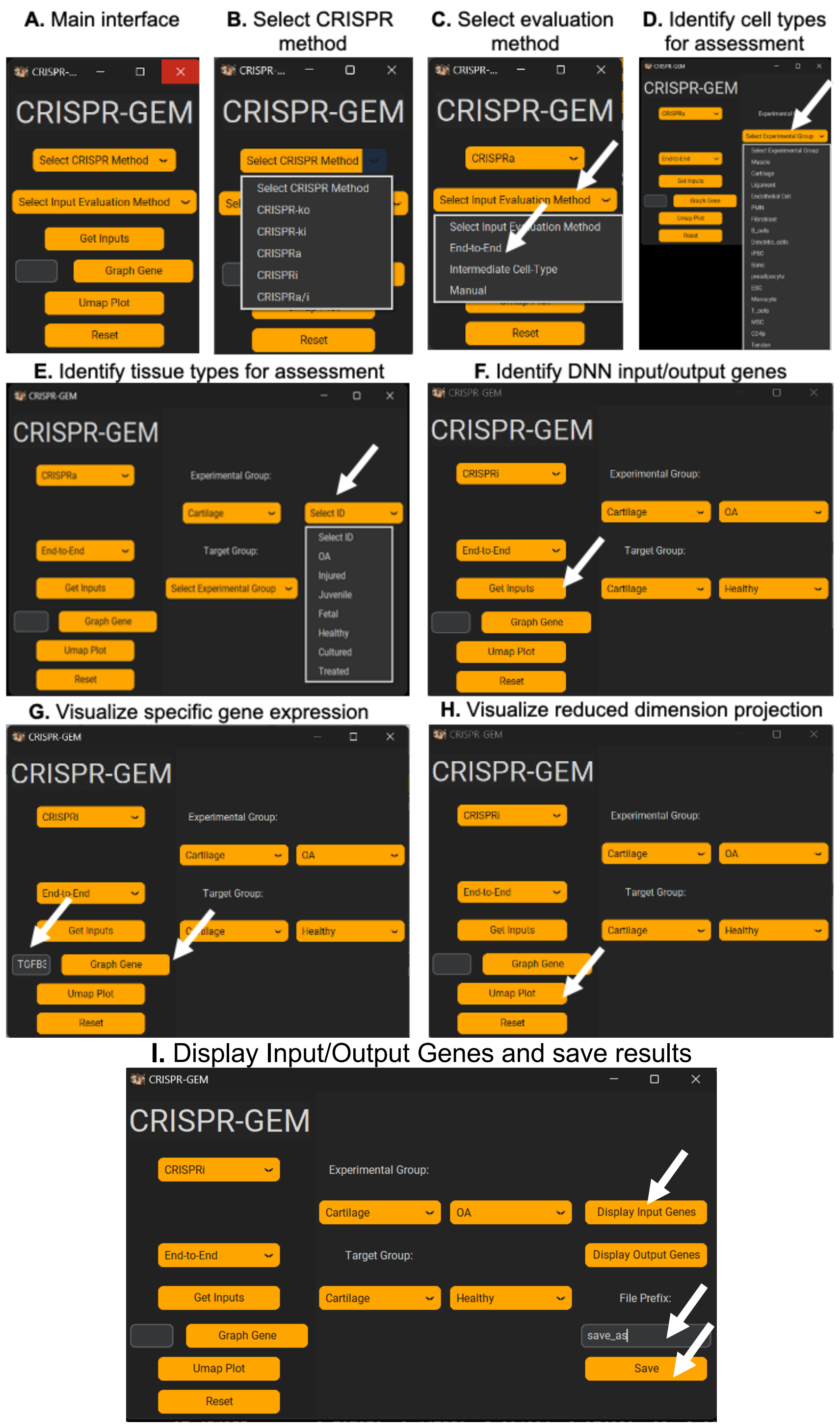

**Supp. Fig. 1.** CRISPR-GEM graphical user interface (GUI). **(A)** The main interface displayed upon opening the application. **(B)** The user must first select the desired CRISPR method (CRISPRko, CRISPRki, CRISPRa, or CRISPRi). **(C)** The user may then determine if input/output genes will be selected using just an experimental and target group (End-to-End), with an additional intermediate group (Intermediate Cell-type), or by manually inputting genes (Manual). Next, the **(D)** cell-type and **(E)** description for each group are specified and the input/output genes are determined through differential expression analysis by pressing the “Get Inputs” button **(F)**. Further analysis of the groups may be performed by graphing the expression of specific genes by typing the gene symbol in the text box and pressing the “Graph Gene” button **(G)** or by plotting the data in a reduced dimension UMAP projection **(H)**. **(I)** Final input and output genes may be viewed in an interactive data frame by clicking the “Display Input Genes” or “Display Output Genes”. Model results can be saved by typing a descriptor for the analysis in the text box and pressing the “Save” button.

**Differential Gene Expression**

**Supplemental Table 2:** Datasets used in differential expression analysis and gene evaluation for CRISPRa induced Treg differentiation.

| Tregs | iPSCs |
|---|---|
| SRR25253975 | SRR2453349 |
| SRR25253997 | SRR2453353 |
| SRR25253703 | SRR2453352 |
| SRR25253642 | SRR2453350 |
| SRR25253780 | SRR2453345 |
| SRR25254015 | SRR2453308 |
| SRR25253654 | SRR2453304 |
| SRR25253961 | SRR5714385 |
| SRR25253922 | SRR2453314 |
| SRR25254012 | SRR2453310 |
| SRR25253684 | SRR2453306 |
| SRR25254010 | SRR2453313 |
| SRR25253672 | SRR2453344 |
| SRR25253978 | SRR2453347 |
| SRR25253976 | SRR2453303 |
| SRR25253664 | SRR2453312 |
| SRR25253963 | SRR2453348 |
| SRR25253974 | SRR2453301 |

**Supplemental Table 3:** Datasets used in differential expression analysis and gene evaluation for CRISPRa induced chondrogenesis.

| MSC | Fetal Cartilage | Juvenile Cartilage |
|---|---|---|
| SRR22560355 | SRR3536062 | SRR6337330 |
| SRR21009233 | SRR3536063 | SRR6337329 |
| SRR13018695 | SRR3536064 | SRR6337331 |
| SRR21009234 | SRR3536065 | SRR6337332 |

**Supplemental table 4:** Datasets used in differential expression analysis and gene evaluation for CRISPRi based reversal of osteoarthritis.

| Healthy Cartilage | OA Cartilage |
|---|---|
| SRR7107487 | SRR7107495 |
| SRR7107486 | SRR7107496 |
| SRR7107483 | SRR7107491 |
| SRR7107484 | SRR7107494 |
| SRR7107488 | SRR7107493 |
| SRR7107481 | SRR7107498 |

**CRISPR-GEM Results**

**Supplemental Table 11.** Top 50 genes identified by CRISPR-GEM for CRISPRa induced Treg differentiation.

| Gene | 1 | 2 | 3 | 4 | 5 | Geometric Mean |
|---|---|---|---|---|---|---|
| **SRGN** | 0.49491 | 0.46499 | 0.5116 | 0.4965 | 0.48416 | 0.490187 |
| **LTB** | 0.50164 | 0.4638 | 0.52063 | 0.51146 | 0.50688 | 0.50049 |
| **ARHGDIB** | 0.50336 | 0.47801 | 0.52359 | 0.50888 | 0.49164 | 0.500858 |
| **SQOR** | 0.49737 | 0.48303 | 0.52659 | 0.50447 | 0.49705 | 0.5015 |
| **LY96** | 0.50504 | 0.47558 | 0.52592 | 0.50885 | 0.49954 | 0.502718 |
| **LSP1** | 0.51017 | 0.47167 | 0.52199 | 0.50433 | 0.51214 | 0.50376 |
| **TRADD** | 0.49861 | 0.47531 | 0.52602 | 0.5162 | 0.50526 | 0.503983 |
| **CD48** | 0.51133 | 0.47429 | 0.52311 | 0.51291 | 0.50076 | 0.504197 |
| **SELL** | 0.51435 | 0.47057 | 0.53016 | 0.49609 | 0.5157 | 0.504951 |
| **CYTIP** | 0.48983 | 0.47175 | 0.53801 | 0.52353 | 0.51044 | 0.506156 |
| **CASP4** | 0.51207 | 0.4813 | 0.52428 | 0.51387 | 0.50081 | 0.506254 |
| **CTSS** | 0.49935 | 0.48069 | 0.53198 | 0.51528 | 0.50621 | 0.506415 |
| **KLF2** | 0.50923 | 0.47308 | 0.53673 | 0.51135 | 0.50763 | 0.507194 |
| **TSC22D3** | 0.51559 | 0.48556 | 0.53041 | 0.51033 | 0.49571 | 0.507279 |
| **S1PR4** | 0.50186 | 0.47861 | 0.52889 | 0.52052 | 0.50884 | 0.507445 |
| **CD69** | 0.52156 | 0.47364 | 0.52567 | 0.5134 | 0.50529 | 0.507564 |
| **CAT** | 0.49739 | 0.48644 | 0.53251 | 0.5095 | 0.51724 | 0.508368 |
| **ITGB2** | 0.50581 | 0.47293 | 0.53359 | 0.51754 | 0.51606 | 0.508778 |
| **ALOX5AP** | 0.50411 | 0.47712 | 0.5342 | 0.51508 | 0.51559 | 0.508869 |
| **PIK3IP1** | 0.51337 | 0.47171 | 0.52673 | 0.52266 | 0.51185 | 0.508875 |
| **CD52** | 0.51449 | 0.47691 | 0.52866 | 0.52013 | 0.50757 | 0.509237 |
| **RASSF5** | 0.50896 | 0.48162 | 0.53045 | 0.51584 | 0.5136 | 0.509839 |
| **FCMR** | 0.50729 | 0.48628 | 0.52824 | 0.51544 | 0.51312 | 0.509888 |
| **GMFG** | 0.5022 | 0.48116 | 0.53236 | 0.51917 | 0.51708 | 0.51009 |
| **LAPTM5** | 0.50828 | 0.48877 | 0.54082 | 0.50712 | 0.50725 | 0.510174 |
| **PTPRCAP** | 0.51086 | 0.48109 | 0.52732 | 0.52594 | 0.5076 | 0.510283 |
| **LYZ** | 0.51418 | 0.48201 | 0.53449 | 0.51357 | 0.5095 | 0.51047 |
| **CEBPD** | 0.52648 | 0.47899 | 0.52217 | 0.52129 | 0.51342 | 0.512169 |
| **HCLS1** | 0.51572 | 0.47891 | 0.53098 | 0.52216 | 0.51486 | 0.512209 |
| **VSIR** | 0.51388 | 0.47742 | 0.53308 | 0.526 | 0.51338 | 0.512386 |
| **TSPO** | 0.51275 | 0.48556 | 0.53524 | 0.51556 | 0.515 | 0.512574 |
| **SP100** | 0.51955 | 0.48696 | 0.52872 | 0.51606 | 0.51507 | 0.513076 |
| **HLA-F** | 0.51861 | 0.48645 | 0.53108 | 0.51607 | 0.51497 | 0.513223 |
| **C1orf162** | 0.52204 | 0.47515 | 0.53531 | 0.5231 | 0.51264 | 0.513226 |
| **BCL2A1** | 0.5105 | 0.49261 | 0.53372 | 0.51359 | 0.52116 | 0.514138 |
| **TYMP** | 0.51297 | 0.49369 | 0.53147 | 0.5222 | 0.51162 | 0.514236 |
| **NAAA** | 0.51092 | 0.47341 | 0.53805 | 0.52891 | 0.52285 | 0.514321 |
| **CD44** | 0.51376 | 0.48324 | 0.53613 | 0.52333 | 0.51765 | 0.514517 |
| **CARD16** | 0.50592 | 0.48544 | 0.54096 | 0.52024 | 0.52282 | 0.51474 |
| **ARHGAP15** | 0.50692 | 0.49458 | 0.53062 | 0.52504 | 0.51766 | 0.5148 |
| **IL2RG** | 0.51163 | 0.48957 | 0.54312 | 0.52355 | 0.50775 | 0.514821 |

| | | | | | | |
|---|---|---|---|---|---|---|
| **RPS27P4** | 0.51986 | 0.47968 | 0.53288 | 0.52868 | 0.51502 | 0.514867 |
| **SELPLG** | 0.51463 | 0.47978 | 0.53826 | 0.52575 | 0.51831 | 0.514967 |
| **ICAM3** | 0.51541 | 0.48572 | 0.53393 | 0.52232 | 0.51884 | 0.514988 |
| **LY86** | 0.51706 | 0.47444 | 0.53461 | 0.52744 | 0.52442 | 0.515138 |
| **GBP2** | 0.51679 | 0.48478 | 0.53546 | 0.52764 | 0.51295 | 0.515226 |
| **TBC1D10C** | 0.51216 | 0.48439 | 0.5257 | 0.53572 | 0.51979 | 0.515254 |
| **TNFRSF14** | 0.51742 | 0.48141 | 0.5465 | 0.52324 | 0.51066 | 0.515413 |
| **LPXN** | 0.51432 | 0.49308 | 0.53537 | 0.51234 | 0.52315 | 0.51546 |
| **ANXA2R** | 0.51474 | 0.4872 | 0.53872 | 0.514 | | |

**Supplemental Table 12.** Top 50 genes identified by CRISPR-GEM for CRISPRa induced chondrogenesis.

| Gene | 1 | 2 | 3 | 4 | 5 | Geometric Mean |
|---|---|---|---|---|---|---|
| **HSPA6** | 0.27006 | 0.26964 | 0.28255 | 0.27084 | 0.27172 | 0.272918 |
| **PRG4** | 0.27048 | 0.2675 | 0.27858 | 0.27952 | 0.2707 | 0.273313 |
| **UCMA** | 0.2795 | 0.26597 | 0.27955 | 0.27829 | 0.2706 | 0.274726 |
| **SERPINA1** | 0.27307 | 0.26848 | 0.28399 | 0.27812 | 0.27235 | 0.275151 |
| **CNMD** | 0.27397 | 0.26806 | 0.28511 | 0.27986 | 0.26981 | 0.275289 |
| **RPL35AP2** | 0.27828 | 0.27511 | 0.28281 | 0.27633 | 0.26483 | 0.275407 |
| **IL1RL1** | 0.27586 | 0.27088 | 0.27906 | 0.2777 | 0.27393 | 0.27547 |
| **MMP10** | 0.27167 | 0.26826 | 0.28116 | 0.28177 | 0.27622 | 0.275766 |
| **FRZB** | 0.27119 | 0.26728 | 0.29077 | 0.2784 | 0.27201 | 0.275807 |
| **KIAA1328** | 0.27768 | 0.27252 | 0.28875 | 0.27279 | 0.27044 | 0.276359 |
| **SINHCAFP3** | 0.2755 | 0.27492 | 0.28721 | 0.28175 | 0.26822 | 0.277445 |
| **NMNAT3** | 0.27901 | 0.27871 | 0.27898 | 0.27805 | 0.27358 | 0.277657 |
| **MAB21L3** | 0.2773 | 0.27527 | 0.28489 | 0.2826 | 0.27201 | 0.278372 |
| **GPC3** | 0.27526 | 0.27332 | 0.28223 | 0.28631 | 0.2769 | 0.278762 |
| **PEG3** | 0.27428 | 0.27528 | 0.28715 | 0.28041 | 0.27719 | 0.278823 |
| **COL9A1** | 0.28251 | 0.27719 | 0.2753 | 0.28154 | 0.27771 | 0.278835 |
| **OR4F17** | 0.28436 | 0.26938 | 0.28611 | 0.27847 | 0.2765 | 0.278898 |
| **H1-4** | 0.2832 | 0.27175 | 0.28647 | 0.28134 | 0.27278 | 0.279048 |
| **ATP6V0A4** | 0.281 | 0.2717 | 0.28168 | 0.28366 | 0.27955 | 0.279486 |
| **ZNF385B** | 0.28028 | 0.27289 | 0.28619 | 0.28278 | 0.27614 | 0.279615 |
| **DLK1** | 0.28419 | 0.27051 | 0.29192 | 0.27954 | 0.27546 | 0.280226 |
| **CCL20** | 0.27542 | 0.27822 | 0.27737 | 0.2861 | 0.28446 | 0.280281 |
| **NACA3P** | 0.28027 | 0.2766 | 0.28277 | 0.28041 | 0.28223 | 0.280448 |
| **ALKAL2** | 0.28046 | 0.2775 | 0.28837 | 0.28492 | 0.2716 | 0.280509 |
| **LPAR4** | 0.28133 | 0.27545 | 0.28476 | 0.28607 | 0.2785 | 0.281195 |
| **BMP2** | 0.27634 | 0.27937 | 0.29135 | 0.28056 | 0.27881 | 0.281237 |
| **PELI1** | 0.28134 | 0.28037 | 0.28267 | 0.28573 | 0.27705 | 0.281417 |
| **NOS2** | 0.28309 | 0.27034 | 0.28922 | 0.28662 | 0.27844 | 0.281462 |
| **BIRC3** | 0.28367 | 0.27615 | 0.28327 | 0.28043 | 0.28541 | 0.281768 |
| **SLC7A2** | 0.28622 | 0.27469 | 0.28872 | 0.28362 | 0.2764 | 0.281877 |
| **GASK1A** | 0.28061 | 0.27378 | 0.29202 | 0.28524 | 0.28002 | 0.282269 |
| **CHRDL2** | 0.28243 | 0.28115 | 0.29189 | 0.28003 | 0.27697 | 0.282448 |
| **ITM2A** | 0.28777 | 0.27137 | 0.28968 | 0.28624 | 0.2784 | 0.282607 |

| | | | | | | |
|---|---|---|---|---|---|---|
| **EIF2S2P4** | 0.28099 | 0.28248 | 0.28692 | 0.2861 | 0.27743 | 0.282761 |
| **REL** | 0.28845 | 0.27715 | 0.28755 | 0.28558 | 0.2756 | 0.282813 |
| **SOX8** | 0.28769 | 0.27837 | 0.28359 | 0.28693 | 0.27803 | 0.282893 |
| **RPL21P119** | 0.28343 | 0.27878 | 0.28649 | 0.28433 | 0.2816 | 0.282913 |
| **H1-3** | 0.29032 | 0.27628 | 0.28678 | 0.28826 | 0.27362 | 0.28297 |
| **MDM4** | 0.28104 | 0.27317 | 0.29214 | 0.28874 | 0.28039 | 0.283018 |
| **ITIH6** | 0.2826 | 0.27877 | 0.29025 | 0.2817 | 0.28219 | 0.283078 |
| **PTGS2** | 0.28706 | 0.27683 | 0.2847 | 0.28726 | 0.28013 | 0.283164 |
| **MYO5B** | 0.2872 | 0.27498 | 0.28642 | 0.28708 | 0.28139 | 0.283374 |
| **SEMA6D** | 0.28207 | 0.27708 | 0.28475 | 0.28415 | 0.289 | 0.283382 |
| **B3GNT7** | 0.28338 | 0.2806 | 0.29204 | 0.28547 | 0.27593 | 0.283432 |
| **TLR2** | 0.29019 | 0.27748 | 0.28366 | 0.28768 | 0.27854 | 0.283467 |
| **NFIA** | 0.28068 | 0.28146 | 0.2897 | 0.28429 | 0.28166 | 0.283538 |
| **JMJD1C** | 0.28281 | 0.27967 | 0.28847 | 0.28691 | 0.28025 | 0.2836 |
| **RPL21P16** | 0.28115 | 0.28 | 0.28226 | 0.289 | 0.28582 | 0.283627 |
| **MATN1** | 0.28571 | 0.28191 | 0.2892 | 0.28171 | 0.28018 | 0.283724 |
| **CXCR4** | 0.27668 | 0.28627 | 0.28853 | 0.28455 | 0.28282 | 0.28374 |

**Supplemental Table 12.** Top 50 genes identified by CRISPR-GEM for CRISPRi based reversal of osteoarthritis.

| Gene | 1 | 2 | 3 | 4 | 5 | Geometric Mean |
|---|---|---|---|---|---|---|
| **FAP** | 0.336541 | 0.326209 | 0.325342 | 0.335197 | 0.305524 | 0.325569 |
| **VCAM1** | 0.342457 | 0.334714 | 0.338503 | 0.342657 | 0.313462 | 0.334178 |
| **IGFL4** | 0.34283 | 0.335216 | 0.339784 | 0.340851 | 0.320926 | 0.335827 |
| **COL1A1** | 0.346349 | 0.340732 | 0.339482 | 0.343549 | 0.31721 | 0.3373 |
| **DIO2** | 0.347713 | 0.339386 | 0.342752 | 0.347416 | 0.323932 | 0.340126 |
| **COL1A2** | 0.343549 | 0.34491 | 0.337815 | 0.349549 | 0.328983 | 0.340888 |
| **TNFAIP6** | 0.346196 | 0.34122 | 0.343334 | 0.349481 | 0.324854 | 0.340908 |
| **SPP1** | 0.349715 | 0.344514 | 0.345019 | 0.350571 | 0.32769 | 0.3434 |
| **TGFBI** | 0.353901 | 0.343188 | 0.346579 | 0.351392 | 0.329958 | 0.3449 |
| **THY1** | 0.349956 | 0.347793 | 0.34863 | 0.352649 | 0.326322 | 0.344935 |
| **EDEM3** | 0.351639 | 0.347303 | 0.347062 | 0.353084 | 0.328089 | 0.345316 |
| **HBB** | 0.353298 | 0.345604 | 0.350006 | 0.353773 | 0.330599 | 0.346549 |
| **POSTN** | 0.355783 | 0.348408 | 0.350051 | 0.35167 | 0.32989 | 0.347042 |
| **PXYLP1** | 0.356875 | 0.347563 | 0.34848 | 0.354692 | 0.330549 | 0.347507 |
| **TREM1** | 0.354892 | 0.350274 | 0.34955 | 0.355297 | 0.329456 | 0.347761 |
| **AMTN** | 0.357332 | 0.347383 | 0.351341 | 0.353804 | 0.330505 | 0.347945 |

| | | | | | | |
|---|---|---|---|---|---|---|
| **CFI** | 0.357291 | 0.349552 | 0.349262 | 0.354607 | 0.333096 | 0.348659 |
| **BEND6** | 0.359677 | 0.351568 | 0.350936 | 0.354818 | 0.330899 | 0.349438 |
| **C14orf119** | 0.359106 | 0.35089 | 0.35056 | 0.354988 | 0.334331 | 0.349872 |
| **THBS2** | 0.357756 | 0.350038 | 0.350466 | 0.356928 | 0.334797 | 0.349898 |
| **SPPL2A** | 0.356029 | 0.352583 | 0.352184 | 0.355093 | 0.335328 | 0.350159 |
| **SHC4** | 0.356058 | 0.352306 | 0.353782 | 0.357081 | 0.333845 | 0.350508 |
| **LRRC15** | 0.35968 | 0.351083 | 0.352454 | 0.356964 | 0.333495 | 0.350613 |
| **CHI3L2** | 0.361451 | 0.349521 | 0.353572 | 0.351904 | 0.337455 | 0.350694 |
| **MMP13** | 0.359181 | 0.35048 | 0.353837 | 0.357949 | 0.335715 | 0.351329 |
| **DKK1** | 0.359166 | 0.352449 | 0.352611 | 0.359664 | 0.333617 | 0.351371 |
| **NOVA1** | 0.358038 | 0.352551 | 0.355542 | 0.356953 | 0.334612 | 0.35143 |
| **ZNF100** | 0.361191 | 0.35375 | 0.353616 | 0.358198 | 0.333907 | 0.352 |
| **ZNF93** | 0.359971 | 0.354036 | 0.354397 | 0.358834 | 0.333922 | 0.352102 |
| **SCRN3** | 0.36078 | 0.353144 | 0.353341 | 0.359349 | 0.335745 | 0.352357 |
| **EVI2A** | 0.361269 | 0.351797 | 0.354432 | 0.358936 | 0.336792 | 0.352539 |
| **IGFBP3** | 0.361193 | 0.352893 | 0.354284 | 0.360261 | 0.334728 | 0.35254 |
| **ZNF761** | 0.360078 | 0.353051 | 0.356427 | 0.359484 | 0.334494 | 0.352578 |
| **CYP46A1** | 0.360264 | 0.355918 | 0.353109 | 0.360674 | 0.335756 | 0.353024 |
| **STEAP4** | 0.360492 | 0.354258 | 0.357754 | 0.359793 | 0.335031 | 0.353336 |
| **ADAM12** | 0.36048 | 0.355315 | 0.355865 | 0.360877 | 0.335314 | 0.353442 |
| **SPOCK1** | 0.360641 | 0.35759 | 0.353423 | 0.361144 | 0.336426 | 0.353724 |
| **CDH11** | 0.360737 | 0.355366 | 0.356288 | 0.360239 | 0.336688 | 0.353751 |
| **PREX2** | 0.362528 | 0.354722 | 0.356994 | 0.36195 | 0.336682 | 0.354447 |
| **ZNF845** | 0.362405 | 0.355621 | 0.356979 | 0.360779 | 0.337304 | 0.354501 |
| **RRM2** | 0.362523 | 0.355591 | 0.356954 | 0.361446 | 0.336732 | 0.354524 |
| **ERAP2** | 0.362269 | 0.356018 | 0.356586 | 0.360407 | 0.339585 | 0.35488 |
| **TMEM184C** | 0.362685 | 0.355596 | 0.355514 | 0.361936 | 0.339253 | 0.354895 |
| **GEN1** | 0.363363 | 0.356142 | 0.357348 | 0.361369 | 0.338679 | 0.35527 |
| **BLID** | 0.363975 | 0.354912 | 0.356348 | 0.362774 | 0.338917 | 0.35527 |
| **TTC9** | 0.36321 | 0.355339 | 0.358015 | 0.362631 | 0.33808 | 0.355334 |
| **SLC38A7** | 0.363491 | 0.354552 | 0.356802 | 0.364628 | 0.338085 | 0.355382 |

| | | | | | | |
|---|---|---|---|---|---|---|
| **PUS7L** | 0.363405 | 0.35651 | 0.358246 | 0.360576 | 0.338983 | 0.355438 |
| **ZKSCAN4** | 0.363504 | 0.357374 | 0.357247 | 0.361908 | 0.338184 | 0.355525 |
| **ALPL** | 0.363055 | 0.356429 | 0.358011 | 0.362395 | 0.33889 | 0.355645 |
